# Parallel basal ganglia and frontal cortical outputs differentially encode context-dependent evaluation and categorical commitment during choice

**DOI:** 10.64898/2026.06.15.732482

**Authors:** Atsushi Yoshida, Richard J. Krauzlis, Okihide Hikosaka

## Abstract

Adaptive choice requires transforming the evaluation of available options into commitment to a specific action but understanding how this transformation is implemented across neural circuits remains a central challenge. Here we recorded well-isolated neurons in the substantia nigra pars reticulata (SNr) and frontal eye field (FEF), which send parallel projections to the superior colliculus for driving the eye movement choice, while monkeys performed a sequential-offer choice task designed to partially dissociate scene-defined ordinal rank from the categorical commitment. Before target onset, SNr displayed stronger scene-related modulation than FEF. During target evaluation, SNr activity showed ordered modulation across behavioral outcomes dominated by ordinal rank, whereas FEF activity categorically separated acceptance from rejection and strongly encoded target direction. Behavioral model decomposition revealed that ordinal rank alone best explained SNr activity, outperforming both reward magnitude and even a composite acceptability measure that incorporated rank together with reward, scene context, and waiting cost, whereas FEF activity was best explained by categorical commitment. This dissociation was consistent across multivariable modeling, single-neuron response patterns, and all three monkeys. Together, these findings support a division of labor in which context-dependent evaluation and categorical commitment are distributed across parallel basal ganglia and frontal cortical output pathways to efficiently guide voluntary choices.

## Introduction

Adaptive choice requires the brain to transform graded evaluation of available options (Gold & Shadlen, 2007) into the categorical commitment to a specific action (Thura & Cisek, 2014). Where and how this transformation is implemented in specific neural circuits remains a central question in systems neuroscience (Chen & Stuphorn, 2015; Zhang et al., 2021).

The superior colliculus (SC) provides a key node for examining this transformation (Hikosaka et al., 2000; Hanes & Wurtz, 2001). It receives convergent input from two major pathways that are well-characterized both anatomically and physiologically: inhibitory basal ganglia output from the substantia nigra pars reticulata (SNr) (Jayaraman et al., 1977; Hikosaka & Wurtz, 1983; May & Hall, 1984; Liu & Basso, 2008) and excitatory frontal cortical input from the frontal eye field (FEF) (Künzle et al., 1976; Helminski & Segraves, 2003; Segraves & Goldberg, 1987; Stanton et al., 1988; Sommer & Wurtz, 2000). Because these pathways operate in parallel and converge immediately upstream of movement execution, they provide a defined circuit in which evaluation-related and commitment-related signals can be compared directly (Hanes & Wurtz, 2001; Hikosaka & Wurtz, 1983). SC neurons are also modulated by reward (Ikeda & Hikosaka, 2003, 2007), selective attention (Herman et al., 2018), and behavioral context (Basso & Wurtz, 1998; Everling et al., 1999; Horwitz & Newsome, 1999; Horwitz et al., 2004), suggesting that this convergence site can integrate evaluative and goal-related information rather than simply relay motor commands.

Prior work suggests that SNr and FEF make functionally distinct contributions to this process. The SNr exerts tonic inhibition over the SC, and classic models emphasize disinhibition as the core mechanism by which the basal ganglia gate actions (Hikosaka & Wurtz, 1983; Chevalier & Deniau, 1990; Mink, 1996). Within the broader cortico-basal ganglia architecture, the SNr is also positioned to integrate contextual and reward-related information before it reaches downstream motor structures (Handel & Glimcher, 2000; Sato & Hikosaka, 2002; Yasuda et al., 2012, Yasuda & Hikosaka, 2015). Consistent with this view, reward-related and contextual modulation can be more prominent in the caudate nucleus and related basal ganglia circuits than in FEF, particularly for reward-context signals (Ding & Hikosaka, 2006; Fan et al., 2020; Doi et al., 2020). The FEF, by contrast, provides topographically organized excitatory drive to the SC (Segraves & Goldberg, 1987; Stanton et al., 1988; Sommer & Wurtz, 2000) and conveys signals associated with target selection and movement initiation (Schall & Hanes, 1993; Hanes & Schall, 1996). These observations raise the possibility that SNr preferentially conveys contextual and evaluative signals, whereas FEF preferentially conveys categorical commitment and directional signals, but this has not been directly tested because evaluation and action are not readily dissociable in most reward-guided tasks (Cisek & Kalaska, 2010).

In such tasks, better options are accepted and worse options are rejected, with rejection typically expressed as the absence of movement (Hanes & Schall, 1996; Tremblay & Schultz, 1999). Under these conditions, neural correlates of option evaluation are difficult to separate from those of motor preparation or suppression (Chen & Stuphorn, 2015). We addressed this confound using a sequential-offer choice task in which a background scene defines the local choice set and thereby the scene-defined ordinal value ranking of each offered object (Yoshida & Hikosaka, 2024, 2025a, 2025b; Yoshida et al., 2026). Crucially, monkeys could reject a lower-ranked option through an active return saccade, and in scenes with small reward differences they sometimes accepted the lower-ranked option. This design allowed us to dissociate scene-defined rank from categorical commitment and to test whether these parallel pathways carry distinct aspects of the evaluation-to-commitment transformation. Importantly, evaluation itself can take multiple forms that are difficult to distinguish behaviorally. One possibility is a continuous acceptability signal that integrates reward magnitude, contextual value, and accumulated waiting cost. Another possibility is a more restricted signal tracking the scene- defined ordinal rank of the offered option within the current choice set. Our task was designed to separate these alternatives and thereby identify which form of evaluation is represented in the basal ganglia output.

We recorded well-isolated neurons in SNr and FEF while monkeys performed this task and found a clear functional dissociation. SNr activity was dominated by scene-defined ordinal rank, whereas FEF activity primarily reflected categorical commitment and movement direction. These results support a model in which context-dependent evaluation and categorical commitment are distributed across parallel output pathways converging on the SC.

## Results

### Monkeys flexibly accepted or rejected sequential offers based on scene-defined rank

Monkeys performed a sequential-offer choice task in which each visual scene defined a specific pair of available objects and thereby the scene-defined ordinal rank of each offer (Fig. 1A–C). We use “good” and “bad” as shorthand for higher-ranked and lower-ranked offers, respectively. Across scenes, monkeys reliably accepted the higher-ranked (“good”) object when it appeared, but they did not always wait for it. Rejection of the higher-ranked object was extremely rare (< 1% of good-offer trials across all monkeys), precluding reliable neural analysis. Accordingly, Reject Good trials were excluded from the outcome-based neural and SRT analyses. In scenes with smaller reward differences, monkeys frequently accepted the lower-ranked (“bad”) object (Accept Bad), indicating that choices reflected a trade-off between waiting for the higher-ranked offer and advancing to the next trial (Fig. 1D). This good-choice rate, the per-session proportion of completed trials in which the monkey accepted the higher-ranked object, was near ceiling in scenes with large reward differences and lower in scenes with smaller reward differences. Notably, the scenes in which the lower-ranked option was accepted differed across monkeys. Monkeys Ch and Cr accepted it mainly when the reward difference between the two options was small, whereas Monkey Sp also accepted it when the lower-ranked object itself carried a relatively large reward. This pattern might reflect a difference in how the monkeys weighed value, with Monkeys Ch and Cr relying more on the relative, scene-defined value of the offered object and Monkey Sp being additionally influenced by its absolute reward magnitude. Monkeys also expressed multiple strategies when rejecting lower-ranked offers. Notably, the predominant rejection strategy across scenes and monkeys was to make a rapid return saccade to central fixation (Fig. 1E), enabling us to analyze rejection as an active movement rather than the absence of movement.

**Figure 1.**
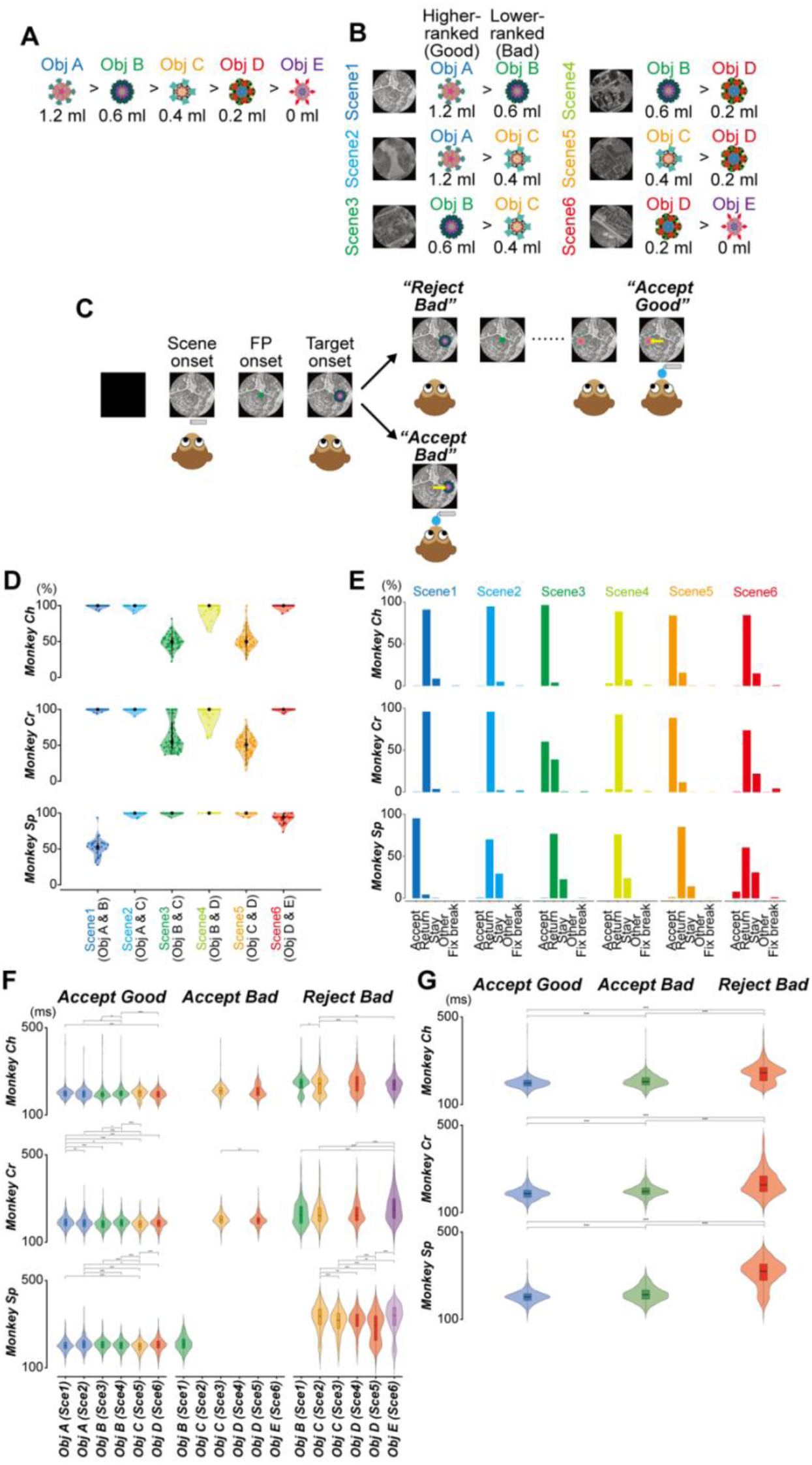
Task design and behavior in a context-dependent sequential-offer choice task. (A) Visual objects (Objects A–E) and associated reward amounts (Object A = 1.2 mL, B = 0.6 mL, C = 0.4 mL, D = 0.2 mL, E = 0 mL). (B) Choice scenes (Scenes 1–6). Each scene (background image) specifies a pair of available objects, thereby defining a higher-ranked (“good”) and lower-ranked (“bad”) option within that scene. (C) Trial structure. After scene presentation and fixation point (FP) onset, a single object is presented peripherally. Monkeys accepted an offer by making a saccade to the object and maintaining fixation (≥400 ms). Monkeys rejected an offer by not maintaining fixation on the object. Rejection could be expressed as a return saccade, staying at center, or other eye movements. Offers were repeated until acceptance. (D) Good-choice rate across scenes. For each scene (Scenes 1–6), the percentage of completed (accepted) trials on which the monkey accepted the higher-ranked (good) object, shown separately for each monkey. Each violin plot shows the distribution across recording sessions, where one session corresponds to the behavioral data collected during isolation of a single neuron. Values near 100% indicate that the monkey almost always obtained the higher-ranked object, whereas lower values indicate more frequent acceptance of the lower-ranked option (Accept Bad) in that scene. See Methods for additional details. (E) Distribution of behavioral outcomes across scenes and monkeys (accept, return, stay, other, and fixation break). Fixation breaks were excluded from all analyses and are plotted only to show how often they occurred. (F) Saccade reaction times (SRTs) by scene and outcome in the choice task (Scenes 1–6). Violin plots show SRT distributions for Accept Good, Accept Bad, and Reject Bad (return) trials for Monkey Ch, Monkey Cr, and Monkey Sp. Accept Bad and Reject Bad distributions are shown only for scenes with sufficient numbers of those trials (e.g., Accept Bad was prominent in Scenes 3 and 5 for Monkey Ch and Monkey Cr, and in Scene 1 for Monkey Sp). (G) Saccade reaction times pooled across all choice scenes (Scenes 1–6) for each outcome and monkey, showing the consistent latency ordering across Accept Good, Accept Bad, and Reject Bad (return) outcomes. Asterisks in (F) and (G) indicate corrected post hoc comparisons, *P < 0.05, **P < 0.01, ***P < 0.001, see Tables S1–S3.

### Saccade reaction times tracked decision outcome rather than scene context

To assess the time course of these decisions, we analyzed saccade reaction times (SRTs) during the choice task (Scenes 1–6). SRT was defined as the latency from target onset to the initiation of the first saccade from central fixation toward the offered object. For rejection trials, SRT was quantified only for return rejections (because stay rejections have no saccade latency), and non–task-directed eye movements were excluded.

SRTs differed strongly across behavioral outcomes (Fig. 1F, G). In all monkeys, Accept Good trials showed the shortest SRTs, Accept Bad trials were intermediate, and Reject Bad (return) trials exhibited the longest SRTs. This pattern provides behavioral evidence that the return saccade is not a reflexive response, but an action performed after evaluating the offered option in its scene context and deciding to reject it. Linear mixed-effects model (LMM) analysis confirmed that behavioral outcome exerted a robust main effect on SRT with large effect sizes in all monkeys (partial *η*² = 0.25–0.61, Table S2). In contrast, within-outcome differences across scenes were modest (partial *η*² ≤ 0.08), indicating that SRT variation was dominated by the decision outcome rather than scene context (Fig. 1F). As a control, in forced-choice trials (Scenes 7–10) where only a single object was available, SRTs were short and consistent, comparable to Accept Good responses (Fig. S1C), supporting the interpretation that the prolonged latencies in Accept Bad and Reject Bad reflect the cognitive demands of context-dependent evaluation and rejection planning rather than motor limitations.

**Figure S1.**
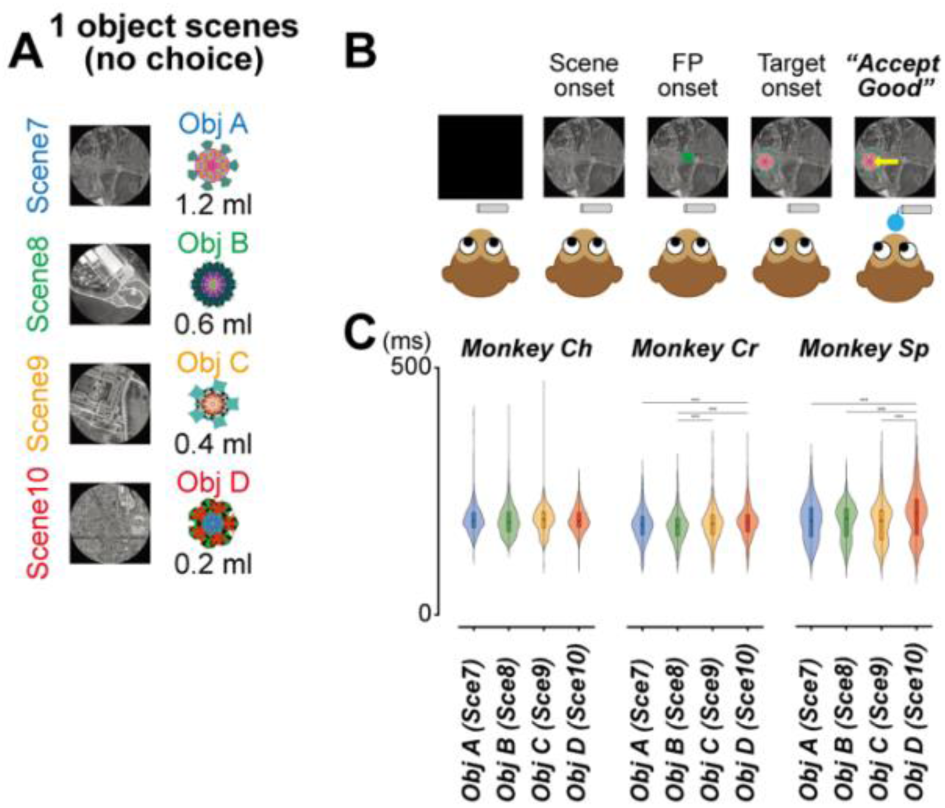
Behavior and SRTs in the forced-choice task. (A) Forced-choice scenes (Scenes 7–10) in which a single object was available (Objects A–D). (B) Trial structure for forced-choice trials. (C) SRT distributions in forced-choice trials for Monkey Ch, Monkey Cr, and Monkey Sp. Asterisks indicate corrected post hoc comparisons, *P < 0.05, **P < 0.01, ***P < 0.001. Descriptive statistics and statistical comparisons are provided in Tables S1–S3.

### SNr shows stronger scene-related modulation than FEF before target onset

Before examining how SNr and FEF encode specific offers during target evaluation, we first asked whether the two areas differ in their sensitivity to scene context prior to target onset. During the scene period (100–300 ms after scene onset), SNr neurons showed prominent scene-related modulation (Fig. 2A, C), indicating sensitivity to the current scene before target evaluation. In contrast, FEF neurons exhibited weaker modulation across scenes in the same window (Fig. 2B, D). A combined LMM analysis across monkeys confirmed this dissociation (Fig. 2C, D and Table S4). SNr activity was strongly modulated by the current scene (F = 176.1, partial η² = 0.66), whereas FEF activity showed substantially weaker scene-related modulation (F = 15.0, partial η² = 0.13). This dissociation was consistent in each monkey (Fig. S2). Post-hoc comparisons revealed that SNr scene-period activity clustered by reward context. Scenes 1 and 2, which both contained the highest-reward object (Object A), were indistinguishable from each other (P = 1.0) but differed strongly from Scenes 3–6 (P < 0.001, Table S4). Although our design does not fully dissociate scene identity from specific reward-related variables such as the best available reward or the scene value gap, this pattern is consistent with the interpretation that pre-target SNr modulation reflects the reward structure of the current scene.

**Figure 2.**
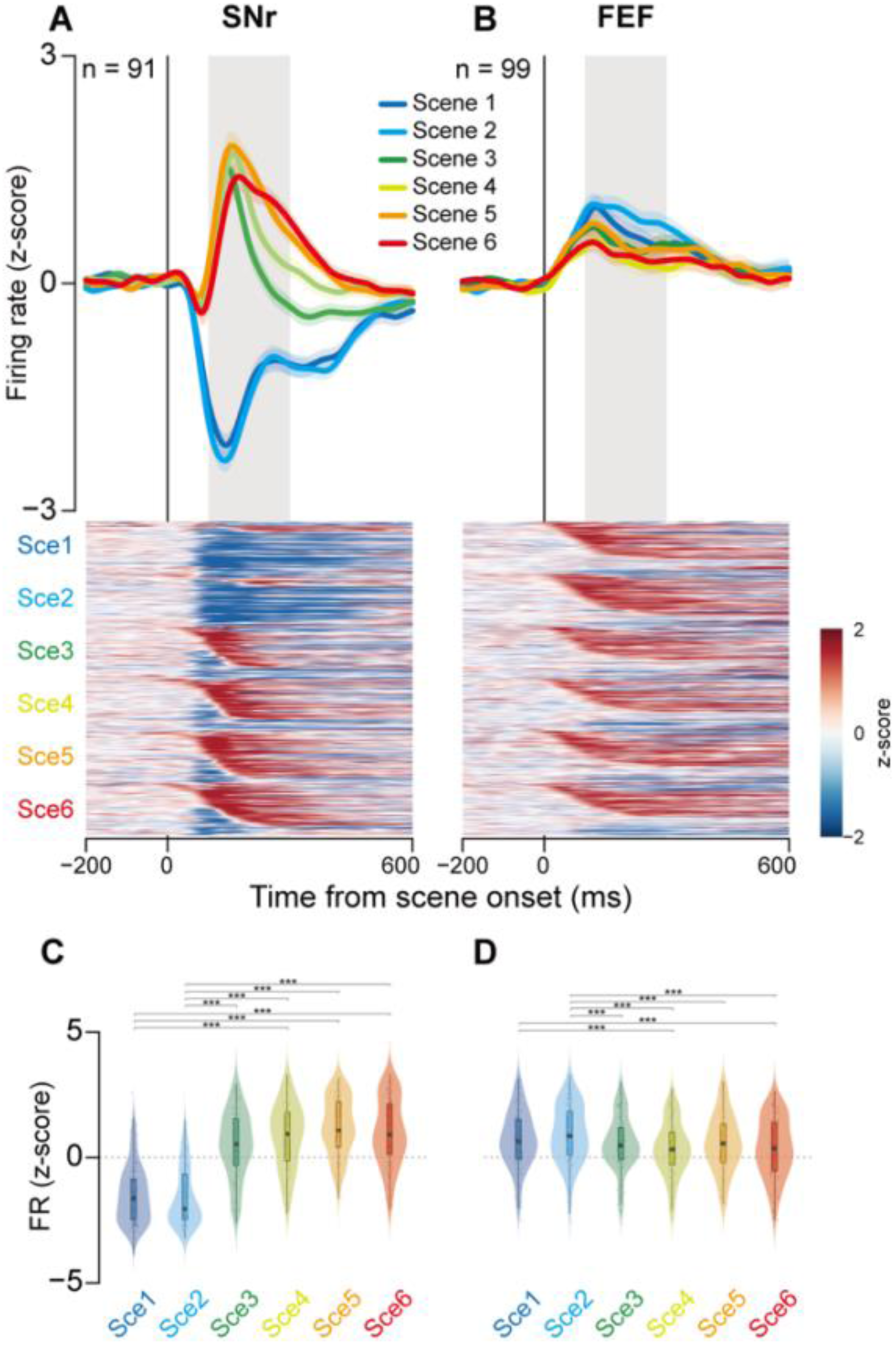
Scene-period activity in SNr and FEF during the choice task, Scenes 1–6. Neuronal activity was z-score normalized and pooled across the three monkeys. Good and Bad refer to the higher-ranked and lower-ranked offers, respectively, within the current scene. Per-monkey data are shown in Fig. S2. (A, B) Population mean activity (top) and heatmaps showing activity of individual neurons (bottom), aligned to scene onset for SNr (A, n = 91) and FEF (B, n = 99). Colored traces indicate Scenes 1–6. Shaded regions indicate the scene-period analysis window, 100–300 ms after scene onset. Heatmaps show z-scored activity of individual neurons grouped by scene. (C, D) Distributions of mean activity during the scene-period analysis window for each scene in SNr (C) and FEF (D). Violin plots show neuron-level distributions, and boxes indicate the median and interquartile range. Scene effects and post hoc pairwise comparisons were tested separately for SNr and FEF. Asterisks indicate corrected pairwise comparisons, *P < 0.05, **P < 0.01, ***P < 0.001. Detailed statistics are provided in Table S4.

**Figure S2.**
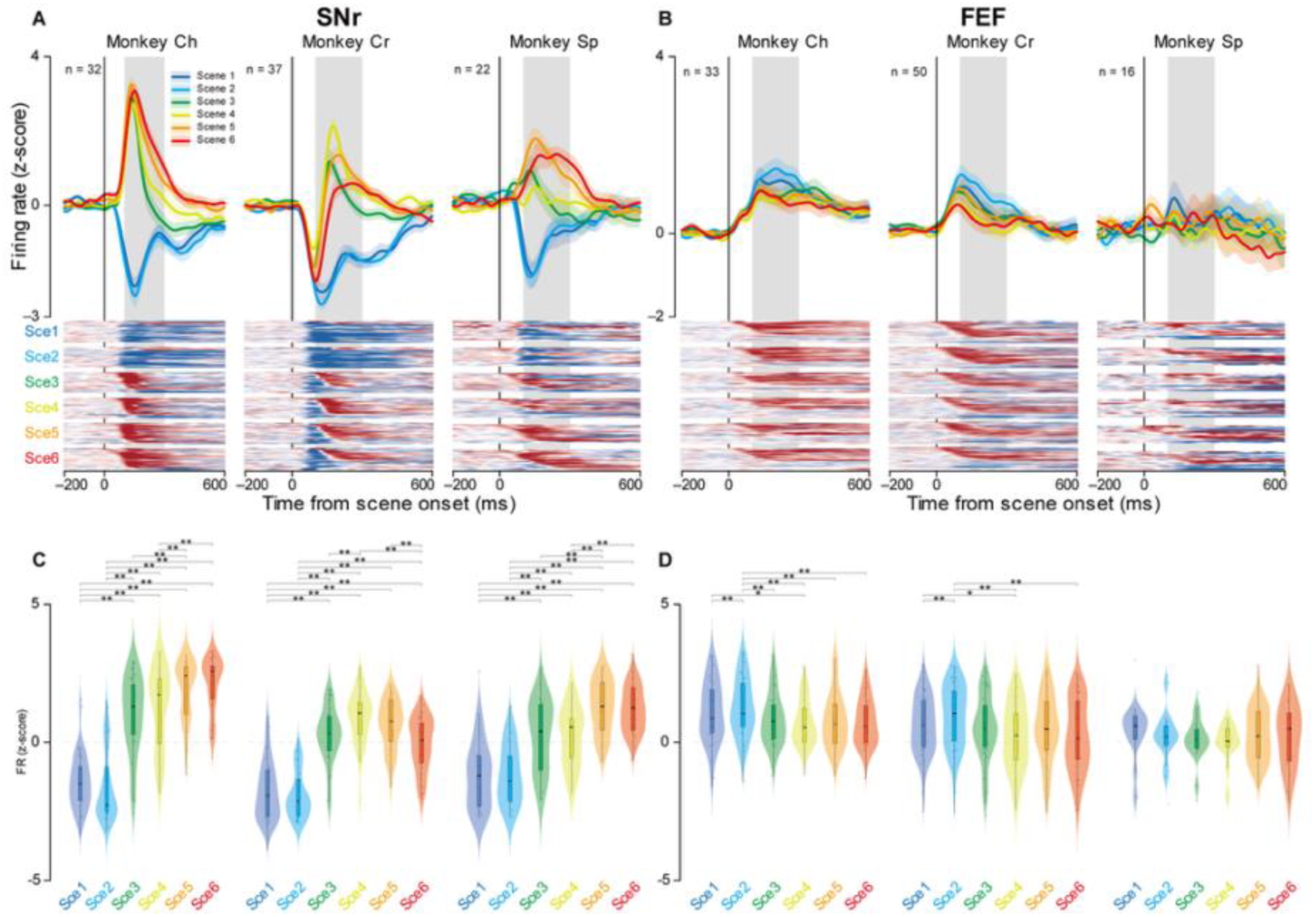
Scene-period activity in SNr and FEF for individual monkeys during the choice task, Scenes 1–6. Neuronal activity was z-score normalized. Data are shown separately for Monkey Ch, Monkey Cr, and Monkey Sp. Neuron counts per monkey are indicated in each panel. (A, B) Population mean activity (top) and heatmaps showing activity of individual neurons (bottom), aligned to scene onset for SNr (A) and FEF (B). Each column shows data from one monkey. Colored traces indicate Scenes 1–6. Shaded regions indicate the scene-period analysis window, 100–300 ms after scene onset. Heatmaps show z-scored activity of individual neurons grouped by scene. (C, D) Distributions of mean activity during the scene-period analysis window for each scene in SNr (C) and FEF (D). Violin plots show neuron-level distributions, and boxes indicate the median and interquartile range. Scene effects and post hoc pairwise comparisons were tested separately for each monkey and brain area. Asterisks indicate corrected pairwise comparisons, *P < 0.05, **P < 0.01, ***P < 0.001.

**Figure S3.**
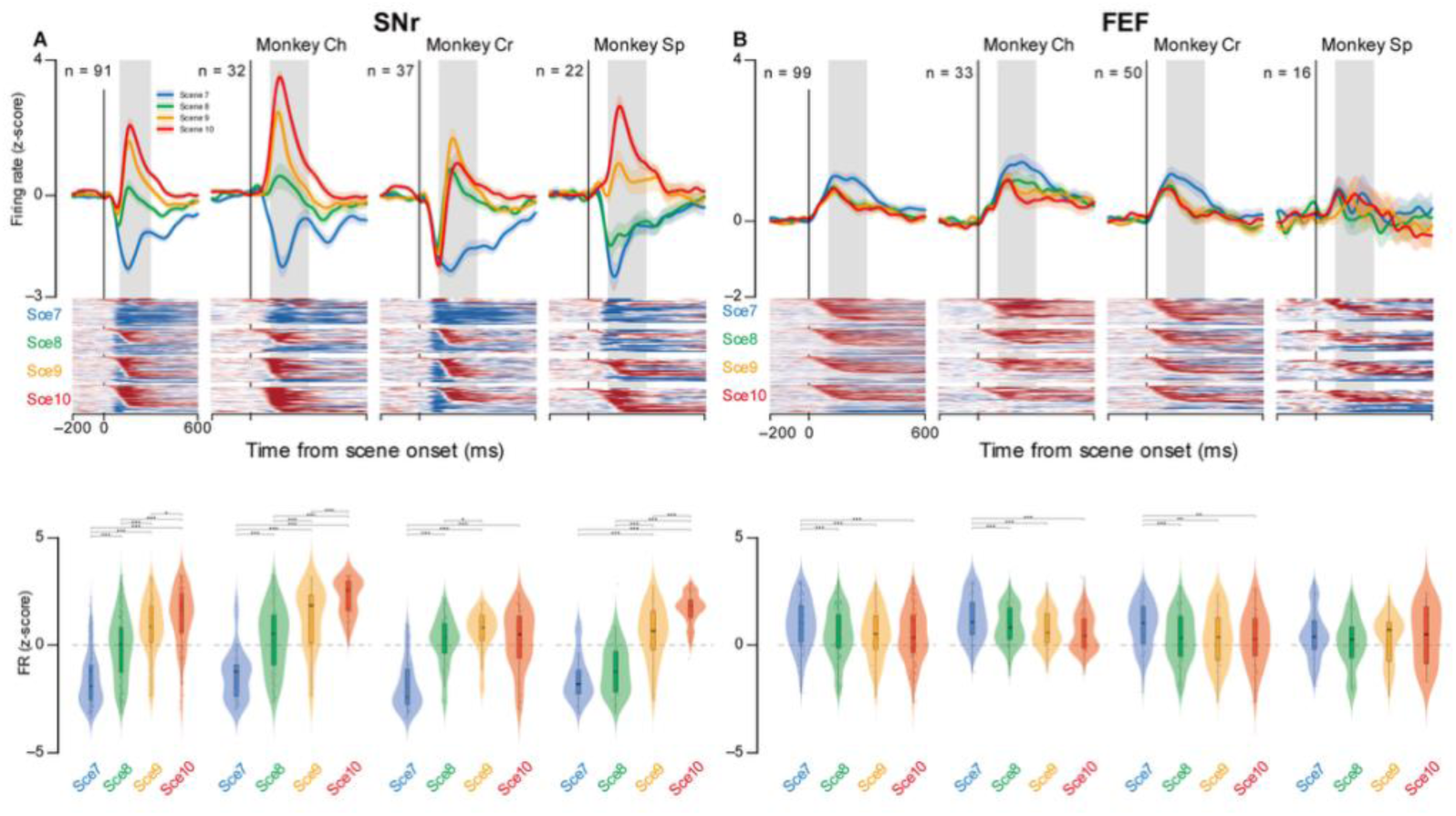
Scene-period activity in SNr and FEF during the forced-choice task, Scenes 7–10. Neuronal activity was z-score normalized. Scenes 7–10 were forced-choice scenes in which a single object was available, and all panels show accepted offers. Neuron counts are indicated in each panel. (A, B) Population mean activity, heatmaps, and violin plots aligned to scene onset for SNr (A) and FEF (B). For each brain area, data are shown first for the pooled three-monkey population, followed by Monkey Ch, Monkey Cr, and Monkey Sp. Colored traces indicate Scenes 7–10. Shaded regions indicate the scene-period analysis window, 100–300 ms after scene onset. Heatmaps show z-scored activity of individual neurons grouped by scene. Violin plots show neuron-level distributions of mean activity during the scene-period analysis window, and boxes indicate the median and interquartile range. Scene effects and post hoc pairwise comparisons were tested separately for the pooled data and for each monkey within each brain area. Asterisks indicate corrected pairwise comparisons, *P < 0.05, **P < 0.01, ***P < 0.001.

### SNr shows ordered modulation across behavioral outcomes, whereas FEF reflects categorical commitment during target evaluation

We then tested how activity in SNr and FEF was modulated by the offered options and decision outcomes during target evaluation. We analyzed neuronal activity in a fixed window (100–300 ms after target onset) for three outcomes: Accept Good, Accept Bad, and Reject Bad (return). In the SNr, activity showed an ordered pattern across outcomes (Fig. 3A, C). Firing was lowest for Accept Good, intermediate firing for Accept Bad, and highest firing for Reject Bad. The difference between Accept Good and Accept Bad was substantially larger than the difference between Accept Bad and Reject Bad, suggesting that the ordered pattern reflects primarily the rank distinction (higher-ranked vs lower-ranked) rather than the accept/reject distinction alone. In the FEF, activity was categorical (Fig. 3B, D). Accept Good and Accept Bad were similar, while Reject Bad was clearly separated. Importantly, the SNr ordered modulation and the FEF categorical separation of acceptance versus rejection were observed in each monkey for both contralateral and ipsilateral targets (Figs. S4 and S5), indicating that the population-level effects are not driven by a single subject. Because Reject Bad trials were quantified using return saccades, in which the monkey initiated a saccade toward the object before redirecting gaze back to center, this accept–reject dissociation in FEF cannot be reduced to a simple movement-versus-no-movement contrast.

**Figure 3.**
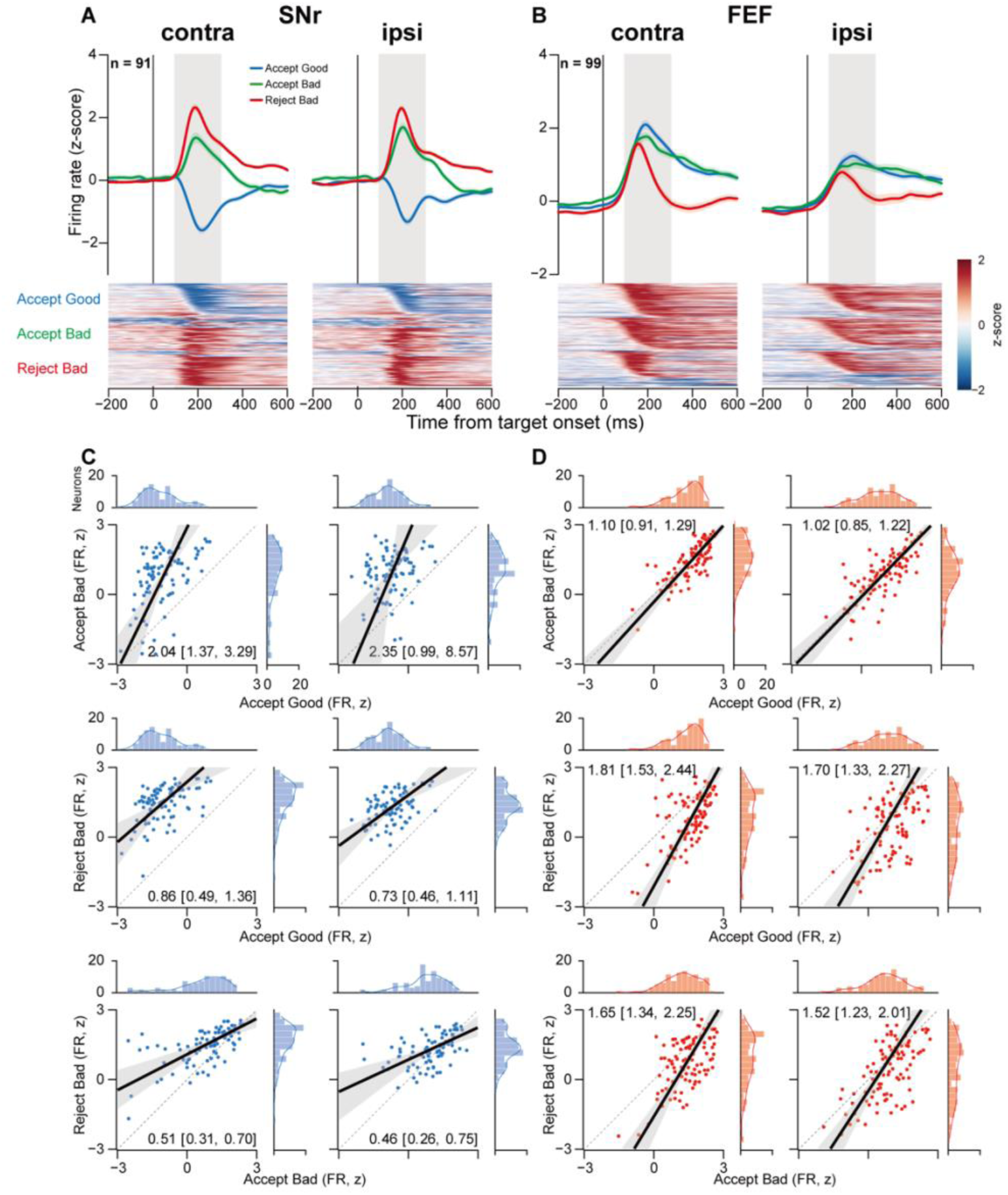
Target-period activity in SNr and FEF during the choice task, Scenes 1–6. Neuronal activity was z-score normalized and pooled across the three monkeys. Good and Bad refer to the higher-ranked and lower-ranked offers, respectively, within the current scene. Per-monkey data are shown in Figs. S4 and S5. (A, B) Population mean activity (top) and heatmaps showing activity of individual neurons (bottom), aligned to target onset for SNr (A, n = 91) and FEF (B, n = 99). Data are shown separately for contralateral and ipsilateral targets and for Accept Good, Accept Bad, and Reject Bad outcomes. Shaded regions indicate the target-period analysis window, 100–300 ms after target onset. Heatmaps show z-scored activity of individual neurons grouped by outcome. (C, D) Neuron-by-neuron comparisons of mean target-period activity for SNr (C) and FEF (D). Scatter plots compare Accept Bad versus Accept Good, Reject Bad versus Accept Good, and Reject Bad versus Accept Bad, separately for contralateral and ipsilateral targets. Each point represents one neuron. Marginal histograms and density curves show the distributions of activity values for each axis. Dotted diagonal lines indicate equality between the two conditions. Black lines show Deming fits, and gray bands indicate bootstrap 95% confidence bands for the Deming fit. Values in each scatter plot indicate the Deming regression slope and bootstrap 95% confidence interval. Paired condition differences were tested using two-sided paired sign-flip permutation tests, with Holm correction across the 12 scatter comparisons (2 regions × 2 target directions × 3 condition contrasts). Bootstrap confidence intervals were used to estimate effect sizes. Detailed statistical results are shown in Tables S5 and S6.

Neuron-by-neuron paired comparisons, tested with two-sided sign-flip permutation tests and Holm correction, statistically verified this dissociation (Fig. 3C, D and Table S5). In SNr, target-period activity differed reliably even between the two acceptance outcomes. The within-neuron Accept Bad minus Accept Good difference was large and positive (mean 1.85 [95% CI 1.63, 2.06] for contralateral and 1.66 [1.47, 1.85] for ipsilateral targets, both P < 0.001) and exceeded the smaller Reject Bad minus Accept Bad difference (0.65 [0.48, 0.84] contralateral, 0.32 [0.18, 0.48] ipsilateral). In FEF, by contrast, the Accept Good versus Accept Bad difference was small (Accept Bad minus Accept Good, −0.18 [−0.27, −0.08] contralateral and −0.09 [−0.22, 0.03] ipsilateral) and much smaller than the difference distinguishing acceptance from rejection (Reject Bad minus Accept Good, −0.80 [−0.97, −0.63] contralateral and −0.55 [−0.75, −0.36] ipsilateral, both P < 0.001), consistent with a categorical separation of acceptance and rejection rather than a graded rank code. Control analyses in forced-choice trials (Scenes 7–10) showed that SNr maintained modulation by reward amount for single objects, whereas FEF modulation by reward amount was comparatively minor (Figs. S6 and S7), supporting the interpretation that this dissociation is not attributable to differences in visual input or movement execution.

**Figure S4.**
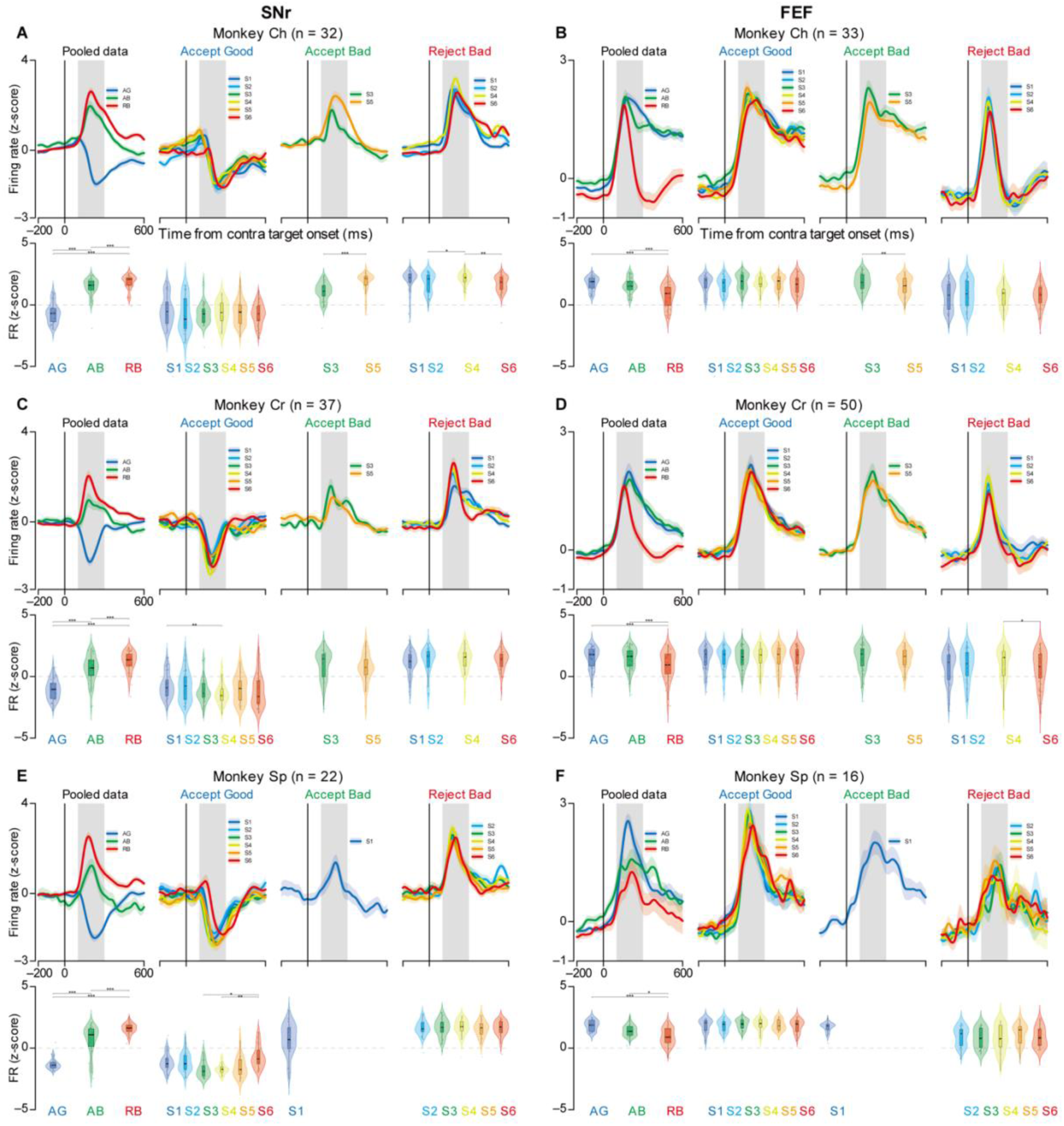
Contra target-period activity in SNr and FEF for individual monkeys during the choice task, Scenes 1–6. Neuronal activity was z-score normalized and aligned to contra target onset. Good and Bad refer to the higher-ranked and lower-ranked offers, respectively, within the current scene. Neuron counts per monkey are indicated in each panel. (A–F) Population mean activity and violin plots for SNr and FEF in Monkey Ch, Monkey Cr, and Monkey Sp. SNr data are shown on the left (A, C, and E), and FEF data are shown on the right (B, D, and F). Within each monkey and brain area, panels show pooled data across Scenes 1–6 for Accept Good (AG), Accept Bad (AB), and Reject Bad (RB), followed by scene-specific activity for Accept Good, Accept Bad, and Reject Bad. Scene-specific Accept Bad and Reject Bad panels include scenes with sufficient data for that monkey and condition. Lines and shaded bands show population mean activity and SEM. Gray shading indicates the target-period analysis window, 100–300 ms after target onset. Violin plots show neuron-level distributions of mean activity during the target-period analysis window, and boxes indicate the median and interquartile range. Condition effects in the pooled panels and scene effects in the scene-specific panels were tested separately for each monkey and brain area. Asterisks indicate corrected pairwise comparisons, *P < 0.05, **P < 0.01, ***P < 0.001. Detailed statistics are provided in Tables S9 and S10.

**Figure S5.**
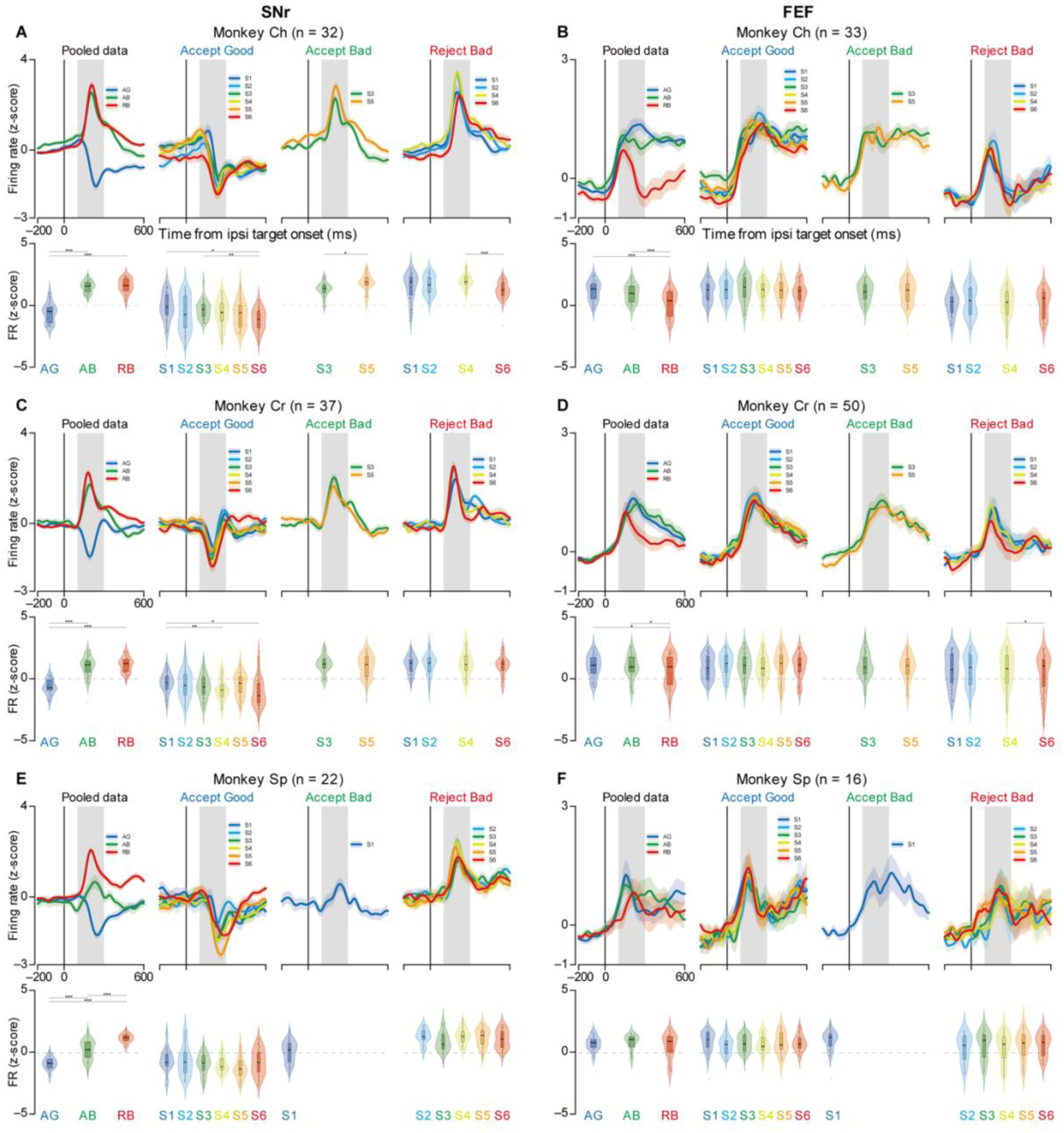
Ipsi target-period activity in SNr and FEF for individual monkeys during the choice task, Scenes 1–6. Neuronal activity was z-score normalized and aligned to ipsi target onset. Good and Bad refer to the higher-ranked and lower-ranked offers, respectively, within the current scene. Neuron counts per monkey are indicated in each panel. (A–F) Population mean activity and violin plots for SNr and FEF in Monkey Ch, Monkey Cr, and Monkey Sp. SNr data are shown on the left (A, C, and E), and FEF data are shown on the right (B, D, and F). Within each monkey and brain area, panels show pooled data across Scenes 1–6 for Accept Good (AG), Accept Bad (AB), and Reject Bad (RB), followed by scene-specific activity for Accept Good, Accept Bad, and Reject Bad. Scene-specific Accept Bad and Reject Bad panels include scenes with sufficient data for that monkey and condition. Lines and shaded bands show population mean activity and SEM. Gray shading indicates the target-period analysis window, 100–300 ms after target onset. Violin plots show neuron-level distributions of mean activity during the target-period analysis window, and boxes indicate the median and interquartile range. Condition effects in the pooled panels and scene effects in the scene-specific panels were tested separately for each monkey and brain area. Asterisks indicate corrected pairwise comparisons, *P < 0.05, **P < 0.01, ***P < 0.001. Detailed statistics are provided in Tables S11 and S12.

**Figure S6.**
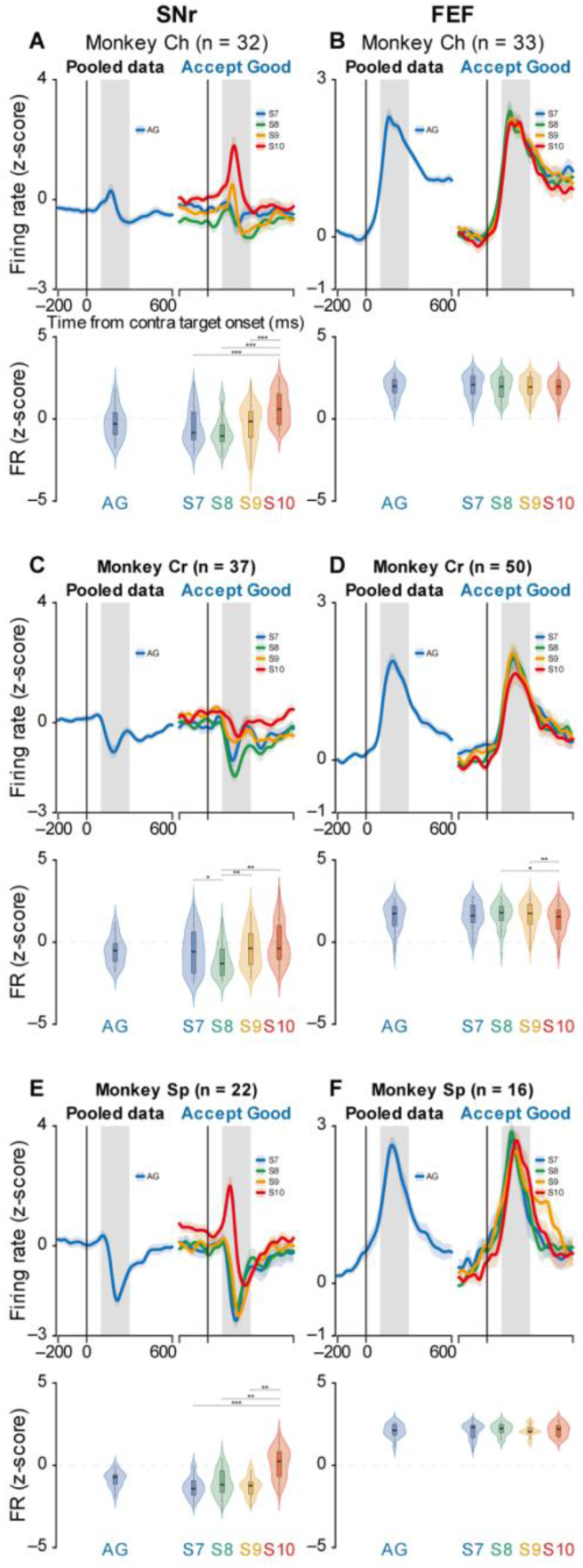
Contra target-period activity in SNr and FEF for individual monkeys during the forced-choice task, Scenes 7–10. Neuronal activity was z-score normalized and aligned to contra target onset. Scenes 7–10 were forced-choice scenes in which a single object was available, and all panels show accepted offers. Neuron counts per monkey are indicated in each panel. (A–F) Population mean activity and violin plots for SNr and FEF in Monkey Ch, Monkey Cr, and Monkey Sp. SNr data are shown on the left (A, C, and E), and FEF data are shown on the right (B, D, and F). Within each monkey and brain area, panels show pooled data across Scenes 7–10 for Accept Good (AG), followed by scene-specific Accept Good activity for Scenes 7–10. Lines and shaded bands show population mean activity and SEM. Gray shading indicates the target-period analysis window, 100–300 ms after target onset. Violin plots show neuron-level distributions of mean activity during the target-period analysis window, and boxes indicate the median and interquartile range. The pooled AG panel contains a single condition and was summarized descriptively. Scene effects and post hoc pairwise comparisons in the scene-specific panels were tested separately for each monkey and brain area. Asterisks indicate corrected pairwise comparisons, *P < 0.05, **P < 0.01, ***P < 0.001. Detailed statistics are provided in Table S13.

**Figure S7.**
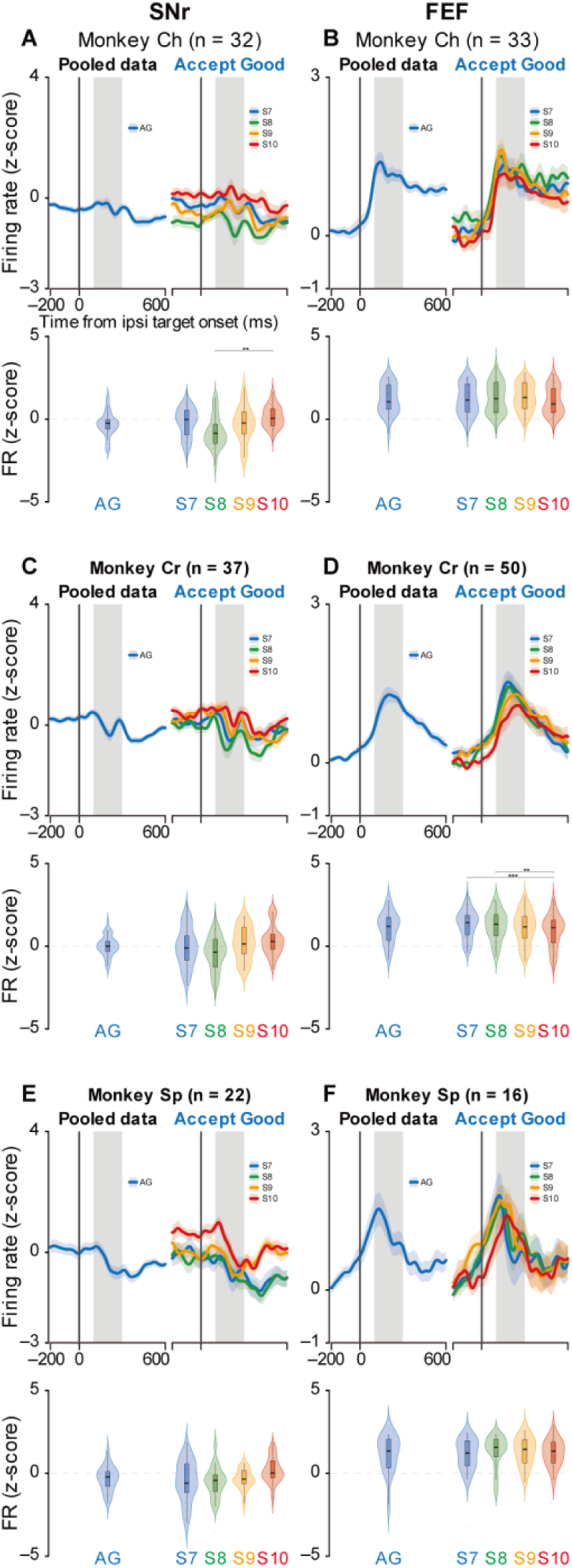
Ipsi target-period activity in SNr and FEF for individual monkeys during the forced-choice task, Scenes 7–10. Neuronal activity was z-score normalized and aligned to ipsi target onset. Scenes 7–10 were forced-choice scenes in which a single object was available, and all panels show accepted offers. Neuron counts per monkey are indicated in each panel. (A–F) Population mean activity and violin plots for SNr and FEF in Monkey Ch, Monkey Cr, and Monkey Sp. SNr data are shown on the left (A, C, and E), and FEF data are shown on the right (B, D, and F). Within each monkey and brain area, panels show pooled data across Scenes 7–10 for Accept Good (AG), followed by scene-specific Accept Good activity for Scenes 7–10. Lines and shaded bands show population mean activity and SEM. Gray shading indicates the target-period analysis window, 100–300 ms after target onset. Violin plots show neuron-level distributions of mean activity during the target-period analysis window, and boxes indicate the median and interquartile range. The pooled AG panel contains a single condition and was summarized descriptively. Scene effects and post hoc pairwise comparisons in the scene-specific panels were tested separately for each monkey and brain area. Asterisks indicate corrected pairwise comparisons, *P < 0.05, **P < 0.01, ***P < 0.001. Detailed statistics are provided in Table S14.

### Distinct task-variable coding quantified by multivariable mixed-effects modeling

Because behavioral outcomes reflect a combination of multiple task factors, we decomposed each offer into component variables for model-based analysis. Specifically, Accept Good and Accept Bad share the same action (accept) but differ in rank (higher-ranked vs lower-ranked within the current scene), whereas Accept Bad and Reject Bad share the same rank class (lower-ranked) but differ in commitment (accept vs reject). In addition, the same rank label can correspond to different reward amounts across scenes and targets can appear in either hemifield, motivating a multivariable model that simultaneously accounts for Action, Rank, Reward amount, and Direction.

To disentangle correlated task factors and quantify their independent contributions, we fit a multivariable LMM to neuronal activity in the extended target window (100–400 ms after target onset; Fig. 4A). The model included fixed effects for Action (accept vs reject), Rank (higher-ranked vs lower-ranked within the current scene), Reward amount (reward magnitude), Direction (contralateral vs ipsilateral), and trial-by-trial SRT, with a random intercept for neuron. In SNr, Rank was the dominant predictor (*β* = −0.64, *P* < 0.001), with smaller but significant effects of Action (*β* = −0.20, *P* < 0.001) and Reward amount (*β* = 0.03, *P* < 0.001). In contrast, FEF activity was dominated by Direction (*β* = 0.66, *P* < 0.001) and Action (*β* = 0.35, *P* < 0.001), while Rank was negligible and not significant (*β* = 0.03, *P* = 0.30), consistent with the categorical representation observed in the outcome-based analyses.

**Figure 4.**
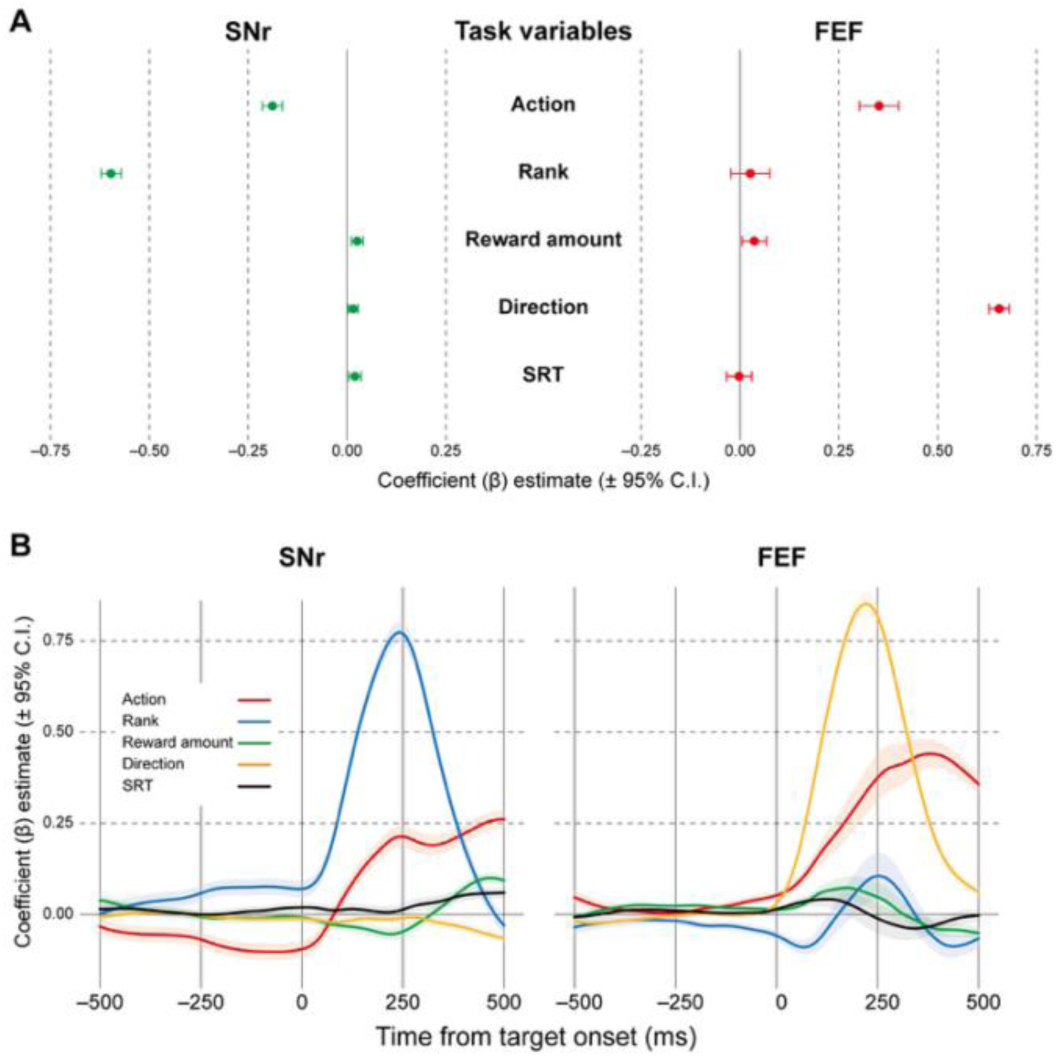
Multivariable mixed-effects modeling dissociates task-variable coding in SNr and FEF. (A) Fixed-effect coefficient estimates (β) from linear mixed-effects models fit to target-period activity (100–400 ms after target onset). The model included Action (accept versus reject), Rank (higher-ranked versus lower-ranked within the current scene), Reward amount (reward magnitude), Direction (contralateral versus ipsilateral), and trial-by-trial saccade reaction time (SRT, log-transformed and standardized within monkey) as fixed effects, with random intercepts for monkey and neuron. Coefficients are shown for SNr (n = 91 neurons) and FEF (n = 99 neurons) separately, and error bars indicate 95% confidence intervals. Analyses were restricted to trials with measurable SRT. Detailed fixed-effect estimates are shown in Table S7. (B) Temporal dynamics of task-variable coding obtained by fitting the same linear mixed-effects model in sliding windows aligned to target onset (window width 200 ms, step 10 ms). Lines show fixed-effect estimates (β) over time, and shaded bands indicate 95% confidence intervals. The vertical line indicates target onset (0 ms), and the horizontal dashed line indicates β = 0. For visual comparison with FEF, SNr coefficients for Action, Rank, Reward amount, and Direction were sign-inverted, such that positive deflections correspond to stronger encoding in the same direction as the FEF traces. This inversion was applied for visualization only, because higher-ranked and accepted options were associated with reduced firing in this inhibitory output pathway. All statistical analyses used the original coefficient signs, which are reported in Figure 4A and Table S7. The SRT coefficient was not sign-inverted and is plotted with its original sign.

To assess whether outcome-related latency differences could account for this dissociation, we examined the SRT effect within the same multivariable model (log-transformed and standardized within monkey), restricting the analysis to trials with measurable SRT. The SRT term was negligible in FEF (*P* = 0.88), and SNr remained dominated by Rank, with only a small additional SRT effect (*P* = 0.011), indicating that the dissociation between SNr and FEF cannot be explained by latency differences.

A sliding-window implementation of the same model revealed distinct temporal dynamics (Fig. 4B). In SNr, Rank encoding was strongest around 200–250 ms after target onset, exceeding the magnitude of the Action signal. In FEF, Action and Direction signals were strongest around 250 ms, while Rank remained near zero throughout. Note that SNr coefficients in Fig. 4B are plotted with inverted sign for visual comparison with FEF (see Fig. 4 legend).

Because the 200-ms analysis window limits temporal resolution, we interpret these dynamics in terms of the relative dominance of each signal rather than precise onset latencies. Nonetheless, these dynamics are consistent with rank-dominant evaluative coding in SNr and categorical commitment coding in FEF.

While these multivariable analyses identify Rank as the dominant predictor among experimenter-defined variables, they do not address whether the SNr ordered pattern reflects a discrete rank code or a graded continuous signal tracking behavioral acceptability. To distinguish between these possibilities, we next used behavioral modeling to derive a continuous acceptability measure from the monkeys’ own choice data and tested whether this composite variable, or its individual components, better explained neuronal activity in SNr versus FEF.

### A behavioral model captures context-dependent choice

We first asked which task factors determine whether a monkey accepts or rejects a given offer. We fit logistic regression models of increasing complexity to the binary accept/reject choice on each offer (Fig. 5A). The simplest model used only the reward amount of the offered object. Additional models incorporated different combinations of predictors, including scene identity, the reward difference between the good and bad options in the current scene (scene value gap), the number of prior rejections within the trial, and the ordinal rank of the offered object. Models ranged from single-predictor (reward amount only) to a full model including all four factors. Each model was evaluated using session-level cross-validated log-likelihood as the primary metric, and the Akaike Information Criterion (AIC) was also computed to penalize models with additional predictors, with AIC values shown in Fig. 5A (see Methods for model specifications and evaluation).

**Figure 5.**
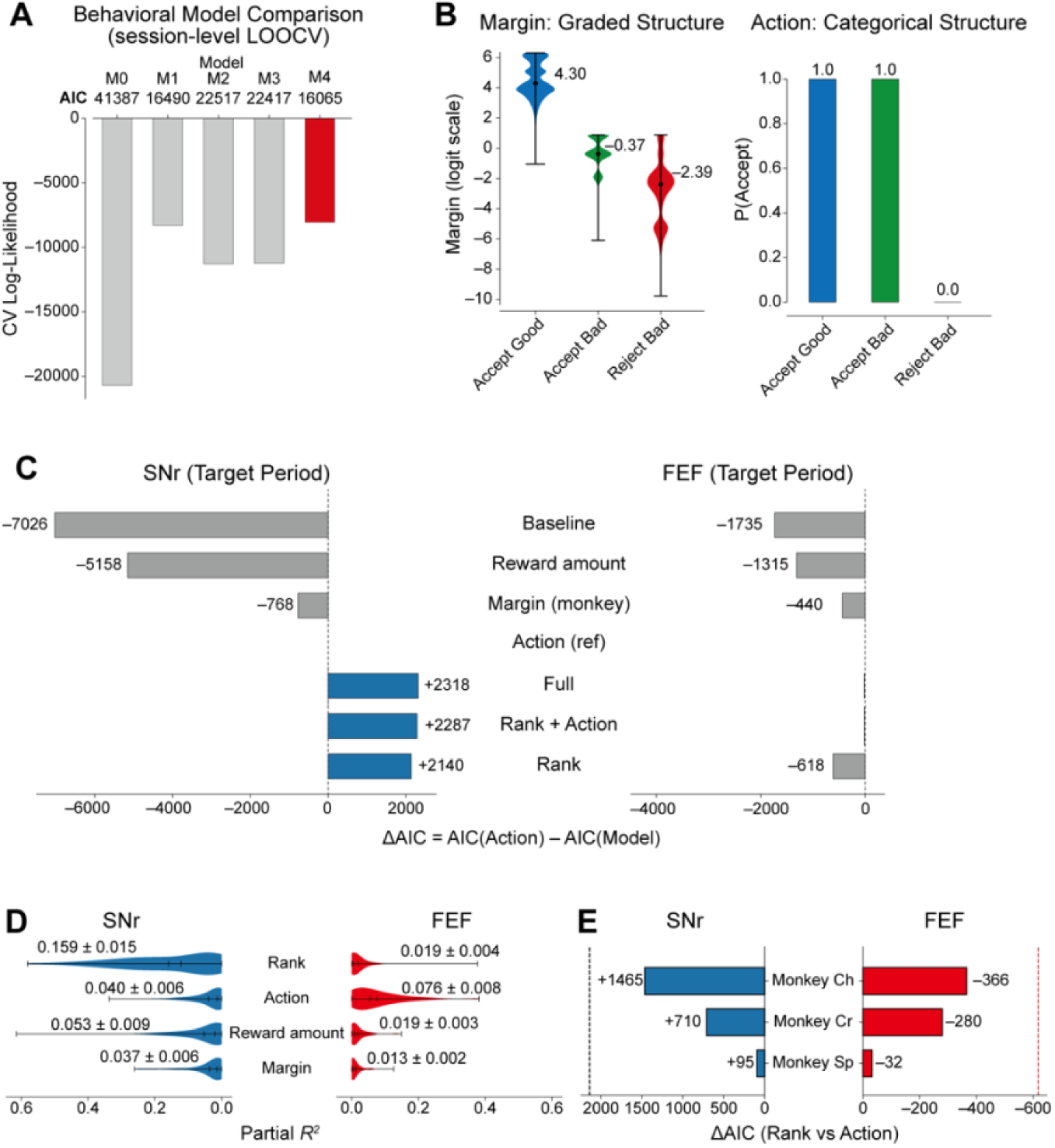
Behavioral and neural modeling identifies ordinal rank as the dominant variable explaining SNr activity. (A) Behavioral model comparison. Session-level leave-one-out cross-validated log-likelihood for logistic regression models predicting offer-level accept or reject decisions. Models ranged from reward amount only (M0) to a full model incorporating reward amount, scene value gap, number of prior rejections, and ordinal rank (M4, red). AIC values from full-data fits are shown for each model. (B) Distributions of Margin (left) and Action (right) across the three behavioral outcomes. Margin, derived from the best behavioral model (M4), showed a graded structure across Accept Good, Accept Bad, and Reject Bad (return) trials. Action was categorical and distinguished acceptance from rejection. (C) Neural model comparison for target-period activity (100–300 ms after target onset). ÄAIC values, defined as AIC(Action model) − AIC(test model), are shown for SNr (left, n = 91 neurons) and FEF (right, n = 99 neurons). Positive values indicate a better fit than the Action model. Models tested included Rank, Reward amount, Margin (monkey-specific), combined models (Rank + Action and Full), and a Baseline model (Direction only). In SNr, Rank, Rank + Action, and the Full neural model (Action + Rank + Reward amount + Direction) were preferred over Action (blue bars), whereas Reward amount and Margin provided a worse fit than Action (gray bars). In FEF, no model was preferred over Action. (D) Single-neuron variance explained. Violin plots show partial R² for Rank, Action, Reward amount, and Margin in SNr (left, blue) and FEF (right, red). Black markers and lines indicate the mean and 95% confidence intervals. In SNr, Rank explained substantially more variance than the other variables. In FEF, Action explained the most variance. (E) Monkey-level consistency. ΔAIC (Rank versus Action) is shown separately for each monkey and brain area. In SNr (left, blue), Rank was preferred over Action in all three monkeys. In FEF (right, red), Action was preferred in all three monkeys. Dashed lines indicate the pooled ΔAIC values.

The best-performing model included all four factors, substantially outperforming reward amount alone in both cross-validated log-likelihood (−8,056 vs −20,698) and AIC (Fig. 5A), confirming that the same object could be accepted or rejected depending on the scene context in which it appeared. We used this model to compute a single continuous score for each offer, termed Margin, representing the overall tendency of the monkey to accept that offer given all relevant task factors (see Methods). A high Margin indicates the offer is very likely to be accepted. A low Margin indicates it is likely to be rejected. We also defined a binary variable, Action, recording what the monkey actually did on each offer (accept = 1, reject = 0).

Margin exhibited a graded structure across the three behavioral outcomes. It was highest for Accept Good offers (median, 4.30), intermediate for Accept Bad (−0.37), and lowest for Reject Bad (−2.39, Fig. 5B, left). By contrast, Action was identical for Accept Good and Accept Bad (both = 1) and different only for Reject Bad (= 0; Fig. 5B, right). The graded structure of Margin and the categorical structure of Action thus provide two candidate explanatory variables for neuronal activity. One distinguishes all three conditions along a continuum, and the other distinguishes only acceptance from rejection.

### SNr activity is better explained by ordinal rank than by overall behavioral acceptability

The ordered SNr pattern across Accept Good, Accept Bad, and Reject Bad could reflect either a continuous signal tracking overall behavioral acceptability or a signal dominated by a specific component of evaluation. To distinguish these possibilities, we used a two-step strategy. We first tested whether the composite behavioral acceptability score (Margin) outperforms the binary variable Action, then decomposed Margin into its individual components.

We first asked whether SNr and FEF activity in the evaluation window (100–300 ms after target onset) was better explained by Margin or by Action, using the difference in AIC (ΔAIC) as the primary comparison metric, the same information criterion used for the behavioral model comparison in Fig. 5A (see Methods). Before applying this comparison to real data, we verified through simulation that the analysis could reliably distinguish graded from categorical neural signals at the sample sizes and noise levels present in our data. Simulation recovery rate at empirical parameter values was 100% for both signal types, and recovery exceeded 80% across all robustness conditions tested (see Methods). Because our goal was to relate neuronal activity to within-monkey behavioral evaluation, our primary analysis used Margin values computed separately for each monkey, thereby removing between-monkey differences in overall acceptance tendency. Under this formulation, Margin did not outperform Action in SNr. Instead, Action provided the better account in all three monkeys. In FEF, Action was also preferred, consistent with the view that frontal cortical output primarily reflects categorical commitment.

Having ruled out the composite Margin, we next asked which individual component drives the SNr signal. Margin is a weighted sum of four task factors (reward amount, scene value gap, number of prior rejections, and ordinal rank). Among these, we focused on ordinal rank and reward amount as the two components most relevant to the SNr dissociation, and tested each against both Action and Margin (Fig. 5C). In SNr, ordinal rank alone was strongly preferred over both Action (ΔAIC = +2,140 relative to Action) and Margin (ΔAIC = +2,908 relative to Margin), while reward amount provided a substantially worse fit than Action (ΔAIC = −5,158). In FEF, no single variable or combination was preferred over Action (Rank, ΔAIC = − 618). This pattern was consistent at the single-neuron level. The mean variance explained by rank in SNr neurons (partial *R*² = 0.159) was approximately four times larger than that explained by action (0.040), whereas in FEF neurons, action explained the most variance (0.076, Fig. 5D). To formalize this dissociation, we summarized each neuron by a signed index, Δ partial R² = partial R²(Rank) − partial R²(Action), so that positive values denote neurons better explained by rank and negative values denote neurons better explained by action. This index was shifted positive in SNr (median = 0.080, with 82% of neurons above zero) and negative in FEF (median = −0.045, with 18% above zero). The between-area difference was large and highly reliable (Mann–Whitney *U* = 7,913, *P* = 1.1 × 10⁻¹⁹, Cohen’s *d* = 1.40), confirming a double dissociation in which rank dominates single-neuron coding in SNr and action dominates it in FEF.

Importantly, the rank-versus-action dissociation was consistent across all three monkeys (Fig. 5E). In SNr, rank was preferred over action in every monkey (ΔAIC: Monkey Ch, +1,465; Monkey Cr, +710; Monkey Sp, +95), whereas in FEF, action was preferred in every monkey (ΔAIC: Monkey Ch, −366; Monkey Cr, −280; Monkey Sp, −32). The failure of Margin to outperform Action, combined with the strong preference for Rank alone, indicates that the non-rank components of Margin (reward amount, scene value gap, prior rejections) do not contribute additively to SNr activity and, when pooled with Rank in the Margin variable, dilute rather than enhance the neural signal. Together, these analyses converge on ordinal rank, that is, whether the current offer is the higher-ranked or lower-ranked option in the scene, rather than the full behavioral acceptability of the offer, as the dominant coding variable in SNr. The large difference between Accept Good and Accept Bad therefore reflects the rank distinction, whereas the smaller difference between Accept Bad and Reject Bad reflects a more modest action-related contribution.

### Single-neuron response profiles are consistent with region-specific functional specializations

The preceding population-level and modeling analyses consistently identified ordinal rank as the dominant coding variable in SNr and categorical commitment as the dominant variable in FEF. We next asked whether these dissociations were also reflected in the response profiles of individual neurons. We classified neurons by the task variable showing the strongest modulation in the time-resolved model (Fig. 6). In SNr, the largest fraction of neurons was assigned to the Rank type (70%), whereas fewer neurons were assigned to the Action type (13%). Because this classification was based on temporally smoothed time-resolved coefficients, these fractions should be interpreted as descriptive summaries of the dominant trend across neurons rather than as exact inferential estimates. In FEF, neurons were predominantly Action type (54%) and Direction type (23%), with smaller fractions classified as Reward amount type (5%) and Rank type (4%), and the remainder unclassified (11%). Population heatmaps and subtype-averaged responses showed that SNr Rank-type neurons showed ordered modulation across outcomes, whereas FEF Action-type neurons showed categorical separation of acceptance versus rejection (Fig. 6B, D). Together, these single-neuron response patterns reinforced the division of labor identified by the population-level and model-based analyses, with the dominant neuron type in each area (Rank in SNr, Action in FEF) matching the variable that dominated its population and model-based coding. The agreement across population, model-based, and single-neuron analyses indicates that the SNr–FEF dissociation reflects the coding of individual neurons rather than only the pooled population.

**Figure 6.**
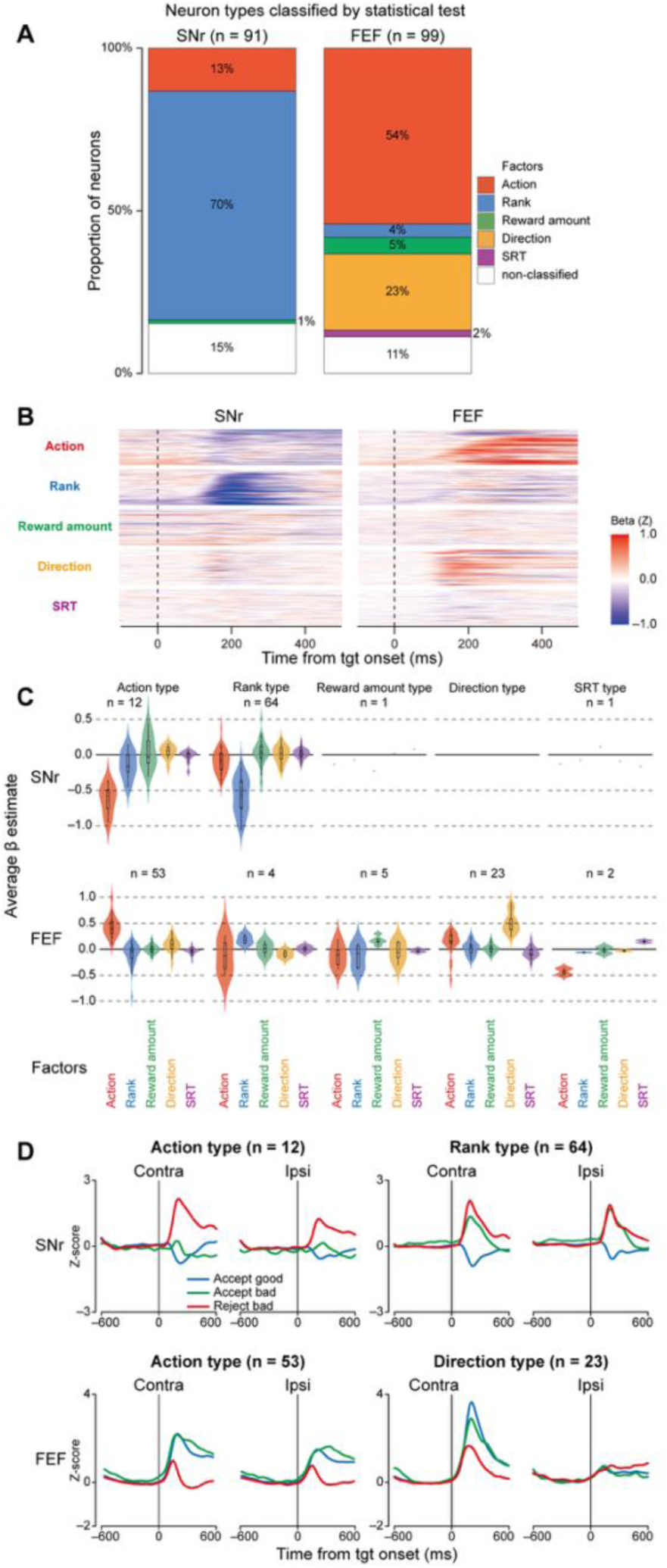
Classification of single-neuron properties based on task-variable encoding. (A) Proportions of neurons classified by their dominant significant task-variable modulation for SNr (n = 91) and FEF (n = 99). Classification was based on paired comparisons of variable-specific β time courses in the 100–400 ms target-period window across Action, Rank, Reward amount, Direction, and SRT, with Bonferroni correction for the 10 pairwise comparisons among the five variables (α = 0.005). (B) Population heatmaps of time-resolved â coefficients for each neuron. Colors indicate â coefficient magnitude in z-score units (red for positive and blue for negative coefficients). Within each brain area, the same neuron order was used for all task-variable rows. Neurons are ordered by the latency of Rank-related β modulation in SNr and Action-related β modulation in FEF. (C) Distributions of mean β estimates for each neuron type. Mean β estimates were calculated over the 100–400 ms target-period window. Each violin plot shows the distribution across neurons within a classified neuron type. Colors indicate the task variable corresponding to each β estimate. Summary statistics are provided in Table S8. (D) Mean population activity for classified neuron types with sufficient sample sizes for population-level visualization. Traces show target-aligned z-scored activity for contra and ipsi targets, with baseline calculated from 500 ms before target onset to target onset. The top row shows SNr Action type (left) and Rank type (right). The bottom row shows FEF Action type (left) and Direction type (right). Traces are colored by behavioral outcome (blue for Accept Good, green for Accept Bad, and red for Reject Bad). SNr Rank-type neurons show ordered activity (lowest for Accept Good, intermediate for Accept Bad, and highest for Reject Bad), whereas FEF Action-type neurons show categorical separation of acceptance and rejection outcomes. SNr Action-type neurons and FEF Direction-type neurons are shown as additional sufficiently sampled subpopulations with distinct modulation patterns.

## Discussion

Our study reveals a functional dissociation between parallel basal ganglia and frontal cortical outputs during choice. SNr activity was dominated by scene-defined ordinal rank, whereas FEF activity primarily reflected categorical commitment and target direction. Below we consider three aspects of this finding. First, we describe how the task design enabled the dissociation of rank from commitment. Second, we interpret the distinct coding schemes of SNr and FEF in relation to their known anatomical and physiological properties. Third, we discuss how these parallel outputs may converge at the superior colliculus to produce a commitment to action.

### Dissociating ordinal rank from categorical commitment

This task was designed to address a specific limitation of our recent scene-cued sequential choice paradigm (Yoshida & Hikosaka, 2024, 2025a, 2025b; Yoshida et al., 2026). The earlier studies used a binary structure in which each scene specified a rewarded and a nonrewarded object, and the same object could reverse its reward value across scenes. The present task replaces this binary structure with an ordinal rank structure in which lower-ranked offers could be accepted when rank differences were small. As in the earlier paradigm, rejection was expressed through active return saccades rather than movement omission, which allowed us to analyze rejection as an active decision rather than the absence of action. Together, the ordinal rank structure and the active rejection response partially dissociated scene-defined rank, overall behavioral acceptability, and categorical commitment within the same task, enabling detection of distinct neural signatures in SNr and FEF.

### SNr shows pre-target scene-related modulation and target-period ordinal rank coding

SNr neurons showed prominent scene-dependent modulation before target onset, whereas FEF modulation was comparatively weak, consistent with prior evidence that basal ganglia circuits integrate contextual and reward-related information (Mink, 1996; Kawagoe et al., 1998; Handel & Glimcher, 2000; Hikosaka et al., 2000; Lauwereyns et al., 2002; Sato & Hikosaka, 2002). Whether this modulation reflects scene context per se, the best available reward, or the scene value gap remains to be dissociated.

During target evaluation, multivariable modeling and behavioral model decomposition converged on the conclusion that SNr activity is dominated by ordinal rank, with comparatively little contribution from reward magnitude. The contributions of scene value gap and rejection history were not tested as individual neural predictors in the present study, though the composite Margin variable that incorporates all four factors performed substantially worse than Rank alone. This extends our recent finding that SNr codes binary good-versus-bad distinctions (Yoshida & Hikosaka, 2025a) by showing that the same pathway encodes ordinal rank across scenes with different reward amounts. The present results further indicate that SNr activity reflects a form of dimensionality reduction at the basal ganglia output stage, where multiple factors that jointly determine behavior to a dominant context-dependent binary distinction transmitted to downstream structures.

In the context of classic disinhibition models (Chevalier & Deniau, 1990; Mink, 1996), these findings suggest that the role of SNr extends beyond simply gating of actions. The data are consistent with a model in which SNr provides a context-dependent inhibitory bias organized primarily around ordinal rank, reducing inhibitory tone for higher-ranked offers (facilitating acceptance) while maintaining stronger inhibition for lower-ranked offers (biasing toward rejection) (Yoshida & Hikosaka, 2025a). This view is compatible with the focused selection and surround inhibition model proposed by Mink (1996, see also Redgrave et al., 1999; Nambu et al., 2002), in which the basal ganglia enable desired actions while suppressing competing alternatives. The ordinal nature of this signal, representing better-versus-worse within a local context rather than absolute reward magnitude, may also be relevant to models emphasizing opponent processing in the direct and indirect pathways (Kravitz et al., 2010; Gerfen & Surmeier, 2011; Collins & Frank, 2014). A binary rank distinction is compatible with the Go/NoGo architecture of the basal ganglia, although whether the rank signal observed in SNr reflects differential engagement of these opponent pathways remains to be tested through pathway-specific recordings or perturbations (e.g., Kravitz et al., 2010; Freeze et al., 2013).

### FEF reflects categorical commitment and directional selection

FEF neurons did not distinguish Accept Good from Accept Bad despite their difference in rank, but robustly separated acceptance from rejection and strongly encoded target direction. This dissociation is notable because Reject Bad trials involved active return saccades in which the monkey initiated a saccade toward the object before redirecting gaze, indicating that the FEF signal cannot be reduced to a simple movement-versus-no-movement contrast. Instead, the dominant FEF signal reflects a committed action state together with directional selection (Schall & Hanes, 1993; Hanes & Schall, 1996; Ding & Gold, 2012), with little sensitivity to rank, consistent with evidence that reward-context signals are weaker in FEF than in basal ganglia circuits (Roesch & Olson, 2003; Ding & Hikosaka, 2006; Fan et al., 2020). The multivariable analyses, together with the single-neuron trends, support this interpretation. Direction and Action were the dominant predictors of FEF activity, and Action-type and Direction-type neurons were the largest FEF subpopulations. Thus, compared with SNr, frontal cortical output emphasizes whether and where an action will be executed rather than the evaluative context in which the decision was made.

### Convergence on SC is consistent with a distributed circuit architecture

These findings are consistent with a circuit model in which SNr provides a rank-organized inhibitory bias and FEF provides categorical excitatory drive, converging on the SC immediately upstream of movement execution (Hikosaka et al., 2000; Cisek & Kalaska, 2010; Cisek, 2012; Schall, 2013). The SC is well positioned to combine these signals, because it receives both basal ganglia inhibitory input and frontal cortical drive (Hikosaka & Wurtz, 1983; Segraves & Goldberg, 1987; Sommer & Wurtz, 2000), and reward-related modulation has been documented in SC itself (Ikeda & Hikosaka, 2003, 2007). In this framework, categorical commitment does not arise in a single region. Rather, the present data are consistent with a model in which categorical commitment reflects the convergence of rank-organized inhibition from basal ganglia output and directionally specific excitatory drive from frontal cortex.

### Conceptual scope and relation to prior work

The results in Figure 5 correspond to a three-level framework. At the behavioral level, accept/reject decisions reflect the integration of multiple factors as captured by the composite Margin variable. At the level of basal ganglia output, SNr predominantly represents one component, ordinal rank, with substantially weaker sensitivity to the other factors. At the level of frontal cortical output, FEF represents the categorical outcome coupled with directional selection.

This SNr pattern is non-trivial. The composite acceptability measure, which integrates reward magnitude, scene value gap, and accumulated waiting cost along with rank, failed to outperform even a simple binary accept/reject variable as an account of SNr activity. In other words, basal ganglia output does not transmit a multi-factor value signal scaled to overall behavioral tendency. It transmits a more parsimonious, context-dependent ordinal distinction.

This framework clarifies that the dissociation between SNr and FEF is not simply between continuous and binary signals. It is between context-dependent evaluation, reflecting the relative standing of an offer in the current choice set, and categorical commitment, reflecting what action the animal will execute and where. This distinction has implications beyond the present task, because many naturalistic decisions require the brain to map context-dependent evaluations onto categorical actions through circuits that must accommodate varying choice sets.

Whereas prior comparisons between basal ganglia and frontal cortical contributions (Ding & Hikosaka, 2006; Ding & Gold, 2012; Ding, 2015; Fan et al., 2020) did not reach the output stage itself, our results fill this gap and extend our recent SNr findings (Yoshida & Hikosaka, 2025a) from a binary to an ordinal task. By directly comparing SNr with FEF, we demonstrate that distinct computations are distributed across parallel output pathways rather than collapsed into a common premotor signal. The conceptual scope should be defined precisely. Rank here refers to ordinal position within a scene-defined choice set, intentionally narrower than cardinal economic value. Our conclusions concern how a context-dependent evaluative signal is transformed into categorical commitment within a defined circuit, rather than claiming that SNr encodes a general, menu-invariant value variable (Zhang et al., 2021).

### Limitations and predictions

Several limitations should be noted. The data are correlational and do not establish causal necessity. We did not record simultaneously from SC, leaving unresolved how rank and commitment signals are combined at the single-neuron level. The upstream sources of the SNr rank signal remain to be specified through pathway-specific perturbations (Kravitz et al., 2010; Freeze et al., 2013). Regarding the pre-target period, the scene-period modulation was not decomposed into component variables, and visual scene identity and scene-defined choice context were not dissociated in the present design, leading us to interpret the pre-target signal conservatively as scene-related modulation. Furthermore, because neurons were selected for inclusion based on target-period responsiveness, the scene-period analyses reflect a population defined by target-related activity rather than scene-related activity. Whether similar scene-dependent modulation is present in a broader population of SNr and FEF neurons remains to be tested.

These limitations generate clear predictions. Simultaneous recordings across SNr, FEF, and SC should reveal how rank-organized inhibition and categorical drive are integrated at the single-neuron level, providing a direct test of the convergent circuit model proposed here. Causal perturbations of each node should differentially affect rank sensitivity, commitment, and movement direction. Such perturbations could be achieved through muscimol inactivation (Hikosaka & Wurtz, 1985a, 1985b; Dias & Segraves, 1999) or optogenetic manipulation (Kravitz et al., 2010; Amita et al., 2020; Katz et al., 2025).

In summary, by exploiting a task that partially dissociated evaluation from commitment, we found evidence that context-dependent ordinal rank and categorical commitment are encoded by distinct parallel output pathways upstream of the SC. This pattern suggests that the evaluation-to-commitment transformation is distributed across output pathways rather than localized within a single area.

## Acknowledgments

This work was supported by the National Eye Institute Intramural Research Program at the National Institutes of Health (ZIA EY000415 (O.H.) and ZIA EY000511 (R.J.K)). The contributions of the NIH authors were made as part of their official duties as NIH federal employees, are in compliance with agency policy requirements, and are considered Works of the United States Government. However, the findings and conclusions presented in this paper are those of the authors and do not necessarily reflect the views of the NIH or the U.S. Department of Health and Human Services. MRI scanning was conducted at the Neurophysiology Imaging Facility Core (National Institute of Mental Health, National Institute of Neurological Disorders and Stroke, and National Eye Institute). We thank D. Parker, H. Warnock, G. Tansey, K. Allen-Worthington, A.M. Nichols, D. Yochelson, J. Fuller-Deets, and M. Robinson for their technical assistance.

## Author contributions

Conceptualization, A.Y., O.H.; Methodology, A.Y., R.J.K; Software, A.Y.; Validation, A.Y.; Formal Analysis, A.Y.; Investigation, A.Y.; Resources, O.H.; Data Curation, A.Y.; Writing – Original Draft, A.Y.; Writing – Review & Editing, A.Y., R.J.K; Visualization, A.Y.; Supervision, O.H.; Funding Acquisition, O.H., R.J.K

## Methods

### Subjects

All experimental procedures adhered to the Public Health Service Policy on Laboratory Animal Care and were approved by the National Eye Institute Animal Care and Use Committee (NEI ACUC). Three male rhesus macaques (*Macaca mulatta*, 8–10 kg), designated as Monkeys Ch, Cr, and Sp, served as subjects. These animals were also used in our previous studies (Yoshida & Hikosaka, 2024, 2025a, 2025b; Yoshida et al., 2026). Surgical implantation of a plastic head holder and recording chambers was performed under isoflurane anesthesia and aseptic conditions. Detailed surgical and postoperative procedures have been described previously (Yoshida & Hikosaka, 2024). Following recovery, the monkeys were trained on oculomotor tasks. During experiments, the head was fixed and eye position was monitored at 1000 Hz using an infrared eye-tracking system (EyeLink 1000; SR Research, Ottawa, ON, Canada). Water intake was regulated according to an established protocol approved by the NEI ACUC. Experiments were conducted in a light- and sound-attenuated room with visual stimuli displayed via an LCD projector (PJ658; ViewSonic, Brea, CA, USA). Task execution and data acquisition were controlled by custom C++ software.

### Context-dependent sequential-offer choice task

#### Task overview and scene–object contingencies

Monkeys performed a context-dependent sequential-offer choice task in which the background scene defined the local choice set and thereby the scene-defined ordinal rank of each offered object. Five visual objects (Objects A–E) were associated with fixed reward amounts (Object A, 1.2 mL; B, 0.6 mL; C, 0.4 mL; D, 0.2 mL; E, 0 mL) (Fig. 1A).

The task comprised two condition types, choice scenes (Scenes 1–6) and forced-choice scenes (Scenes 7–10). In the choice scenes, each scene indicated a specific pair of available objects (Scene 1, A vs. B; Scene 2, A vs. C; Scene 3, B vs. C; Scene 4, B vs. D; Scene 5, C vs. D; and Scene 6, D vs. E) (Fig. 1B). Within each choice scene, one object was higher-ranked (“good”) and the other was lower-ranked (“bad”). This design ensured that the rank of an intermediate object varied with scene context (e.g., Object B was the lower-ranked option in Scene 1 but the higher-ranked option in Scenes 3 and 4). In the forced-choice scenes, only one object was available (Scene 7, A; Scene 8, B; Scene 9, C; and Scene 10, D). These forced-choice scenes served as a control condition.

Scenes 1–10 were presented in repeated pseudorandom blocks. In each block, all 10 scenes appeared once in a random order (without replacement), and blocks were repeated throughout the session.

### Trial and offer definitions

A trial was defined as the period from scene onset until the monkey accepted an offer and obtained the reward. Within each trial, one object was presented at a time. Each object presentation constituted an offer. After each rejection, a new offer was presented, and this sequence continued until acceptance. To prevent excessive trial duration, the number of offers per trial was set to a maximum of 15.

### Event sequence and response contingencies

Each trial started with presentation of a scene (1000 ms), followed by appearance of a central fixation point. After the monkey maintained central fixation for 700 ms, one object was presented peripherally at 15° eccentricity. Targets appeared at one of six predetermined locations spanning both contralateral and ipsilateral hemifields relative to the recording site.

In each choice scene, the identity of the offered object (higher-ranked vs. lower-ranked within that scene; hereafter good vs. bad) was drawn with equal probability (50% each) on every offer. Importantly, following a rejection, the identity of the next offer was re-sampled independently from the same scene-specific object pair with the same 50/50 probability. Thus, the same object could appear repeatedly across successive offers.

Monkeys could accept or reject each offer. Acceptance was registered when the monkey made a saccade to the object and maintained fixation on it for ≥400 ms, which triggered immediate delivery of the corresponding reward and terminated the trial. Rejection was registered when fixation on the object was not maintained for ≥400 ms. Rejection behavior was categorized into three strategies, return (saccade toward the object followed by gaze return to center within < 400 ms), stay (maintained central fixation for 1000 ms without making a saccade to the object), and other (saccade away from both center and object). Trials in which the monkey broke fixation before target onset were classified as fixation breaks and excluded from all behavioral and neuronal analyses. These are shown in Figure 1E for completeness but were not treated as a rejection strategy. The return strategy effectively shortened the inter-offer interval because the next offer could begin immediately after gaze returned to center, whereas the stay strategy required 1000 ms of central fixation, favoring use of return saccades as a rapid rejection strategy.

### MRI

Recording sites were localized using a 3-T MRI system (MAGNETOM Prisma; Siemens Healthineers, Erlangen, Germany) with T1-weighted and T2-weighted sequences (0.5-mm voxel size). SNr recording sites were targeted using quantitative susceptibility mapping (QSM), which improves SNr visualization in younger animals (Yoshida et al., 2021). FEF recording sites were targeted based on anatomical landmarks on T1-weighted images, specifically the anterior bank of the arcuate sulcus, where saccades could be evoked by low-threshold electrical stimulation (<50 μA, 100 Hz). Detailed MRI protocols and QSM reconstruction procedures have been described previously (Yoshida et al., 2021).

### Electrophysiology

Neuronal recordings began after the monkeys demonstrated stable performance. Single-unit activity in SNr and FEF was recorded using tungsten microelectrodes (1–9 MΩ; either Frederick Haer & Co., Bowdoin, ME, USA or Alpha Omega Engineering, Nof HaGalil, Israel). Electrodes were advanced through stainless steel guide tubes using a hydraulic micromanipulator (MO-973A, Narishige, Tokyo, Japan). Neural signals were amplified, bandpass filtered (0.3–10 kHz; A-M Systems, Sequim, WA, USA), and digitized at 40 kHz. Custom voltage-time window discrimination software was used to isolate individual neurons online by setting amplitude thresholds and inclusion/exclusion windows based on waveform morphology. Data collection commenced once stable isolation was achieved.

A neuron was included in the dataset if its mean firing rate during the target period (100–300 ms after target onset) significantly differed from the baseline firing rate during the 500 ms preceding target onset (paired *t*-test, *P* < 0.05). Given the broad spatial tuning often observed in SNr and FEF movement-related neurons, specific receptive-field mapping was not performed for each neuron. Instead, responses were analyzed for targets in both contralateral and ipsilateral hemifields.

### Data analysis

All behavioral and neurophysiological data were preprocessed using MATLAB 2022b (MathWorks, Natick, MA, USA). To compare normalized neuronal activity across different conditions, linear mixed-effects modeling of neuronal activity was performed to reduce type I errors and better represent the data structure (Yu et al., 2022) using R (version 4.4.2) with the *lme4 lmerTest*, *pbkrtest* packages (Bates et al., 2015; Kuznetsova et al., 2017; Halekoh & Højsgaard, 2014). Behavioral modeling, simulation recovery analyses, and condition-averaged SRT modeling were performed in Python 3.11 using statsmodels (Seabold & Perktold, 2010), with NumPy (Harris et al., 2020) and SciPy (Virtanen et al., 2020) for numerical computation. Data were managed using pandas (McKinney, 2010), and figures were generated using Matplotlib (Hunter, 2007).

In neural LMM analyses pooled across monkeys, monkey and neuron were included as random intercepts, (1|Monkey) + (1|Neuron). Analyses performed separately within a single monkey used a neuron random intercept, (1|Neuron). To confirm that pooled results were not driven by a single subject, all neural analyses were additionally performed separately for each monkey. The direction of effects was consistent across all three subjects (see Supplementary Tables). SRT analyses used a different random-effects structure (random intercept for recording session) as described below.

### Good-choice rate

For the good-choice rate (Fig. 1D), we computed for each scene and recording session the percentage of completed trials on which the monkey accepted the higher-ranked (good) object, defined as N(Accept Good) / [N(Accept Good) + N(Accept Bad)] × 100. One session corresponded to the behavioral data collected during isolation of a single neuron. Because trials almost always terminated in acceptance and Reject Good trials were rare (< 1% of good-offer trials), this measure reflects how often the monkey ultimately obtained the higher-ranked object in each scene. Distributions across sessions are shown as violin plots for each monkey.

### Saccade reaction time (SRT) Analysis

Saccade onset was defined as the time at which eye velocity exceeded 40°/s within 500 ms following target onset. SRT was defined as the latency from target onset to saccade initiation. For accept trials (Accept Good, Accept Bad), SRT corresponded to the initial saccade from central fixation toward the object. For reject trials, only return rejections (Reject Bad) were analyzed, and SRT was defined as the latency to the initial saccade from central fixation toward the object (i.e., the first component of the return sequence). Stay and other rejection types were excluded from SRT analyses because they did not involve task-directed saccade initiation toward the object.

To quantify the effects of decision outcome and scene context on SRTs, we fit separate linear mixed-effects models. Between-group models tested the effect of behavioral outcome (Accept Good, Accept Bad, Reject Bad) pooled across choice scenes, whereas within-outcome models tested the effect of scene identity within each behavioral outcome. Models included the relevant fixed effect and a random intercept for recording session to account for repeated measures. Statistical significance was determined using Type III Analysis of Variance (ANOVA) with Kenward–Roger’s method for degree-of-freedom approximation, followed by *post hoc* pairwise comparisons with Bonferroni correction. Effect sizes were estimated using partial eta squared (η²_p_).

### Neuronal data preprocessing and normalization

Neural data were aligned to scene onset and target onset. Spike density functions (SDFs) were generated by convolving spike trains with a Gaussian kernel (*σ* = 20 ms). To facilitate comparisons across neurons with different baseline firing rates, neuronal activity was z-score normalized following previously described approaches (Adler et al., 2012; Kaplan et al., 2020). For each neuron, the mean firing rate during the 500 ms preceding event onset (−500 to 0 ms relative to scene or target onset) was subtracted from the trial-averaged spike density function, and the result was divided by the standard deviation of that activity computed across the full peri-event time window. The pre-event baseline thus defined the subtracted mean, whereas the normalizing standard deviation was computed from the entire peri-event response rather than from the baseline period.

### Analysis of scene- and target-period modulation (LMM)

To quantify neural modulation during specific task epochs, we performed LMM analyses on the mean normalized firing rates in a fixed time window (100–300 ms after scene or target onset), with neuron treated as the random unit as described above.

- **Scene Period:** the model included scene identity (Scenes 1–6) as a fixed effect and a random-intercept structure of (1|Monkey) + (1|Neuron).
- **Target Period:** the model included behavioral outcome (Accept Good, Accept Bad, Reject Bad) as a fixed effect and a random-intercept structure of (1|Monkey) + (1|Neuron).

Statistical significance was assessed using Type III ANOVA with Kenward–Roger’s method. Effect sizes were estimated using partial eta squared (η²_p_). For single-monkey analyses summarized in the supplementary tables, significance was additionally assessed using permutation ANOVA (10,000 randomizations).

### Quantification of task-variable contributions (Multivariable LMM)

To disentangle the independent contributions of correlated task variables to neuronal activity, we fit a multivariable LMM to firing rates in the target-period window (100–400 ms after target onset) from the choice task (Scenes 1–6). The model included the following fixed effects:

1. **Action:** Coded as 1 for “Accept” and −1 for “Reject”. Action here refers to the monkey’s accept/reject commitment, not merely the presence or absence of a saccade, because Reject Bad (return) trials also involved an initial saccade toward the object.
2. **Rank:** Coded as 1 for higher-ranked and −1 for lower-ranked within the current scene.
3. **Reward amount:** a continuous variable representing reward magnitude, standardized (z-scored).
4. **Direction:** Coded as 1 for contralateral and −1 for ipsilateral targets.
5. **SRT:** log-transformed and z-scored within each monkey.

Trials with missing SRT values (e.g., due to blinks or eye-tracker loss) were excluded from SRT-controlled analyses.

The model specification was as follows.

*FiringRate ∼ Action + Rank + RewardAmount + Direction + SRT + (1|Monkey) + (1|Neuron)*, where (1|Monkey) and (1|Neuron) represent random intercepts for monkey and neuron, respectively. Monkey was included as an additional random intercept to account for between-animal variability. Monkey-level consistency was further verified by repeating the analysis separately for each monkey.

### Sliding-window analysis

To examine the temporal dynamics of task-variable encoding, we applied the same multivariable LMM using a sliding-window approach. A 200-ms analysis window was shifted in 10-ms steps from −500 ms to +500 ms relative to target onset. At each time step, the coefficients (*β*) for each fixed effect were estimated to construct the time course of variable encoding. The sign-inversion applied to SNr coefficients in Fig. 4B affects visualization only and is described in the Fig. 4 legend. All statistical analyses used the original coefficient signs.

### Neuron-type classification

To determine the primary encoding variable of individual neurons, we estimated time-resolved regression coefficients for each neuron by fitting a multiple linear regression (Gaussian GLM) at each time point, using standardized spike density as the dependent variable and Action, Rank, RewardAmount, Direction, and SRT as predictors. This classification was intended as a descriptive summary of single-neuron response profiles rather than a formal hypothesis test, for the reasons detailed at the end of this section. We then classified neurons using the mean *β* values obtained from this time-resolved single-neuron regression in a fixed window (100–400 ms after target onset). We used a slightly broader window for classification to capture neurons with later-onset effects that might not peak within the narrower 100–300 ms evaluation window used for the main coefficient summaries.

Classification criteria were defined based on the known dominant response polarities of each region. For SNr neurons (typically inhibitory or disinhibitory), we identified the variable with the strongest negative modulation (minimum *β*). For FEF neurons (typically excitatory), we identified the variable with the strongest positive modulation (maximum *β*).

A neuron was classified into a specific type (Action, Rank, Reward amount, Direction, or SRT) only if the magnitude of its dominant *β* was significantly larger than all four other *β* values. For each pairwise comparison, we performed a paired *t*-test across time bins within the classification window (100–400 ms after target onset), treating the *β* difference at each time bin as a paired observation. Statistical significance was assessed with Bonferroni correction for multiple comparisons (*α* = 0.05 / 10 comparisons = 0.005). Because classification was based on temporally smoothed time-resolved *β* traces, adjacent time bins were not statistically independent. Due to this dependence structure, the procedure is interpreted as a descriptive summary of each neuron’s dominant response profile rather than as a formal hypothesis test, and the resulting subtype proportions should not be taken as exact inferential estimates. Neurons that did not meet this criterion were categorized as “Unclassified.”

### Computational modeling

To complement the multivariable regression analysis (Fig. 4), we developed computational models to identify which task factors best predicted accept/reject decisions, extract a continuous acceptability measure (Margin), and test whether this variable or its components better explained SNr versus FEF activity.

#### Behavioral model comparison

We fit logistic regression models to offer-level choice data from the choice task (Scenes 1–6), predicting the binary accept/reject outcome on each offer. Behavioral models used all offers (including rare Reject Good) to maximize statistical power, whereas neural and SRT analyses excluded Reject Good trials due to insufficient trial counts per neuron. Five models of increasing complexity were compared:

- M0: reward amount (the reward magnitude associated with the offered object)
- M1: reward amount + scene identity (separate intercept for each scene)
- M2: reward amount + scene value gap (difference between the good and bad reward amounts in the current scene)
- M3: M2 + number of prior rejections within the trial (NPrevRejects)
- M4: M3 + ordinal rank (1 = higher-ranked within the current scene; 0 = lower-ranked)

Models were fit using generalized linear models (binomial family, logit link). To evaluate generalization performance while respecting the non-independence of offers within a recording session, we used session-level leave-one-out cross-validation. For each of the approximately 190 sessions, the model was fit on all other sessions, and the log-likelihood of the held-out session’s choices was computed. The total cross-validated log-likelihood across all held-out sessions was used as the primary model selection criterion. AIC and BIC from full-data fits were reported as secondary metrics.

#### Margin and Action

From the best behavioral model, we extracted Margin, defined as the linear predictor on the logit scale, representing the signed distance from the decision boundary. For comparison, we defined Action as a binary variable recording the monkey’s actual choice (accept = 1, reject = 0).

Monkey-specific Margin values were also computed by fitting M4 separately for each monkey and applying that monkey’s parameters to its own offers. This removed between-monkey differences in average Margin that could confound the neuronal analysis.

#### Simulation recovery

Before comparing Margin and Action models on real neural data, we verified that the analysis could reliably distinguish graded from categorical neural signals. We generated synthetic firing-rate data under two scenarios, (1) a graded scenario in which firing rates were a linear function of Margin plus noise, and (2) a categorical scenario in which firing rates were a linear function of Action plus noise. For each scenario, we fit both the Margin model and the Action model and asked whether the correct model was selected by ΔAIC (see below). Parameters for the simulation (effect sizes, residual noise, number of neurons, number of trials per neuron, and directional tuning) were estimated separately from the real SNr and FEF data. We ran 500 simulations per scenario and additionally tested robustness across a grid of effect-size and noise multipliers (0.25–2.0× each). Recovery rates exceeded 80% in all conditions tested.

#### Neural model comparison

To test which variables best explained target-period neuronal activity (100–300 ms after target onset), we fit linear mixed-effects models to z-score-normalized firing rates from the three main behavioral conditions (Accept Good, Accept Bad, Reject Bad return). The following models were compared for each brain area:

- Baseline: FR ∼ Direction + (1|Monkey) + (1|Neuron)
- Action model: FR ∼ Action + Direction + (1|Monkey) + (1|Neuron)
- Margin model: FR ∼ Margin + Direction + (1|Monkey) + (1|Neuron)
- Rank model: FR ∼ Rank + Direction + (1|Monkey) + (1|Neuron)
- Reward amount model: FR ∼ Reward amount + Direction + (1|Monkey) + (1|Neuron)

where Direction encodes the target location as contralateral (+1) or ipsilateral (−1) relative to the recording hemisphere, and (1|Monkey) and (1|Neuron) represent random intercepts for monkey and neuron. Combined models were also fit, including Rank + Action + Direction and a Full model (Action + Rank + Reward amount + Direction). Scene value gap and number of prior rejections were not tested as individual neural predictors because the primary question was whether the dominant SNr signal reflects ordinal rank or reward amount. These two variables were included only as components of the composite Margin variable.

All LMMs used for model comparison were fit with maximum likelihood (ML; REML = FALSE), as REML-based likelihoods are not comparable across models with different fixed-effects structures.

Model comparison used ΔAIC = AIC(Action model) − AIC(test model), such that positive values indicate the test model explains neuronal activity better than the Action model. To assess the dissociation at the single-neuron level, we fit neuron-wise ordinary least squares component-decomposition models in which the full model included Rank, Action, Reward amount, monkey-specific Margin, and Direction. For each task variable, partial R² was computed as 1 − (residual sum of squares of the full model) / (residual sum of squares of the reduced model omitting that variable). For each neuron we then formed a signed index, Δ partial R² = partial R²(Rank) − partial R²(Action), and compared this index between SNr and FEF using a one-sided Mann–Whitney *U* test (testing whether SNr exceeded FEF). Effect sizes were summarized by Cohen’s *d* and the common-language effect size, and the median between-area difference was bracketed by a bootstrap 95% confidence interval.

For combined models containing both Rank (or Margin) and Action, we computed variance inflation factors (VIF) to assess collinearity. Partial regression coefficients from these models were interpreted only when VIF < 5.

Neuron-wise z-scoring was performed by subtracting each neuron’s mean firing rate and dividing by its standard deviation across all included trials. Neurons with zero variance were excluded.

All neuronal analyses were performed for the pooled population and for each monkey separately.

### Use of generative AI and AI-assisted technologies

During the preparation of this work, the authors used Claude Opus 4.8 (Anthropic) and ChatGPT 5.5 (OpenAI) to check for typographical and grammatical errors and to improve the readability of author-written text. All original analysis scripts were written by the authors. These generative AI tools were used only to review and suggest improvements to the authors’ own code. After using these tools, the authors reviewed and edited the content and code as needed and take full responsibility for the content of the published article.

## Data availability

The data supporting the findings of this study are provided with this paper as a Source Data file (numerical data underlying all figures, comprising behavioral metrics and normalized neuronal firing rates). The processed behavioral and electrophysiological datasets have been deposited in Zenodo under accession 10.5281/zenodo.20707929, and will be made publicly available upon publication of this article. No restrictions apply to data availability, and this study did not use clinical or third-party datasets.

## Code availability

Custom code used for behavioral modeling, linear mixed-effects analyses, simulation-recovery analyses, and figure generation (R 4.4.2 and Python 3.11) is available from the corresponding author (A.Y.) upon reasonable request.

## References

Gold, J. I. & Shadlen, M. N. The neural basis of decision making. Annu. Rev. Neurosci. 30, 535–574 (2007).

Thura, D. & Cisek, P. Deliberation and commitment in the premotor and primary motor cortex during dynamic decision making. Neuron 81, 1401–1416 (2014).

Chen, X. & Stuphorn, V. Sequential selection of economic good and action in medial frontal cortex of macaques during value-based decisions. eLife 4, e09418 (2015).

Zhang, B. et al. Transforming absolute value to categorical choice in primate superior colliculus during value-based decision making. Nat. Commun. 12, 3410 (2021).

Hikosaka, O., Takikawa, Y. & Kawagoe, R. Role of the basal ganglia in the control of purposive saccadic eye movements. Physiol. Rev. 80, 953–978 (2000).

Hanes, D. P. & Wurtz, R. H. Interaction of the frontal eye field and superior colliculus for saccade generation. J. Neurophysiol. 85, 804–815 (2001).

Jayaraman, A., Batton, R. R. III & Carpenter, M. B. Nigrotectal projections in the monkey: an autoradiographic study. Brain Res. 135, 147–152 (1977).

Hikosaka, O. & Wurtz, R. H. Visual and oculomotor functions of monkey substantia nigra pars reticulata. IV. Relation of substantia nigra to superior colliculus. J. Neurophysiol. 49, 1285–1305 (1983).

May, P. J. & Hall, W. C. Relationships between the nigrotectal pathway and the cells of origin of the predorsal bundle. J. Comp. Neurol. 226, 337–376 (1984).

Liu, P. & Basso, M. A. Substantia nigra stimulation influences monkey superior colliculus neuronal activity bilaterally. J. Neurophysiol. 100, 1098–1112 (2008).

Künzle, H., Akert, K. & Wurtz, R. H. Projection of area 8 (frontal eye field) to superior colliculus in the monkey. An autoradiographic study. Brain Res. 117, 487–492 (1976).

Helminski, J. O. & Segraves, M. A. Macaque frontal eye field input to saccade-related neurons in the superior colliculus. J. Neurophysiol. 90, 1046–1062 (2003).

Segraves, M. A. & Goldberg, M. E. Functional properties of corticotectal neurons in the monkey’s frontal eye field. J. Neurophysiol. 58, 1387–1419 (1987).

Stanton, G. B., Goldberg, M. E. & Bruce, C. J. Frontal eye field efferents in the macaque monkey: II. Topology of terminal fields in midbrain and pons. J. Comp. Neurol. 271, 493–506 (1988).

Sommer, M. A. & Wurtz, R. H. Composition and topographic organization of signals sent from the frontal eye field to the superior colliculus. J. Neurophysiol. 83, 1979–2001 (2000).

Ikeda, T. & Hikosaka, O. Reward-dependent gain and bias of visual responses in primate superior colliculus. Neuron 39, 693–700 (2003).

Ikeda, T. & Hikosaka, O. Positive and negative modulation of motor response in primate superior colliculus by reward expectation. J. Neurophysiol. 98, 3163–3170 (2007).

Herman, J. P., Katz, L. N. & Krauzlis, R. J. Midbrain activity can explain perceptual decisions during an attention task. Nat. Neurosci. 21, 1651–1655 (2018).

Basso, M. A. & Wurtz, R. H. Modulation of neuronal activity in superior colliculus by changes in target probability. J. Neurosci. 18, 7519–7534 (1998).

Everling, S., Dorris, M. C., Klein, R. M. & Munoz, D. P. Role of primate superior colliculus in preparation and execution of anti-saccades and pro-saccades. J. Neurosci. 19, 2740–2754 (1999).

Horwitz, G. D., Batista, A. P. & Newsome, W. T. Representation of an abstract perceptual decision in macaque superior colliculus. J. Neurophysiol. 91, 2281–2296 (2004).

Horwitz, G. D. & Newsome, W. T. Separate signals for target selection and movement specification in the superior colliculus. Science 284, 1158–1161 (1999).

Chevalier, G. & Deniau, J. M. Disinhibition as a basic process in the expression of striatal functions. Trends Neurosci. 13, 277–280 (1990).

Mink, J. W. The basal ganglia: focused selection and inhibition of competing motor programs. Prog. Neurobiol. 50, 381–425 (1996).

Handel, A. & Glimcher, P. W. Contextual modulation of substantia nigra pars reticulata neurons. J. Neurophysiol. 83, 3042–3048 (2000).

Sato, M. & Hikosaka, O. Role of primate substantia nigra pars reticulata in reward-oriented saccadic eye movement. J. Neurosci. 22, 2363–2373 (2002).

Yasuda, M. & Hikosaka, O. Functional territories in primate substantia nigra pars reticulata separately signaling stable and flexible values. J. Neurophysiol. 113, 1681–1696 (2015).

Yasuda, M., Yamamoto, S. & Hikosaka, O. Robust representation of stable object values in the oculomotor basal ganglia. J. Neurosci. 32, 16917–16932 (2012).

Ding, L. & Hikosaka, O. Comparison of reward modulation in the frontal eye field and caudate of the macaque. J. Neurosci. 26, 6695–6703 (2006).

Fan, Y., Gold, J. I. & Ding, L. Frontal eye field and caudate neurons make different contributions to reward-biased perceptual decisions. eLife 9, e60535 (2020).

Doi, T., Fan, Y., Gold, J. I. & Ding, L. The caudate nucleus contributes causally to decisions that balance reward and uncertain visual information. eLife 9, e56694 (2020).

Schall, J. D. & Hanes, D. P. Neural basis of saccade target selection in frontal eye field during visual search. Nature 366, 467–469 (1993).

Hanes, D. P. & Schall, J. D. Neural control of voluntary movement initiation. Science 274, 427–430 (1996).

Cisek, P. & Kalaska, J. F. Neural mechanisms for interacting with a world full of action choices. Annu. Rev. Neurosci. 33, 269–298 (2010).

Tremblay, L. & Schultz, W. Relative reward preference in primate orbitofrontal cortex. Nature 398, 704–708 (1999).

Yoshida, A. & Hikosaka, O. Involvement of neurons in the nonhuman primate anterior striatum in proactive inhibition. J. Neurosci. 44, e0866242024 (2024).

Yoshida, A. & Hikosaka, O. Contribution of glutamatergic projections to neurons in the nonhuman primate substantia nigra pars reticulata for reactive inhibition. Proc. Natl Acad. Sci. USA 122, e2427032122 (2025a).

Yoshida, A. & Hikosaka, O. Excitatory drive to the globus pallidus external segment facilitates action initiation in non-human primates. Curr. Biol. 35, 1–16 (2025b).

Yoshida, A., Krauzlis, R. J. & Hikosaka, O. Beyond the Brake: the Subthalamic Nucleus Predominantly Facilitates Action in Non-human Primates. Prog. Neurobiol. 262, 102921 (2026).

Cisek, P. Making decisions through a distributed consensus. Curr. Opin. Neurobiol. 22, 927–936 (2012).

Schall, J. D. Macrocircuits: decision networks. Curr. Opin. Neurobiol. 23, 269–274 (2013).

Kawagoe, R., Takikawa, Y. & Hikosaka, O. Expectation of reward modulates cognitive signals in the basal ganglia. Nat. Neurosci. 1, 411–416 (1998).

Lauwereyns, J. et al. Feature-based anticipation of cues that predict reward in monkey caudate nucleus. Neuron 33, 463–473 (2002).

Redgrave, P., Prescott, T. J. & Gurney, K. The basal ganglia: a vertebrate solution to the selection problem? Neuroscience 89, 1009–1023 (1999).

Nambu, A., Tokuno, H. & Takada, M. Functional significance of the cortico-subthalamo-pallidal ‘hyperdirect’ pathway. Neurosci. Res. 43, 111–117 (2002).

Kravitz, A. V. et al. Regulation of parkinsonian motor behaviours by optogenetic control of basal ganglia circuitry. Nature 466, 622–626 (2010).

Gerfen, C. R. & Surmeier, D. J. Modulation of striatal projection systems by dopamine. Annu. Rev. Neurosci. 34, 441–466 (2011).

Collins, A. G. E. & Frank, M. J. Opponent actor learning (OpAL): modeling interactive effects of striatal dopamine on reinforcement learning and choice incentive. Psychol. Rev. 121, 337–366 (2014).

Freeze, B. S., Kravitz, A. V., Hammack, N., Berke, J. D. & Kreitzer, A. C. Control of basal ganglia output by direct and indirect pathway projection neurons. J. Neurosci. 33, 18531–18539 (2013).

Ding, L. & Gold, J. I. Neural correlates of perceptual decision making before, during, and after decision commitment in monkey frontal eye field. Cereb. Cortex 22, 1052–1067 (2012).

Roesch, M. R. & Olson, C. R. Impact of expected reward on neuronal activity in prefrontal cortex, frontal and supplementary eye fields and premotor cortex. J. Neurophysiol. 90, 1766–1789 (2003).

Ding, L. Distinct dynamics of ramping activity in the frontal cortex and caudate nucleus in monkeys. J. Neurophysiol. 114, 1850–1861 (2015).

Hikosaka, O. & Wurtz, R. H. Modification of saccadic eye movements by GABA-related substances. I. Effect of muscimol and bicuculline in monkey superior colliculus. J. Neurophysiol. 53, 266–291 (1985a).

Hikosaka, O. & Wurtz, R. H. Modification of saccadic eye movements by GABA-related substances. II. Effects of muscimol in monkey substantia nigra pars reticulata. J. Neurophysiol. 53, 292–308 (1985b).

Dias, E. C. & Segraves, M. A. Muscimol-induced inactivation of monkey frontal eye field: effects on visually and memory-guided saccades. J. Neurophysiol. 81, 2191–2214 (1999).

Amita, H., Kim, H. F., Inoue, K., Takada, M. & Hikosaka, O. Optogenetic manipulation of a value-coding pathway from the primate caudate tail facilitates saccadic gaze shift. Nat. Commun. 11, 1876 (2020).

Katz, L. N., Bohlen, M. O., Yu, G., Mejias-Aponte, C., Sommer, M. A. & Krauzlis, R. J. Optogenetic manipulation of covert attention in the nonhuman primate. J. Cogn. Neurosci. 37, 266–285 (2025).

Yoshida, A., Ye, F. Q., Yu, D. K., Leopold, D. A. & Hikosaka, O. Visualization of iron-rich subcortical structures in non-human primates in vivo by quantitative susceptibility mapping at 3T MRI. NeuroImage 241, 118429 (2021).

Yu, Z., Wan, Y., and Yu, X. Beyond t test and ANOVA: applications of mixed-effects models for more rigorous statistical analysis in neuroscience research. Neuron 110, 21–35 (2022).

Bates, D., Mächler, M., Bolker, B., and Walker, S. Fitting linear mixed-effects models using lme4. J. Stat. Softw. 67, 1–48 (2015).

Kuznetsova, A., Brockhoff, P. B., and Christensen, R. H. B. Imer Test Package: Tests in Linear Mixed Effects Models. J. Stat. Softw. 82, 1–26 (2017).

Halekoh, U., and Højsgaard, S. A Kenward-Roger approximation and parametric bootstrap methods for tests in linear mixed models – the R package pbkrtest. J. Stat. Softw. 59, 1–30 (2014).

Harris, C. R. et al. Array programming with NumPy. Nature 585, 357–362 (2020).

Hunter, J. D. Matplotlib: A 2D graphics environment. Comput. Sci. Eng. 9, 90–95 (2007).

McKinney, W. Data structures for statistical computing in Python. Proc. 9th Python Sci. Conf. 56–61 (2010).

Seabold, S. & Perktold, J. Statsmodels: Econometric and statistical modeling with Python. Proc. 9th Python Sci. Conf. 92–96 (2010).

Virtanen, P. et al. SciPy 1.0: fundamental algorithms for scientific computing in Python. Nat. Methods 17, 261–272 (2020).

Adler, A., Katabi, S., Finkes, I., Israel, Z., Prut, Y. & Bergman, H. Temporal convergence of dynamic cell assemblies in the striato-pallidal network. J. Neurosci. 32, 2473–2484 (2012).

Kaplan, A., Mizrahi-Kliger, A. D., Israel, Z., Adler, A. & Bergman, H. Dissociable roles of ventral pallidum neurons in the basal ganglia reinforcement learning network. Nat. Neurosci. 23, 556–564 (2020).

