## Supplemental Table 1 for "Parallel basal ganglia and frontal cortical outputs differentially encode context-dependent evaluation and categorical commitment during choice"

**Table S1. Descriptive statistics of saccade reaction times across behavioral outcomes and scenes.**

Summary of saccade reaction times (SRTs) for each monkey (Monkey Ch, Monkey Cr, Monkey Sp) corresponding to the behavioral results in Figure 1F, 1G, and Figure S1C. Data are categorized by Analysis type: “Within” rows break down SRTs by specific Scenes (1–6) for each behavioral outcome, indicated by the suffix _AG, _AB, or _RB in the Analysis column; “Between_Groups” rows aggregate data across all choice scenes; “Forced-choice” rows correspond to control scenes (7–10) where only a single object was presented.

Condition: specifies the Scene ID (1–10) or the aggregated behavioral outcome.

n: number of trials. Mean SRT (ms): mean saccade reaction time in milliseconds. SD (ms): standard deviation. 95% CI (ms): 95% confidence interval of the mean.

Abbreviations: AG, Accept Good; AB, Accept Bad; RB, Reject Bad (return).

Note: For Reject Bad trials classified as “return,” SRT is defined as the latency of the initial saccade from the central fixation point toward the object, not the subsequent return saccade back to the center.

| **Monkey** | **Analysis** | **Condition** | **n** | **Mean SRT (ms)** | **SD (ms)** | **95% CI (ms)** |
| --- | --- | --- | --- | --- | --- | --- |
| Monkey Ch | Within_AG | Scene 1 | 1034 | 200.09 | 24.39 | [198.60, 201.57] |
| Monkey Ch | Within_AG | Scene 2 | 1053 | 197.79 | 21.95 | [196.46, 199.12] |
| Monkey Ch | Within_AG | Scene 3 | 667 | 197.04 | 27.97 | [194.91, 199.17] |
| Monkey Ch | Within_AG | Scene 4 | 1060 | 200.89 | 23.73 | [199.46, 202.32] |
| Monkey Ch | Within_AG | Scene 5 | 718 | 198.25 | 22.80 | [196.58, 199.92] |
| Monkey Ch | Within_AG | Scene 6 | 1008 | 194.97 | 21.59 | [193.64, 196.31] |
| Monkey Ch | Within_AB | Scene 3 | 675 | 206.97 | 24.74 | [205.11, 208.84] |
| Monkey Ch | Within_AB | Scene 5 | 669 | 206.38 | 27.98 | [204.26, 208.51] |
| Monkey Ch | Within_RB | Scene 1 | 1064 | 243.65 | 40.83 | [241.19, 246.10] |
| Monkey Ch | Within_RB | Scene 2 | 1125 | 237.97 | 45.40 | [235.31, 240.63] |
| Monkey Ch | Within_RB | Scene 4 | 995 | 247.05 | 46.04 | [244.19, 249.92] |
| Monkey Ch | Within_RB | Scene 6 | 974 | 244.32 | 45.54 | [241.45, 247.18] |
| Monkey Ch | Between_Groups | Accept Good | 5540 | 198.27 | 23.67 | [197.64, 198.89] |
| Monkey Ch | Between_Groups | Accept Bad | 1344 | 206.68 | 26.40 | [205.27, 208.09] |
| Monkey Ch | Between_Groups | Reject Bad | 4158 | 243.08 | 44.58 | [241.73, 244.44] |
| Monkey Ch | Forced choice | Scene 7 | 1061 | 192.80 | 26.50 | [191.21, 194.40] |
| Monkey Ch | Forced choice | Scene 8 | 1064 | 190.80 | 30.29 | [188.98, 192.62] |
| Monkey Ch | Forced choice | Scene 9 | 1047 | 191.78 | 26.15 | [190.19, 193.36] |
| Monkey Ch | Forced choice | Scene 10 | 1053 | 192.57 | 24.00 | [191.12, 194.02] |
| Monkey Cr | Within_AG | Scene 1 | 1699 | 191.06 | 27.63 | [189.74, 192.37] |
| Monkey Cr | Within_AG | Scene 2 | 1661 | 187.81 | 27.55 | [186.49, 189.14] |
| Monkey Cr | Within_AG | Scene 3 | 1154 | 184.36 | 26.89 | [182.81, 185.91] |
| Monkey Cr | Within_AG | Scene 4 | 1631 | 188.15 | 26.48 | [186.86, 189.43] |
| Monkey Cr | Within_AG | Scene 5 | 999 | 182.88 | 24.88 | [181.33, 184.42] |
| Monkey Cr | Within_AG | Scene 6 | 1574 | 185.88 | 25.96 | [184.60, 187.17] |
| Monkey Cr | Within_AB | Scene 3 | 744 | 200.56 | 26.95 | [198.62, 202.50] |
| Monkey Cr | Within_AB | Scene 5 | 923 | 197.05 | 27.66 | [195.27, 198.84] |
| Monkey Cr | Within_RB | Scene 1 | 1695 | 228.19 | 56.28 | [225.51, 230.87] |
| Monkey Cr | Within_RB | Scene 2 | 1643 | 228.59 | 52.32 | [226.06, 231.12] |
| Monkey Cr | Within_RB | Scene 4 | 1634 | 231.15 | 53.62 | [228.55, 233.75] |
| Monkey Cr | Within_RB | Scene 6 | 1157 | 257.34 | 65.18 | [253.58, 261.10] |
| Monkey Cr | Between_Groups | Accept Good | 8718 | 187.14 | 26.82 | [186.57, 187.70] |
| Monkey Cr | Between_Groups | Accept Bad | 1667 | 198.62 | 27.40 | [197.30, 199.94] |
| Monkey Cr | Between_Groups | Reject Bad | 6129 | 234.59 | 57.43 | [233.15, 236.03] |
| Monkey Cr | Forced choice | Scene 7 | 1693 | 180.82 | 31.83 | [179.31, 182.34] |
| Monkey Cr | Forced choice | Scene 8 | 1676 | 178.55 | 29.97 | [177.11, 179.99] |
| Monkey Cr | Forced choice | Scene 9 | 1661 | 182.62 | 32.38 | [181.06, 184.18] |
| Monkey Cr | Forced choice | Scene 10 | 1543 | 186.74 | 31.59 | [185.16, 188.32] |
| Monkey Sp | Within_AG | Scene 1 | 358 | 202.89 | 26.77 | [200.10, 205.67] |
| Monkey Sp | Within_AG | Scene 2 | 649 | 209.79 | 29.76 | [207.49, 212.08] |
| Monkey Sp | Within_AG | Scene 3 | 655 | 206.22 | 23.83 | [204.39, 208.04] |
| Monkey Sp | Within_AG | Scene 4 | 659 | 202.85 | 24.70 | [200.96, 204.74] |
| Monkey Sp | Within_AG | Scene 5 | 647 | 197.06 | 25.46 | [195.09, 199.02] |
| Monkey Sp | Within_AG | Scene 6 | 593 | 204.83 | 29.46 | [202.46, 207.21] |
| Monkey Sp | Within_AB | Scene 1 | 334 | 215.99 | 33.96 | [212.34, 219.65] |
| Monkey Sp | Within_RB | Scene 2 | 423 | 328.10 | 61.08 | [322.26, 333.94] |
| Monkey Sp | Within_RB | Scene 3 | 514 | 310.42 | 54.68 | [305.69, 315.16] |
| Monkey Sp | Within_RB | Scene 4 | 509 | 315.30 | 48.51 | [311.08, 319.53] |
| Monkey Sp | Within_RB | Scene 5 | 485 | 285.97 | 68.32 | [279.88, 292.07] |
| Monkey Sp | Within_RB | Scene 6 | 346 | 324.40 | 69.27 | [317.08, 331.73] |
| Monkey Sp | Between_Groups | Accept Good | 3561 | 204.01 | 26.99 | [203.13, 204.90] |
| Monkey Sp | Between_Groups | Accept Bad | 334 | 215.99 | 33.96 | [212.34, 219.65] |
| Monkey Sp | Between_Groups | Reject Bad | 2277 | 311.71 | 61.88 | [309.17, 314.26] |
| Monkey Sp | Forced choice | Scene 7 | 650 | 189.77 | 40.68 | [186.64, 192.90] |
| Monkey Sp | Forced choice | Scene 8 | 652 | 187.79 | 35.78 | [185.04, 190.54] |
| Monkey Sp | Forced choice | Scene 9 | 634 | 186.05 | 40.39 | [182.90, 189.20] |
| Monkey Sp | Forced choice | Scene 10 | 610 | 203.57 | 49.91 | [199.60, 207.54] |
