## Supplemental Table 2 for "Parallel basal ganglia and frontal cortical outputs differentially encode context-dependent evaluation and categorical commitment during choice"

**Table S2. Statistical results of linear mixed-effects analysis on saccade reaction times.**

Results of Type III analysis of variance (ANOVA) on SRTs (in milliseconds) for each monkey (related to Figure 1F, 1G, and Figure S1C).

The Analysis column specifies the fixed effect tested in each model:

Within_XX (e.g., Within_AG): tests the effect of Scene identity (Scenes 1–6, or 7–10 for Within_Forced) on SRTs within a specific behavioral outcome.

Between_Groups: tests the effect of behavioral outcome (Accept Good, Accept Bad, Reject Bad) on SRTs pooled across all choice scenes.

Statistics: F, F-statistic from the ANOVA; df (num, den), numerator and denominator degrees of freedom (Satterthwaite); P, P-value (values below 0.001 are shown as “< 0.001”); Partial η², partial eta squared (η²p) as a measure of effect size, computed as F·df_num / (F·df_num + df_den); Effect size, qualitative magnitude of the effect.

Abbreviations: AG, Accept Good; AB, Accept Bad; RB, Reject Bad (return); Forced, forced-choice trials (Scenes 7–10).

Note: Between_Groups (behavioral outcome) yielded large effect sizes in all monkeys (Partial η² = 0.245–0.614), whereas Within-scene effects were comparatively small (Partial η² ≤ 0.077), indicating that SRT variation is primarily driven by decision outcome rather than scene context.

| **Monkey** | **Analysis** | **F** | **df (num, den)** | **P** | **Partial η²** | **Effect size** |
| --- | --- | --- | --- | --- | --- | --- |
| Monkey Ch | Within_AG | 8.21 | 5, 4730.7 | 9.8 × 10⁻⁸ | 0.009 | Negligible |
| Monkey Ch | Within_AB | 0.17 | 1, 693.5 | 0.680 | 0.000 | Negligible |
| Monkey Ch | Within_RB | 8.07 | 3, 3149.6 | 2.3 × 10⁻⁵ | 0.008 | Negligible |
| Monkey Ch | Between_Groups | 2220.52 | 2, 10415.4 | < 0.001 | 0.299 | Large |
| Monkey Ch | Within_Forced | 1.21 | 3, 3211.4 | 0.305 | 0.001 | Negligible |
| Monkey Cr | Within_AG | 16.21 | 5, 7312.7 | 6.3 × 10⁻¹⁶ | 0.011 | Small |
| Monkey Cr | Within_AB | 6.87 | 1, 908.3 | 0.009 | 0.008 | Negligible |
| Monkey Cr | Within_RB | 79.47 | 3, 4714.2 | 3.8 × 10⁻⁵⁰ | 0.048 | Small |
| Monkey Cr | Between_Groups | 2522.21 | 2, 15533.4 | < 0.001 | 0.245 | Large |
| Monkey Cr | Within_Forced | 22.85 | 3, 4930.5 | 1.1 × 10⁻¹⁴ | 0.014 | Small |
| Monkey Sp | Within_AG | 20.51 | 5, 2952.1 | 3.5 × 10⁻²⁰ | 0.034 | Small |
| Monkey Sp | Within_RB | 38.26 | 4, 1836.7 | 8.6 × 10⁻³¹ | 0.077 | Medium |
| Monkey Sp | Between_Groups | 4636.67 | 2, 5828.8 | < 0.001 | 0.614 | Large |
| Monkey Sp | Within_Forced | 23.94 | 3, 1917.2 | 3.3 × 10⁻¹⁵ | 0.036 | Small |
