## Supplemental Table 3 for "Parallel basal ganglia and frontal cortical outputs differentially encode context-dependent evaluation and categorical commitment during choice"

**Table S3. Post-hoc pairwise comparisons of saccade reaction times.**

Detailed results of pairwise contrasts from the linear mixed-effects model analysis of SRTs (related to Figure 1F, 1G).

Analysis: scope of the comparison (Within a specific behavioral outcome, Between groups, or Forced choice). Contrast: the pair compared (e.g., S1 − S2 denotes Scene 1 vs Scene 2; AG − AB denotes Accept Good vs Accept Bad).

Estimate (ms): difference in mean SRT, in milliseconds. SE (ms): standard error of the difference. df: Satterthwaite degrees of freedom. t: t-statistic for the contrast. P: p-value adjusted for multiple comparisons (Bonferroni; values below 0.001 shown as “< 0.001”). Cohen’s d: effect-size for the pairwise difference. Effect size: qualitative magnitude of Cohen’s d.

Abbreviations: S, Scene; AG, Accept Good; AB, Accept Bad; RB, Reject Bad.

Note: The Between-groups contrasts show highly significant and large differences between behavioral outcomes (AG vs RB, AB vs RB) across all monkeys, confirming that rejection (return saccades) involves significantly longer latencies than acceptance.

| **Monkey** | **Analysis** | **Contrast** | **Estimate (ms)** | **SE (ms)** | **df** | **t** | **P (Bonferroni)** | **Cohen’s d** | **Effect size** |
| --- | --- | --- | --- | --- | --- | --- | --- | --- | --- |
| Monkey Ch | Within_AG | S1 − S2 | 2.30 | 1.03 | 4596.8 | 2.22 | 0.393 | 0.10 | Negligible |
| Monkey Ch | Within_AG | S1 − S3 | 3.05 | 1.17 | 4902.2 | 2.60 | 0.141 | 0.13 | Negligible |
| Monkey Ch | Within_AG | S1 − S4 | −0.80 | 1.03 | 4597.2 | −0.78 | 1.000 | −0.03 | Negligible |
| Monkey Ch | Within_AG | S1 − S5 | 1.84 | 1.15 | 4847.2 | 1.60 | 1.000 | 0.08 | Negligible |
| Monkey Ch | Within_AG | S1 − S6 | 5.11 | 1.04 | 4639.7 | 4.89 | 1.5 × 10⁻⁵ | 0.22 | Small |
| Monkey Ch | Within_AG | S2 − S3 | 0.75 | 1.17 | 4880.8 | 0.64 | 1.000 | 0.03 | Negligible |
| Monkey Ch | Within_AG | S2 − S4 | −3.10 | 1.03 | 4563.3 | −3.02 | 0.039 | −0.13 | Negligible |
| Monkey Ch | Within_AG | S2 − S5 | −0.46 | 1.14 | 4828.4 | −0.40 | 1.000 | −0.02 | Negligible |
| Monkey Ch | Within_AG | S2 − S6 | 2.81 | 1.04 | 4606.5 | 2.71 | 0.102 | 0.12 | Negligible |
| Monkey Ch | Within_AG | S3 − S4 | −3.85 | 1.17 | 4896.1 | −3.30 | 0.015 | −0.16 | Negligible |
| Monkey Ch | Within_AG | S3 − S5 | −1.21 | 1.27 | 4642.9 | −0.95 | 1.000 | −0.05 | Negligible |
| Monkey Ch | Within_AG | S3 − S6 | 2.07 | 1.18 | 4881.0 | 1.75 | 1.000 | 0.09 | Negligible |
| Monkey Ch | Within_AG | S4 − S5 | 2.64 | 1.14 | 4847.3 | 2.31 | 0.310 | 0.11 | Negligible |
| Monkey Ch | Within_AG | S4 − S6 | 5.91 | 1.04 | 4606.9 | 5.70 | 2.0 × 10⁻⁷ | 0.25 | Small |
| Monkey Ch | Within_AG | S5 − S6 | 3.27 | 1.15 | 4827.2 | 2.84 | 0.068 | 0.14 | Negligible |
| Monkey Ch | Within_RB | S1 − S2 | 5.65 | 1.88 | 3130.2 | 3.01 | 0.016 | 0.13 | Negligible |
| Monkey Ch | Within_RB | S1 − S4 | −3.41 | 1.93 | 3137.5 | −1.76 | 0.470 | −0.08 | Negligible |
| Monkey Ch | Within_RB | S1 − S6 | −0.67 | 1.94 | 3153.6 | −0.34 | 1.000 | −0.02 | Negligible |
| Monkey Ch | Within_RB | S2 − S4 | −9.06 | 1.91 | 3182.2 | −4.74 | 1.3 × 10⁻⁵ | −0.21 | Small |
| Monkey Ch | Within_RB | S2 − S6 | −6.32 | 1.92 | 3198.1 | −3.29 | 0.006 | −0.14 | Negligible |
| Monkey Ch | Within_RB | S4 − S6 | 2.74 | 1.98 | 3096.1 | 1.39 | 0.996 | 0.06 | Negligible |
| Monkey Ch | Between_Groups | AG − AB | −8.39 | 1.01 | 10536.8 | −8.30 | 3.6 × 10⁻¹⁶ | −0.25 | Small |
| Monkey Ch | Between_Groups | AG − RB | −44.80 | 0.68 | 10186.6 | −65.71 | < 0.001 | −1.35 | Large |
| Monkey Ch | Between_Groups | AB − RB | −36.41 | 1.04 | 10671.5 | −34.86 | 8.2 × 10⁻²⁵² | −1.10 | Large |
| Monkey Cr | Within_AG | S1 − S2 | 3.32 | 0.88 | 7118.2 | 3.78 | 0.002 | 0.13 | Negligible |
| Monkey Cr | Within_AG | S1 − S3 | 6.15 | 0.98 | 7477.5 | 6.30 | 4.8 × 10⁻⁹ | 0.24 | Small |
| Monkey Cr | Within_AG | S1 − S4 | 2.90 | 0.88 | 7140.3 | 3.29 | 0.015 | 0.11 | Negligible |
| Monkey Cr | Within_AG | S1 − S5 | 8.02 | 1.02 | 7584.6 | 7.84 | 7.8 × 10⁻¹⁴ | 0.32 | Small |
| Monkey Cr | Within_AG | S1 − S6 | 5.19 | 0.89 | 7164.5 | 5.83 | 8.7 × 10⁻⁸ | 0.20 | Small |
| Monkey Cr | Within_AG | S2 − S3 | 2.83 | 0.98 | 7439.6 | 2.88 | 0.059 | 0.11 | Negligible |
| Monkey Cr | Within_AG | S2 − S4 | −0.42 | 0.89 | 7099.6 | −0.48 | 1.000 | −0.02 | Negligible |
| Monkey Cr | Within_AG | S2 − S5 | 4.70 | 1.03 | 7540.8 | 4.58 | 7.2 × 10⁻⁵ | 0.19 | Negligible |
| Monkey Cr | Within_AG | S2 − S6 | 1.87 | 0.89 | 7128.2 | 2.09 | 0.550 | 0.07 | Negligible |
| Monkey Cr | Within_AG | S3 − S4 | −3.25 | 0.98 | 7412.1 | −3.31 | 0.014 | −0.13 | Negligible |
| Monkey Cr | Within_AG | S3 − S5 | 1.87 | 1.10 | 7201.1 | 1.70 | 1.000 | 0.07 | Negligible |
| Monkey Cr | Within_AG | S3 − S6 | −0.96 | 0.99 | 7481.1 | −0.97 | 1.000 | −0.04 | Negligible |
| Monkey Cr | Within_AG | S4 − S5 | 5.12 | 1.03 | 7521.8 | 4.97 | 1.0 × 10⁻⁵ | 0.20 | Small |
| Monkey Cr | Within_AG | S4 − S6 | 2.29 | 0.90 | 7133.7 | 2.55 | 0.161 | 0.09 | Negligible |
| Monkey Cr | Within_AG | S5 − S6 | −2.83 | 1.04 | 7589.7 | −2.72 | 0.097 | −0.11 | Negligible |
| Monkey Cr | Within_AB | S3 − S5 | 3.52 | 1.34 | 908.3 | 2.62 | 0.009 | 0.13 | Negligible |
| Monkey Cr | Within_RB | S1 − S2 | −0.41 | 1.93 | 4548.7 | −0.21 | 1.000 | −0.01 | Negligible |
| Monkey Cr | Within_RB | S1 − S4 | −2.96 | 1.93 | 4557.3 | −1.53 | 0.757 | −0.05 | Negligible |
| Monkey Cr | Within_RB | S1 − S6 | −29.13 | 2.13 | 4935.8 | −13.67 | 5.3 × 10⁻⁴¹ | −0.52 | Medium |
| Monkey Cr | Within_RB | S2 − S4 | −2.55 | 1.95 | 4535.3 | −1.31 | 1.000 | −0.05 | Negligible |
| Monkey Cr | Within_RB | S2 − S6 | −28.71 | 2.14 | 4925.5 | −13.39 | 2.1 × 10⁻³⁹ | −0.52 | Medium |
| Monkey Cr | Within_RB | S4 − S6 | −26.17 | 2.15 | 4904.2 | −12.19 | 6.4 × 10⁻³³ | −0.47 | Small |
| Monkey Cr | Between_Groups | AG − AB | −11.36 | 1.08 | 15918.7 | −10.47 | 4.3 × 10⁻²⁵ | −0.28 | Small |
| Monkey Cr | Between_Groups | AG − RB | −47.45 | 0.67 | 14998.4 | −70.61 | < 0.001 | −1.18 | Large |
| Monkey Cr | Between_Groups | AB − RB | −36.09 | 1.12 | 16015.7 | −32.17 | 1.2 × 10⁻²¹⁹ | −0.90 | Large |
| Monkey Cr | Forced choice | S7 − S8 | 2.16 | 0.96 | 4899.2 | 2.24 | 0.150 | 0.08 | Negligible |
| Monkey Cr | Forced choice | S7 − S9 | −1.88 | 0.97 | 4906.5 | −1.95 | 0.311 | −0.07 | Negligible |
| Monkey Cr | Forced choice | S7 − S10 | −5.74 | 0.99 | 4959.8 | −5.82 | 3.8 × 10⁻⁸ | −0.21 | Small |
| Monkey Cr | Forced choice | S8 − S9 | −4.04 | 0.97 | 4904.2 | −4.17 | 1.8 × 10⁻⁴ | −0.14 | Negligible |
| Monkey Cr | Forced choice | S8 − S10 | −7.90 | 0.99 | 4952.8 | −7.99 | 1.0 × 10⁻¹⁴ | −0.28 | Small |
| Monkey Cr | Forced choice | S9 − S10 | −3.86 | 0.99 | 4965.1 | −3.89 | 6.0 × 10⁻⁴ | −0.14 | Negligible |
| Monkey Sp | Within_AG | S1 − S2 | −4.40 | 1.60 | 3046.0 | −2.75 | 0.090 | −0.18 | Negligible |
| Monkey Sp | Within_AG | S1 − S3 | −0.79 | 1.60 | 3047.1 | −0.49 | 1.000 | −0.03 | Negligible |
| Monkey Sp | Within_AG | S1 − S4 | 2.58 | 1.60 | 3050.5 | 1.61 | 1.000 | 0.11 | Negligible |
| Monkey Sp | Within_AG | S1 − S5 | 8.38 | 1.60 | 3046.3 | 5.24 | 2.6 × 10⁻⁶ | 0.35 | Small |
| Monkey Sp | Within_AG | S1 − S6 | 0.20 | 1.62 | 3027.5 | 0.12 | 1.000 | 0.01 | Negligible |
| Monkey Sp | Within_AG | S2 − S3 | 3.61 | 1.32 | 2905.3 | 2.72 | 0.097 | 0.15 | Negligible |
| Monkey Sp | Within_AG | S2 − S4 | 6.97 | 1.32 | 2910.2 | 5.27 | 2.2 × 10⁻⁶ | 0.29 | Small |
| Monkey Sp | Within_AG | S2 − S5 | 12.78 | 1.33 | 2903.5 | 9.62 | 2.0 × 10⁻²⁰ | 0.53 | Medium |
| Monkey Sp | Within_AG | S2 − S6 | 4.59 | 1.36 | 2930.8 | 3.37 | 0.011 | 0.19 | Negligible |
| Monkey Sp | Within_AG | S3 − S4 | 3.37 | 1.32 | 2906.0 | 2.55 | 0.161 | 0.14 | Negligible |
| Monkey Sp | Within_AG | S3 − S5 | 9.17 | 1.33 | 2906.2 | 6.92 | 8.2 × 10⁻¹¹ | 0.38 | Small |
| Monkey Sp | Within_AG | S3 − S6 | 0.99 | 1.36 | 2933.2 | 0.73 | 1.000 | 0.04 | Negligible |
| Monkey Sp | Within_AG | S4 − S5 | 5.81 | 1.32 | 2911.1 | 4.39 | 1.8 × 10⁻⁴ | 0.24 | Small |
| Monkey Sp | Within_AG | S4 − S6 | −2.38 | 1.36 | 2937.8 | −1.75 | 1.000 | −0.10 | Negligible |
| Monkey Sp | Within_AG | S5 − S6 | −8.19 | 1.36 | 2931.8 | −6.01 | 3.2 × 10⁻⁸ | −0.34 | Small |
| Monkey Sp | Within_RB | S2 − S3 | 18.27 | 3.81 | 1842.5 | 4.80 | 1.7 × 10⁻⁵ | 0.32 | Small |
| Monkey Sp | Within_RB | S2 − S4 | 13.41 | 3.82 | 1835.3 | 3.51 | 0.005 | 0.23 | Small |
| Monkey Sp | Within_RB | S2 − S5 | 42.64 | 3.86 | 1824.6 | 11.05 | 1.5 × 10⁻²⁶ | 0.74 | Medium |
| Monkey Sp | Within_RB | S2 − S6 | 3.08 | 4.21 | 1861.8 | 0.73 | 1.000 | 0.05 | Negligible |
| Monkey Sp | Within_RB | S3 − S4 | −4.86 | 3.62 | 1777.8 | −1.34 | 1.000 | −0.08 | Negligible |
| Monkey Sp | Within_RB | S3 − S5 | 24.37 | 3.67 | 1797.5 | 6.65 | 3.9 × 10⁻¹⁰ | 0.42 | Small |
| Monkey Sp | Within_RB | S3 − S6 | −15.19 | 4.05 | 1897.9 | −3.75 | 0.002 | −0.26 | Small |
| Monkey Sp | Within_RB | S4 − S5 | 29.23 | 3.67 | 1791.5 | 7.96 | 3.1 × 10⁻¹⁴ | 0.51 | Medium |
| Monkey Sp | Within_RB | S4 − S6 | −10.33 | 4.05 | 1894.0 | −2.55 | 0.109 | −0.18 | Negligible |
| Monkey Sp | Within_RB | S5 − S6 | −39.56 | 4.09 | 1882.0 | −9.67 | 1.3 × 10⁻²⁰ | −0.68 | Medium |
| Monkey Sp | Between_Groups | AG − AB | −13.92 | 2.41 | 5783.4 | −5.78 | 2.4 × 10⁻⁸ | −0.33 | Small |
| Monkey Sp | Between_Groups | AG − RB | −108.37 | 1.14 | 5935.6 | −95.44 | < 0.001 | −2.60 | Large |
| Monkey Sp | Between_Groups | AB − RB | −94.45 | 2.46 | 5676.3 | −38.42 | 8.3 × 10⁻²⁸⁷ | −2.26 | Large |
| Monkey Sp | Forced choice | S7 − S8 | 1.99 | 2.25 | 1897.0 | 0.88 | 1.000 | 0.05 | Negligible |
| Monkey Sp | Forced choice | S7 − S9 | 3.70 | 2.27 | 1907.8 | 1.63 | 0.618 | 0.09 | Negligible |
| Monkey Sp | Forced choice | S7 − S10 | −13.84 | 2.29 | 1927.5 | −6.03 | 1.2 × 10⁻⁸ | −0.34 | Small |
| Monkey Sp | Forced choice | S8 − S9 | 1.71 | 2.27 | 1907.8 | 0.76 | 1.000 | 0.04 | Negligible |
| Monkey Sp | Forced choice | S8 − S10 | −15.83 | 2.29 | 1925.9 | −6.91 | 4.1 × 10⁻¹¹ | −0.39 | Small |
| Monkey Sp | Forced choice | S9 − S10 | −17.55 | 2.31 | 1939.1 | −7.60 | 2.8 × 10⁻¹³ | −0.43 | Small |
