## Supplemental Table 4 for "Parallel basal ganglia and frontal cortical outputs differentially encode context-dependent evaluation and categorical commitment during choice"

**Table S4. Scene-period LMM and post hoc statistics for Fig. 2C,D.**

Statistics summarize neuron-level mean z-scored activity during the scene-period analysis window, 100–300 ms after scene onset, for pooled three-monkey SNr and FEF data in Scenes 1–6.

Scene effects were tested separately for SNr and FEF using a linear mixed-effects model, z-scored activity ~ Scene + (1 | Monkey) + (1 | Neuron), followed by Bonferroni-corrected emmeans pairwise contrasts.

**A. Omnibus scene effects**

| **Brain area** | **Scene set** | **n neurons** | **F** | **df (num, den)** | **P** | **Partial η²** | **Effect size** |
| --- | --- | --- | --- | --- | --- | --- | --- |
| SNr | S1–S6 | 91 | 176.122 | 5, 450 | 1.6 × 10⁻¹⁰³ | 0.662 | Large |
| FEF | S1–S6 | 99 | 15.022 | 5, 490 | 9.8 × 10⁻¹⁴ | 0.133 | Medium |

**B. Bonferroni-corrected pairwise comparisons**

| **Brain area** | **Scene set** | **Contrast** | **Estimate** | **SE** | **df** | **t** | **Adjusted P** | **Significance** |
| --- | --- | --- | --- | --- | --- | --- | --- | --- |
| SNr | S1–S6 | S1 vs S2 | 0.113 | 0.136 | 450 | 0.831 | 1.000 | n.s. |
| SNr | S1–S6 | S1 vs S3 | −1.984 | 0.136 | 450 | −14.560 | 1.9 × 10⁻³⁸ | *** |
| SNr | S1–S6 | S1 vs S4 | −2.319 | 0.136 | 450 | −17.015 | 2.6 × 10⁻⁴⁹ | *** |
| SNr | S1–S6 | S1 vs S5 | −2.714 | 0.136 | 450 | −19.918 | 1.4 × 10⁻⁶² | *** |
| SNr | S1–S6 | S1 vs S6 | −2.485 | 0.136 | 450 | −18.235 | 7.4 × 10⁻⁵⁵ | *** |
| SNr | S1–S6 | S2 vs S3 | −2.098 | 0.136 | 450 | −15.391 | 4.5 × 10⁻⁴² | *** |
| SNr | S1–S6 | S2 vs S4 | −2.432 | 0.136 | 450 | −17.846 | 4.4 × 10⁻⁵³ | *** |
| SNr | S1–S6 | S2 vs S5 | −2.828 | 0.136 | 450 | −20.749 | 2.1 × 10⁻⁶⁶ | *** |
| SNr | S1–S6 | S2 vs S6 | −2.598 | 0.136 | 450 | −19.067 | 1.2 × 10⁻⁵⁸ | *** |
| SNr | S1–S6 | S3 vs S4 | −0.335 | 0.136 | 450 | −2.455 | 0.217 | n.s. |
| SNr | S1–S6 | S3 vs S5 | −0.730 | 0.136 | 450 | −5.358 | 2.0 × 10⁻⁶ | *** |
| SNr | S1–S6 | S3 vs S6 | −0.501 | 0.136 | 450 | −3.675 | 0.004 | ** |
| SNr | S1–S6 | S4 vs S5 | −0.396 | 0.136 | 450 | −2.903 | 0.058 | n.s. |
| SNr | S1–S6 | S4 vs S6 | −0.166 | 0.136 | 450 | −1.220 | 1.000 | n.s. |
| SNr | S1–S6 | S5 vs S6 | 0.229 | 0.136 | 450 | 1.682 | 1.000 | n.s. |
| FEF | S1–S6 | S1 vs S2 | −0.204 | 0.078 | 490 | −2.614 | 0.138 | n.s. |
| FEF | S1–S6 | S1 vs S3 | 0.170 | 0.078 | 490 | 2.182 | 0.444 | n.s. |
| FEF | S1–S6 | S1 vs S4 | 0.373 | 0.078 | 490 | 4.777 | 3.5 × 10⁻⁵ | *** |
| FEF | S1–S6 | S1 vs S5 | 0.154 | 0.078 | 490 | 1.977 | 0.729 | n.s. |
| FEF | S1–S6 | S1 vs S6 | 0.328 | 0.078 | 490 | 4.208 | 4.6 × 10⁻⁴ | *** |
| FEF | S1–S6 | S2 vs S3 | 0.374 | 0.078 | 490 | 4.796 | 3.2 × 10⁻⁵ | *** |
| FEF | S1–S6 | S2 vs S4 | 0.577 | 0.078 | 490 | 7.392 | 9.5 × 10⁻¹² | *** |
| FEF | S1–S6 | S2 vs S5 | 0.358 | 0.078 | 490 | 4.591 | 8.4 × 10⁻⁵ | *** |
| FEF | S1–S6 | S2 vs S6 | 0.532 | 0.078 | 490 | 6.823 | 4.0 × 10⁻¹⁰ | *** |
| FEF | S1–S6 | S3 vs S4 | 0.203 | 0.078 | 490 | 2.596 | 0.146 | n.s. |
| FEF | S1–S6 | S3 vs S5 | −0.016 | 0.078 | 490 | −0.205 | 1.000 | n.s. |
| FEF | S1–S6 | S3 vs S6 | 0.158 | 0.078 | 490 | 2.027 | 0.648 | n.s. |
| FEF | S1–S6 | S4 vs S5 | −0.219 | 0.078 | 490 | −2.801 | 0.080 | n.s. |
| FEF | S1–S6 | S4 vs S6 | −0.044 | 0.078 | 490 | −0.569 | 1.000 | n.s. |
| FEF | S1–S6 | S5 vs S6 | 0.174 | 0.078 | 490 | 2.232 | 0.391 | n.s. |

Note: Estimate indicates the first minus the second scene in each contrast. P values in panel B are Bonferroni-adjusted. Significance labels are n.s., not significant; *P < 0.05; **P < 0.01; ***P < 0.001. Values are rounded for display.
