## Supplemental Table 5 for "Parallel basal ganglia and frontal cortical outputs differentially encode context-dependent evaluation and categorical commitment during choice"

**Table S5. Primary paired-difference statistics for the Fig. 3 per-neuron target-period scatter plots.**

Statistics summarize paired comparisons of normalized target-period firing rates (100–300 ms after target onset) for the pooled scene 1–6 conditions. For each neuron, x and y correspond to the two conditions plotted on the scatter plot, and differences are calculated as y − x.

Primary test: two-sided paired sign-flip permutation test on the mean within-neuron difference (100,000 random sign flips; seed 20260527), with Holm correction across the 12 Fig. 3 scatter comparisons.

Confidence intervals: 95% CIs are monkey-stratified bootstrap intervals that resample neurons within each monkey while preserving the original monkey sample sizes.

Abbreviations: AG, Accept Good; AB, Accept Bad; RB, Reject Bad; SNr, substantia nigra pars reticulata; FEF, frontal eye field. Contra and Ipsi indicate target direction relative to the recording site. All P values are two-sided. Values are rounded for display.

| **Region** | **Direction** | **Comparison (y vs x)** | **n** | **n by monkey** | **Mean diff [95% CI]** | **Median diff [95% CI]** | **Sign-flip P (Holm)** | **% y > x** |
| --- | --- | --- | --- | --- | --- | --- | --- | --- |
| SNr | Contra | Accept Bad vs Accept Good | 91 | Ch 32; Cr 37; Sp 22 | 1.85 [1.63, 2.06] | 1.99 [1.71, 2.29] | 1.2 × 10⁻⁴ | 93.4% |
| SNr | Contra | Reject Bad vs Accept Good | 91 | Ch 32; Cr 37; Sp 22 | 2.50 [2.34, 2.66] | 2.60 [2.35, 2.78] | 1.2 × 10⁻⁴ | 100.0% |
| SNr | Contra | Reject Bad vs Accept Bad | 91 | Ch 32; Cr 37; Sp 22 | 0.65 [0.48, 0.84] | 0.49 [0.30, 0.73] | 1.2 × 10⁻⁴ | 80.2% |
| SNr | Ipsi | Accept Bad vs Accept Good | 91 | Ch 32; Cr 37; Sp 22 | 1.66 [1.47, 1.85] | 1.72 [1.59, 1.86] | 1.2 × 10⁻⁴ | 95.6% |
| SNr | Ipsi | Reject Bad vs Accept Good | 91 | Ch 32; Cr 37; Sp 22 | 1.99 [1.84, 2.13] | 2.04 [1.92, 2.12] | 1.2 × 10⁻⁴ | 98.9% |
| SNr | Ipsi | Reject Bad vs Accept Bad | 91 | Ch 32; Cr 37; Sp 22 | 0.32 [0.18, 0.48] | 0.16 [0.04, 0.35] | 7.5 × 10⁻⁴ | 61.5% |
| FEF | Contra | Accept Bad vs Accept Good | 98 | Ch 33; Cr 49; Sp 16 | −0.18 [−0.27, −0.08] | −0.16 [−0.31, 0.02] | 0.002 | 41.8% |
| FEF | Contra | Reject Bad vs Accept Good | 99 | Ch 33; Cr 50; Sp 16 | −0.80 [−0.97, −0.63] | −0.93 [−1.09, −0.72] | 1.2 × 10⁻⁴ | 16.2% |
| FEF | Contra | Reject Bad vs Accept Bad | 98 | Ch 33; Cr 49; Sp 16 | −0.66 [−0.83, −0.49] | −0.61 [−0.85, −0.36] | 1.2 × 10⁻⁴ | 25.5% |
| FEF | Ipsi | Accept Bad vs Accept Good | 98 | Ch 33; Cr 49; Sp 16 | −0.09 [−0.22, 0.03] | 0.00 [−0.20, 0.08] | 0.163 | 49.0% |
| FEF | Ipsi | Reject Bad vs Accept Good | 99 | Ch 33; Cr 50; Sp 16 | −0.55 [−0.75, −0.36] | −0.38 [−0.77, −0.21] | 1.2 × 10⁻⁴ | 32.3% |
| FEF | Ipsi | Reject Bad vs Accept Bad | 98 | Ch 33; Cr 49; Sp 16 | −0.47 [−0.65, −0.29] | −0.39 [−0.65, −0.18] | 1.2 × 10⁻⁴ | 32.7% |
