## Supplemental Table 6 for "Parallel basal ganglia and frontal cortical outputs differentially encode context-dependent evaluation and categorical commitment during choice"

**Table S6. Sensitivity analyses and scatter-relationship statistics for the Fig. 3 per-neuron target-period scatter plots.**

Sensitivity analyses summarize paired comparisons and scatter-relationship statistics for the same per-neuron target-period values shown in the Fig. 3 scatter plots. Deming slope confidence intervals are monkey-stratified bootstrap intervals.

Sensitivity tests: Wilcoxon signed-rank tests and paired t-tests were applied to within-neuron differences, with Holm correction across the 12 Fig. 3 scatter comparisons.

Scatter relationship: Pearson and Spearman correlations quantify per-neuron covariation; Deming regression slopes are shown with 95% monkey-stratified bootstrap confidence intervals.

Abbreviations: AG, Accept Good; AB, Accept Bad; RB, Reject Bad; SNr, substantia nigra pars reticulata; FEF, frontal eye field. Contra and Ipsi indicate target direction relative to the recording site. All P values are two-sided. Values are rounded for display.

| **Region** | **Direction** | **Comparison (y vs x)** | **Wilcoxon P (Holm)** | **Paired t P (Holm)** | **Pearson r** | **Pearson P (Holm)** | **Spearman rho** | **Spearman P (Holm)** | **Deming slope [95% CI]** |
| --- | --- | --- | --- | --- | --- | --- | --- | --- | --- |
| SNr | Contra | Accept Bad vs Accept Good | 9.9 × 10⁻¹⁵ | 4.2 × 10⁻²⁸ | 0.49 | 5.0 × 10⁻⁶ | 0.48 | 5.9 × 10⁻⁶ | 2.04 [1.38, 3.22] |
| SNr | Contra | Reject Bad vs Accept Good | 1.5 × 10⁻¹⁵ | 1.2 × 10⁻⁴⁶ | 0.47 | 9.9 × 10⁻⁶ | 0.48 | 5.9 × 10⁻⁶ | 0.86 [0.48, 1.39] |
| SNr | Contra | Reject Bad vs Accept Bad | 1.5 × 10⁻⁹ | 7.8 × 10⁻⁹ | 0.63 | 1.9 × 10⁻¹⁰ | 0.68 | 8.7 × 10⁻¹³ | 0.51 [0.32, 0.71] |
| SNr | Ipsi | Accept Bad vs Accept Good | 1.3 × 10⁻¹⁴ | 1.0 × 10⁻²⁶ | 0.26 | 0.013 | 0.29 | 0.005 | 2.35 [1.09, 8.43] |
| SNr | Ipsi | Reject Bad vs Accept Good | 1.5 × 10⁻¹⁵ | 1.8 × 10⁻⁴³ | 0.44 | 2.3 × 10⁻⁵ | 0.40 | 1.4 × 10⁻⁴ | 0.73 [0.47, 1.10] |
| SNr | Ipsi | Reject Bad vs Accept Bad | 0.003 | 0.001 | 0.47 | 9.9 × 10⁻⁶ | 0.55 | 1.4 × 10⁻⁷ | 0.46 [0.28, 0.70] |
| FEF | Contra | Accept Bad vs Accept Good | 0.003 | 0.002 | 0.78 | 4.5 × 10⁻²⁰ | 0.74 | 3.2 × 10⁻¹⁷ | 1.10 [0.92, 1.30] |
| FEF | Contra | Reject Bad vs Accept Good | 6.5 × 10⁻¹¹ | 5.0 × 10⁻¹⁴ | 0.60 | 3.1 × 10⁻¹⁰ | 0.55 | 3.3 × 10⁻⁸ | 1.81 [1.51, 2.48] |
| FEF | Contra | Reject Bad vs Accept Bad | 1.2 × 10⁻⁸ | 3.9 × 10⁻¹⁰ | 0.59 | 1.2 × 10⁻⁹ | 0.49 | 1.5 × 10⁻⁶ | 1.65 [1.35, 2.22] |
| FEF | Ipsi | Accept Bad vs Accept Good | 0.186 | 0.162 | 0.75 | 1.1 × 10⁻¹⁷ | 0.75 | 1.1 × 10⁻¹⁷ | 1.02 [0.84, 1.22] |
| FEF | Ipsi | Reject Bad vs Accept Good | 3.0 × 10⁻⁵ | 3.7 × 10⁻⁶ | 0.55 | 2.4 × 10⁻⁸ | 0.51 | 3.5 × 10⁻⁷ | 1.70 [1.35, 2.31] |
| FEF | Ipsi | Reject Bad vs Accept Bad | 4.4 × 10⁻⁵ | 9.8 × 10⁻⁶ | 0.64 | 9.5 × 10⁻¹² | 0.59 | 2.1 × 10⁻⁹ | 1.52 [1.24, 2.06] |
