## Supplemental Table 7 for "Parallel basal ganglia and frontal cortical outputs differentially encode context-dependent evaluation and categorical commitment during choice"

**Table S7. Fixed-effect estimates of task variables and saccade reaction time on neural activity.**

Coefficients (β) estimated from a multivariable linear mixed-effects model (LMM) fitted to normalized firing rates during the target period, corresponding to the results shown in Figure 4A. The model included Action (Accept vs. Reject), Rank (higher-ranked vs. lower-ranked within the current scene), Reward amount (reward magnitude), Direction (Contralateral vs. Ipsilateral), and SRT (log-transformed and standardized saccade reaction time) as fixed effects, with a random intercept for neurons.

Values indicate the coefficient estimate followed by the standard error in parentheses.

Coding: Action: Accept (1), Reject (−1); Rank: Higher-ranked (1), Lower-ranked (−1); Direction: Contralateral (1), Ipsilateral (−1); SRT: Z-scored log-transformed reaction time.

Significance: Asterisks indicate statistical significance derived from the lmerTest package (* P < 0.05, *** P < 0.001).

Note: In SNr, Rank is the strongest predictor (large negative coefficient), whereas in FEF, Action and Direction are the dominant predictors. SRT shows a small but significant effect in SNr, but no significant effect in FEF.

| **Region** | **Action** | **Rank** | **Reward amount** | **Direction** | **SRT (log)** |
| --- | --- | --- | --- | --- | --- |
| **SNr** | −0.19 (0.01) *** | −0.60 (0.01) *** | 0.03 (0.01) *** | 0.02 (0.01) * | 0.02 (0.01) * |
| **FEF** | 0.35 (0.03) *** | 0.03 (0.03) | 0.04 (0.02) * | 0.66 (0.01) *** | 0.00 (0.02) |
