## Supplemental Table 8 for "Parallel basal ganglia and frontal cortical outputs differentially encode context-dependent evaluation and categorical commitment during choice"

**Table S8. Descriptive statistics of regression coefficients for classified neuron types.**

Summary of fixed-effect coefficients (β) for each neuron class identified in the single-neuron classification analysis (corresponding to Figure 6C).

Neurons were classified based on the task variable showing the strongest significant modulation in time-resolved single-neuron multiple regression (Gaussian GLM) (see Methods for classification criteria).

Neuron type (n): the primary variable encoded by the neuron group, with the number of neurons (n) assigned to that class. Variable: the task variable for which the coefficient statistics are shown. Mean β: average coefficient across neurons in the class. SD: standard deviation (—, not defined for single-neuron classes).

Note: For SNr, the dominant coefficients are typically negative (e.g., Action-type neurons show a large negative β for Action), consistent with the inhibitory nature of SNr output, in which acceptance or higher-ranked offers tend to correspond to suppression. For FEF, dominant coefficients are typically positive (e.g., Action-type neurons show a large positive β for Action), consistent with stronger activity for acceptance.

| **Brain area** | **Neuron type (n)** | **Variable** | **Mean β** | **SD** |
| --- | --- | --- | --- | --- |
| SNr | Action (n = 12) | Action | −0.624 | 0.198 |
| SNr | Action (n = 12) | Rank | −0.137 | 0.186 |
| SNr | Action (n = 12) | Reward amount | 0.042 | 0.227 |
| SNr | Action (n = 12) | Direction | 0.041 | 0.093 |
| SNr | Action (n = 12) | SRT | −0.017 | 0.085 |
| SNr | Rank (n = 64) | Action | −0.091 | 0.153 |
| SNr | Rank (n = 64) | Rank | −0.584 | 0.248 |
| SNr | Rank (n = 64) | Reward amount | 0.029 | 0.166 |
| SNr | Rank (n = 64) | Direction | 0.011 | 0.116 |
| SNr | Rank (n = 64) | SRT | 0.021 | 0.068 |
| SNr | Reward amount (n = 1) | Action | −0.132 | — |
| SNr | Reward amount (n = 1) | Rank | −0.072 | — |
| SNr | Reward amount (n = 1) | Reward amount | −0.224 | — |
| SNr | Reward amount (n = 1) | Direction | 0.020 | — |
| SNr | Reward amount (n = 1) | SRT | 0.080 | — |
| SNr | SRT (n = 1) | Action | −0.127 | — |
| SNr | SRT (n = 1) | Rank | −0.074 | — |
| SNr | SRT (n = 1) | Reward amount | 0.119 | — |
| SNr | SRT (n = 1) | Direction | −0.091 | — |
| SNr | SRT (n = 1) | SRT | −0.162 | — |
| FEF | Action (n = 53) | Action | 0.542 | 0.242 |
| FEF | Action (n = 53) | Rank | −0.105 | 0.245 |
| FEF | Action (n = 53) | Reward amount | 0.005 | 0.099 |
| FEF | Action (n = 53) | Direction | 0.114 | 0.172 |
| FEF | Action (n = 53) | SRT | −0.048 | 0.083 |
| FEF | Rank (n = 4) | Action | −0.142 | 0.348 |
| FEF | Rank (n = 4) | Rank | 0.165 | 0.091 |
| FEF | Rank (n = 4) | Reward amount | −0.012 | 0.133 |
| FEF | Rank (n = 4) | Direction | −0.111 | 0.074 |
| FEF | Rank (n = 4) | SRT | −0.014 | 0.056 |
| FEF | Reward amount (n = 5) | Action | −0.122 | 0.221 |
| FEF | Reward amount (n = 5) | Rank | −0.132 | 0.228 |
| FEF | Reward amount (n = 5) | Reward amount | 0.172 | 0.082 |
| FEF | Reward amount (n = 5) | Direction | −0.048 | 0.198 |
| FEF | Reward amount (n = 5) | SRT | −0.029 | 0.029 |
| FEF | Direction (n = 23) | Action | 0.140 | 0.231 |
| FEF | Direction (n = 23) | Rank | 0.029 | 0.128 |
| FEF | Direction (n = 23) | Reward amount | 0.024 | 0.081 |
| FEF | Direction (n = 23) | Direction | 0.528 | 0.194 |
| FEF | Direction (n = 23) | SRT | −0.044 | 0.106 |
| FEF | SRT (n = 2) | Action | −0.432 | 0.084 |
| FEF | SRT (n = 2) | Rank | −0.061 | 0.010 |
| FEF | SRT (n = 2) | Reward amount | −0.019 | 0.084 |
| FEF | SRT (n = 2) | Direction | −0.026 | 0.022 |
| FEF | SRT (n = 2) | SRT | 0.150 | 0.034 |
