## Supplemental Table 9 for "Parallel basal ganglia and frontal cortical outputs differentially encode context-dependent evaluation and categorical commitment during choice"

**Table S9. LMM statistics for pooled contra target-onset conditions in Fig. S4.**

Statistics compare pooled scene 1–6 normalized target-period firing rates among Accept Good (AG), Accept Bad (AB), and Reject Bad (RB), separately for each monkey and recording area.

Model: linear mixed-effects model, z-scored activity ~ condition + (1 | Neuron), with Kenward–Roger degrees of freedom. Pairwise post-hoc comparisons use Bonferroni correction.

Abbreviations: AG, Accept Good; AB, Accept Bad; RB, Reject Bad; SNr, substantia nigra pars reticulata; FEF, frontal eye field. Values are rounded for display.

| **Region** | **Monkey** | **n** | **Set** | **F** | **df (num, den)** | **Overall P** | **Partial η²** | **Post-hoc** | **Estimate** | **SE** | **t** | **Post-hoc P** | **Sig.** |
| --- | --- | --- | --- | --- | --- | --- | --- | --- | --- | --- | --- | --- | --- |
| SNr | Ch | 32 | Pooled AG/AB/RB | 217.82 | 2, 62 | 9.1 × 10⁻²⁹ | 0.875 | AG vs AB | −2.09 | 0.13 | −16.23 | 1.6 × 10⁻²³ | *** |
| SNr | Ch | 32 | Pooled AG/AB/RB | 217.82 | 2, 62 | 9.1 × 10⁻²⁹ | 0.875 | AG vs RB | −2.51 | 0.13 | −19.48 | 1.2 × 10⁻²⁷ | *** |
| SNr | Ch | 32 | Pooled AG/AB/RB | 217.82 | 2, 62 | 9.1 × 10⁻²⁹ | 0.875 | AB vs RB | −0.42 | 0.13 | −3.25 | 0.006 | ** |
| SNr | Cr | 37 | Pooled AG/AB/RB | 112.23 | 2, 72 | 7.5 × 10⁻²³ | 0.757 | AG vs AB | −1.63 | 0.16 | −10.40 | 1.7 × 10⁻¹⁵ | *** |
| SNr | Cr | 37 | Pooled AG/AB/RB | 112.23 | 2, 72 | 7.5 × 10⁻²³ | 0.757 | AG vs RB | −2.28 | 0.16 | −14.54 | 1.2 × 10⁻²² | *** |
| SNr | Cr | 37 | Pooled AG/AB/RB | 112.23 | 2, 72 | 7.5 × 10⁻²³ | 0.757 | AB vs RB | −0.65 | 0.16 | −4.14 | 2.8 × 10⁻⁴ | *** |
| SNr | Sp | 22 | Pooled AG/AB/RB | 75.96 | 2, 42 | 1.1 × 10⁻¹⁴ | 0.783 | AG vs AB | −1.87 | 0.24 | −7.91 | 2.3 × 10⁻⁹ | *** |
| SNr | Sp | 22 | Pooled AG/AB/RB | 75.96 | 2, 42 | 1.1 × 10⁻¹⁴ | 0.783 | AG vs RB | −2.88 | 0.24 | −12.14 | 7.6 × 10⁻¹⁵ | *** |
| SNr | Sp | 22 | Pooled AG/AB/RB | 75.96 | 2, 42 | 1.1 × 10⁻¹⁴ | 0.783 | AB vs RB | −1.00 | 0.24 | −4.24 | 3.7 × 10⁻⁴ | *** |
| FEF | Ch | 33 | Pooled AG/AB/RB | 40.68 | 2, 64 | 4.0 × 10⁻¹² | 0.560 | AG vs AB | 0.16 | 0.12 | 1.38 | 0.516 | n.s. |
| FEF | Ch | 33 | Pooled AG/AB/RB | 40.68 | 2, 64 | 4.0 × 10⁻¹² | 0.560 | AG vs RB | 0.97 | 0.12 | 8.41 | 1.8 × 10⁻¹¹ | *** |
| FEF | Ch | 33 | Pooled AG/AB/RB | 40.68 | 2, 64 | 4.0 × 10⁻¹² | 0.560 | AB vs RB | 0.81 | 0.12 | 7.03 | 5.0 × 10⁻⁹ | *** |
| FEF | Cr | 49 | Pooled AG/AB/RB | 20.94 | 2, 96 | 2.8 × 10⁻⁸ | 0.304 | AG vs AB | 0.10 | 0.12 | 0.87 | 1.000 | n.s. |
| FEF | Cr | 49 | Pooled AG/AB/RB | 20.94 | 2, 96 | 2.8 × 10⁻⁸ | 0.304 | AG vs RB | 0.70 | 0.12 | 5.99 | 1.1 × 10⁻⁷ | *** |
| FEF | Cr | 49 | Pooled AG/AB/RB | 20.94 | 2, 96 | 2.8 × 10⁻⁸ | 0.304 | AB vs RB | 0.60 | 0.12 | 5.12 | 4.7 × 10⁻⁶ | *** |
| FEF | Sp | 16 | Pooled AG/AB/RB | 14.19 | 2, 30 | 4.6 × 10⁻⁵ | 0.486 | AG vs AB | 0.44 | 0.18 | 2.44 | 0.063 | n.s. |
| FEF | Sp | 16 | Pooled AG/AB/RB | 14.19 | 2, 30 | 4.6 × 10⁻⁵ | 0.486 | AG vs RB | 0.95 | 0.18 | 5.32 | 2.8 × 10⁻⁵ | *** |
| FEF | Sp | 16 | Pooled AG/AB/RB | 14.19 | 2, 30 | 4.6 × 10⁻⁵ | 0.486 | AB vs RB | 0.52 | 0.18 | 2.88 | 0.022 | * |
