## Supplemental Table 10 for "Parallel basal ganglia and frontal cortical outputs differentially encode context-dependent evaluation and categorical commitment during choice"

**Table S10. Scene-level permutation statistics for contra target-onset panels in Fig. S4.**

Statistics compare scene-specific normalized target-period firing rates within each target-condition group, separately for each monkey and recording area.

Test: repeated-measures permutation ANOVA across scenes (10,000 permutations; seed 20260528). When the overall test was significant, pairwise sign-flip permutation tests were performed with Bonferroni correction.

Mean diff is the first minus the second scene for the listed post-hoc contrast. Pairwise tests are shown only for groups with a significant overall permutation ANOVA, matching the Fig. S4 plotting logic.

| **Region** | **Monkey** | **Group** | **Scenes** | **n** | **F** | **Overall P** | **Post-hoc** | **n pairs** | **Mean diff** | **Raw P** | **Adj. P** | **Sig.** |
| --- | --- | --- | --- | --- | --- | --- | --- | --- | --- | --- | --- | --- |
| SNr | Ch | Accept Good | S1/S2/S3/S4/S5/S6 | 31 | 0.44 | 0.825 | Overall only |  |  |  |  |  |
| SNr | Ch | Accept Bad | S3/S5 | 31 | 17.17 | 4.0 × 10⁻⁴ | S3 vs S5 | 31 | −0.72 | 3.0 × 10⁻⁴ | 3.0 × 10⁻⁴ | *** |
| SNr | Ch | Reject Bad | S1/S2/S4/S6 | 32 | 5.23 | 0.002 | S1 vs S2 | 32 | 0.16 | 0.247 | 1.000 | n.s. |
| SNr | Ch | Reject Bad | S1/S2/S4/S6 | 32 | 5.23 | 0.002 | S1 vs S4 | 32 | −0.20 | 0.149 | 0.897 | n.s. |
| SNr | Ch | Reject Bad | S1/S2/S4/S6 | 32 | 5.23 | 0.002 | S1 vs S6 | 32 | 0.28 | 0.034 | 0.203 | n.s. |
| SNr | Ch | Reject Bad | S1/S2/S4/S6 | 32 | 5.23 | 0.002 | S2 vs S4 | 32 | −0.36 | 0.002 | 0.012 | * |
| SNr | Ch | Reject Bad | S1/S2/S4/S6 | 32 | 5.23 | 0.002 | S2 vs S6 | 32 | 0.12 | 0.432 | 1.000 | n.s. |
| SNr | Ch | Reject Bad | S1/S2/S4/S6 | 32 | 5.23 | 0.002 | S4 vs S6 | 32 | 0.48 | 2.0 × 10⁻⁴ | 0.001 | ** |
| SNr | Cr | Accept Good | S1/S2/S3/S4/S5/S6 | 37 | 2.76 | 0.021 | S1 vs S2 | 37 | 0.08 | 0.648 | 1.000 | n.s. |
| SNr | Cr | Accept Good | S1/S2/S3/S4/S5/S6 | 37 | 2.76 | 0.021 | S1 vs S3 | 37 | 0.39 | 0.037 | 0.562 | n.s. |
| SNr | Cr | Accept Good | S1/S2/S3/S4/S5/S6 | 37 | 2.76 | 0.021 | S1 vs S4 | 37 | 0.62 | 1.0 × 10⁻⁴ | 0.001 | ** |
| SNr | Cr | Accept Good | S1/S2/S3/S4/S5/S6 | 37 | 2.76 | 0.021 | S1 vs S5 | 37 | 0.26 | 0.146 | 1.000 | n.s. |
| SNr | Cr | Accept Good | S1/S2/S3/S4/S5/S6 | 37 | 2.76 | 0.021 | S1 vs S6 | 37 | 0.48 | 0.043 | 0.648 | n.s. |
| SNr | Cr | Accept Good | S1/S2/S3/S4/S5/S6 | 37 | 2.76 | 0.021 | S2 vs S3 | 37 | 0.31 | 0.115 | 1.000 | n.s. |
| SNr | Cr | Accept Good | S1/S2/S3/S4/S5/S6 | 37 | 2.76 | 0.021 | S2 vs S4 | 37 | 0.55 | 0.006 | 0.090 | n.s. |
| SNr | Cr | Accept Good | S1/S2/S3/S4/S5/S6 | 37 | 2.76 | 0.021 | S2 vs S5 | 37 | 0.18 | 0.451 | 1.000 | n.s. |
| SNr | Cr | Accept Good | S1/S2/S3/S4/S5/S6 | 37 | 2.76 | 0.021 | S2 vs S6 | 37 | 0.40 | 0.085 | 1.000 | n.s. |
| SNr | Cr | Accept Good | S1/S2/S3/S4/S5/S6 | 37 | 2.76 | 0.021 | S3 vs S4 | 37 | 0.23 | 0.083 | 1.000 | n.s. |
| SNr | Cr | Accept Good | S1/S2/S3/S4/S5/S6 | 37 | 2.76 | 0.021 | S3 vs S5 | 37 | −0.13 | 0.558 | 1.000 | n.s. |
| SNr | Cr | Accept Good | S1/S2/S3/S4/S5/S6 | 37 | 2.76 | 0.021 | S3 vs S6 | 37 | 0.09 | 0.734 | 1.000 | n.s. |
| SNr | Cr | Accept Good | S1/S2/S3/S4/S5/S6 | 37 | 2.76 | 0.021 | S4 vs S5 | 37 | −0.36 | 0.075 | 1.000 | n.s. |
| SNr | Cr | Accept Good | S1/S2/S3/S4/S5/S6 | 37 | 2.76 | 0.021 | S4 vs S6 | 37 | −0.14 | 0.540 | 1.000 | n.s. |
| SNr | Cr | Accept Good | S1/S2/S3/S4/S5/S6 | 37 | 2.76 | 0.021 | S5 vs S6 | 37 | 0.22 | 0.279 | 1.000 | n.s. |
| SNr | Cr | Accept Bad | S3/S5 | 22 | 0.50 | 0.488 | Overall only |  |  |  |  |  |
| SNr | Cr | Reject Bad | S1/S2/S4/S6 | 36 | 1.08 | 0.360 | Overall only |  |  |  |  |  |
| SNr | Sp | Accept Good | S1/S2/S3/S4/S5/S6 | 22 | 5.65 | 3.0 × 10⁻⁴ | S1 vs S2 | 22 | −0.01 | 0.954 | 1.000 | n.s. |
| SNr | Sp | Accept Good | S1/S2/S3/S4/S5/S6 | 22 | 5.65 | 3.0 × 10⁻⁴ | S1 vs S3 | 22 | 0.52 | 0.023 | 0.340 | n.s. |
| SNr | Sp | Accept Good | S1/S2/S3/S4/S5/S6 | 22 | 5.65 | 3.0 × 10⁻⁴ | S1 vs S4 | 22 | 0.53 | 0.016 | 0.234 | n.s. |
| SNr | Sp | Accept Good | S1/S2/S3/S4/S5/S6 | 22 | 5.65 | 3.0 × 10⁻⁴ | S1 vs S5 | 22 | 0.31 | 0.145 | 1.000 | n.s. |
| SNr | Sp | Accept Good | S1/S2/S3/S4/S5/S6 | 22 | 5.65 | 3.0 × 10⁻⁴ | S1 vs S6 | 22 | −0.46 | 0.146 | 1.000 | n.s. |
| SNr | Sp | Accept Good | S1/S2/S3/S4/S5/S6 | 22 | 5.65 | 3.0 × 10⁻⁴ | S2 vs S3 | 22 | 0.53 | 0.022 | 0.327 | n.s. |
| SNr | Sp | Accept Good | S1/S2/S3/S4/S5/S6 | 22 | 5.65 | 3.0 × 10⁻⁴ | S2 vs S4 | 22 | 0.54 | 0.005 | 0.082 | n.s. |
| SNr | Sp | Accept Good | S1/S2/S3/S4/S5/S6 | 22 | 5.65 | 3.0 × 10⁻⁴ | S2 vs S5 | 22 | 0.32 | 0.109 | 1.000 | n.s. |
| SNr | Sp | Accept Good | S1/S2/S3/S4/S5/S6 | 22 | 5.65 | 3.0 × 10⁻⁴ | S2 vs S6 | 22 | −0.45 | 0.156 | 1.000 | n.s. |
| SNr | Sp | Accept Good | S1/S2/S3/S4/S5/S6 | 22 | 5.65 | 3.0 × 10⁻⁴ | S3 vs S4 | 22 | 0.01 | 0.959 | 1.000 | n.s. |
| SNr | Sp | Accept Good | S1/S2/S3/S4/S5/S6 | 22 | 5.65 | 3.0 × 10⁻⁴ | S3 vs S5 | 22 | −0.21 | 0.343 | 1.000 | n.s. |
| SNr | Sp | Accept Good | S1/S2/S3/S4/S5/S6 | 22 | 5.65 | 3.0 × 10⁻⁴ | S3 vs S6 | 22 | −0.98 | 0.003 | 0.043 | * |
| SNr | Sp | Accept Good | S1/S2/S3/S4/S5/S6 | 22 | 5.65 | 3.0 × 10⁻⁴ | S4 vs S5 | 22 | −0.22 | 0.247 | 1.000 | n.s. |
| SNr | Sp | Accept Good | S1/S2/S3/S4/S5/S6 | 22 | 5.65 | 3.0 × 10⁻⁴ | S4 vs S6 | 22 | −0.99 | 3.0 × 10⁻⁴ | 0.004 | ** |
| SNr | Sp | Accept Good | S1/S2/S3/S4/S5/S6 | 22 | 5.65 | 3.0 × 10⁻⁴ | S5 vs S6 | 22 | −0.77 | 0.020 | 0.297 | n.s. |
| SNr | Sp | Accept Bad | S1 | 22 | — | — | Overall only |  |  |  |  |  |
| SNr | Sp | Reject Bad | S2/S3/S4/S5/S6 | 20 | 0.55 | 0.704 | Overall only |  |  |  |  |  |
| FEF | Ch | Accept Good | S1/S2/S3/S4/S5/S6 | 33 | 1.42 | 0.219 | Overall only |  |  |  |  |  |
| FEF | Ch | Accept Bad | S3/S5 | 33 | 7.45 | 0.010 | S3 vs S5 | 33 | 0.31 | 0.010 | 0.010 | ** |
| FEF | Ch | Reject Bad | S1/S2/S4/S6 | 33 | 1.31 | 0.278 | Overall only |  |  |  |  |  |
| FEF | Cr | Accept Good | S1/S2/S3/S4/S5/S6 | 50 | 0.44 | 0.814 | Overall only |  |  |  |  |  |
| FEF | Cr | Accept Bad | S3/S5 | 49 | 0.10 | 0.757 | Overall only |  |  |  |  |  |
| FEF | Cr | Reject Bad | S1/S2/S4/S6 | 49 | 3.52 | 0.017 | S1 vs S2 | 49 | −0.12 | 0.216 | 1.000 | n.s. |
| FEF | Cr | Reject Bad | S1/S2/S4/S6 | 49 | 3.52 | 0.017 | S1 vs S4 | 49 | −0.21 | 0.056 | 0.337 | n.s. |
| FEF | Cr | Reject Bad | S1/S2/S4/S6 | 49 | 3.52 | 0.017 | S1 vs S6 | 49 | 0.12 | 0.315 | 1.000 | n.s. |
| FEF | Cr | Reject Bad | S1/S2/S4/S6 | 49 | 3.52 | 0.017 | S2 vs S4 | 49 | −0.09 | 0.393 | 1.000 | n.s. |
| FEF | Cr | Reject Bad | S1/S2/S4/S6 | 49 | 3.52 | 0.017 | S2 vs S6 | 49 | 0.25 | 0.062 | 0.373 | n.s. |
| FEF | Cr | Reject Bad | S1/S2/S4/S6 | 49 | 3.52 | 0.017 | S4 vs S6 | 49 | 0.34 | 0.002 | 0.011 | * |
| FEF | Sp | Accept Good | S1/S2/S3/S4/S5/S6 | 16 | 0.45 | 0.819 | Overall only |  |  |  |  |  |
| FEF | Sp | Accept Bad | S1 | 16 | — | — | Overall only |  |  |  |  |  |
| FEF | Sp | Reject Bad | S2/S3/S4/S5/S6 | 16 | 0.90 | 0.475 | Overall only |  |  |  |  |  |
