## Supplemental Table 11 for "Parallel basal ganglia and frontal cortical outputs differentially encode context-dependent evaluation and categorical commitment during choice"

**Table S11. LMM statistics for pooled ipsi target-onset conditions in Fig. S5.**

Statistics compare pooled scene 1–6 normalized target-period firing rates among Accept Good (AG), Accept Bad (AB), and Reject Bad (RB), separately for each monkey and recording area.

Model: linear mixed-effects model, z-scored activity ~ condition + (1 | Neuron), with Kenward–Roger degrees of freedom. Pairwise post-hoc comparisons use Bonferroni correction.

Abbreviations: AG, Accept Good; AB, Accept Bad; RB, Reject Bad; SNr, substantia nigra pars reticulata; FEF, frontal eye field. Values are rounded for display.

| **Region** | **Monkey** | **n** | **Set** | **F** | **df (num, den)** | **Overall P** | **Partial η²** | **Post-hoc** | **Estimate** | **SE** | **t** | **Post-hoc P** | **Sig.** |
| --- | --- | --- | --- | --- | --- | --- | --- | --- | --- | --- | --- | --- | --- |
| SNr | Ch | 32 | Pooled AG/AB/RB | 255.47 | 2, 62 | 1.2 × 10⁻³⁰ | 0.892 | AG vs AB | −2.12 | 0.11 | −19.27 | 2.2 × 10⁻²⁷ | *** |
| SNr | Ch | 32 | Pooled AG/AB/RB | 255.47 | 2, 62 | 1.2 × 10⁻³⁰ | 0.892 | AG vs RB | −2.18 | 0.11 | −19.87 | 4.2 × 10⁻²⁸ | *** |
| SNr | Ch | 32 | Pooled AG/AB/RB | 255.47 | 2, 62 | 1.2 × 10⁻³⁰ | 0.892 | AB vs RB | −0.07 | 0.11 | −0.60 | 1.000 | n.s. |
| SNr | Cr | 37 | Pooled AG/AB/RB | 76.86 | 2, 72 | 1.4 × 10⁻¹⁸ | 0.681 | AG vs AB | −1.64 | 0.16 | −10.21 | 3.7 × 10⁻¹⁵ | *** |
| SNr | Cr | 37 | Pooled AG/AB/RB | 76.86 | 2, 72 | 1.4 × 10⁻¹⁸ | 0.681 | AG vs RB | −1.80 | 0.16 | −11.20 | 6.0 × 10⁻¹⁷ | *** |
| SNr | Cr | 37 | Pooled AG/AB/RB | 76.86 | 2, 72 | 1.4 × 10⁻¹⁸ | 0.681 | AB vs RB | −0.16 | 0.16 | −0.99 | 0.977 | n.s. |
| SNr | Sp | 22 | Pooled AG/AB/RB | 94.86 | 2, 42 | 2.7 × 10⁻¹⁶ | 0.819 | AG vs AB | −1.04 | 0.15 | −7.14 | 2.8 × 10⁻⁸ | *** |
| SNr | Sp | 22 | Pooled AG/AB/RB | 94.86 | 2, 42 | 2.7 × 10⁻¹⁶ | 0.819 | AG vs RB | −2.01 | 0.15 | −13.77 | 1.1 × 10⁻¹⁶ | *** |
| SNr | Sp | 22 | Pooled AG/AB/RB | 94.86 | 2, 42 | 2.7 × 10⁻¹⁶ | 0.819 | AB vs RB | −0.97 | 0.15 | −6.64 | 1.5 × 10⁻⁷ | *** |
| FEF | Ch | 33 | Pooled AG/AB/RB | 28.19 | 2, 64 | 1.7 × 10⁻⁹ | 0.468 | AG vs AB | 0.26 | 0.14 | 1.89 | 0.191 | n.s. |
| FEF | Ch | 33 | Pooled AG/AB/RB | 28.19 | 2, 64 | 1.7 × 10⁻⁹ | 0.468 | AG vs RB | 0.98 | 0.14 | 7.24 | 2.1 × 10⁻⁹ | *** |
| FEF | Ch | 33 | Pooled AG/AB/RB | 28.19 | 2, 64 | 1.7 × 10⁻⁹ | 0.468 | AB vs RB | 0.72 | 0.14 | 5.35 | 3.8 × 10⁻⁶ | *** |
| FEF | Cr | 49 | Pooled AG/AB/RB | 5.07 | 2, 96 | 0.008 | 0.096 | AG vs AB | 0.04 | 0.14 | 0.30 | 1.000 | n.s. |
| FEF | Cr | 49 | Pooled AG/AB/RB | 5.07 | 2, 96 | 0.008 | 0.096 | AG vs RB | 0.40 | 0.14 | 2.90 | 0.014 | * |
| FEF | Cr | 49 | Pooled AG/AB/RB | 5.07 | 2, 96 | 0.008 | 0.096 | AB vs RB | 0.36 | 0.14 | 2.60 | 0.033 | * |
| FEF | Sp | 16 | Pooled AG/AB/RB | 1.43 | 2, 30 | 0.255 | 0.087 | AG vs AB | −0.09 | 0.18 | −0.50 | 1.000 | n.s. |
| FEF | Sp | 16 | Pooled AG/AB/RB | 1.43 | 2, 30 | 0.255 | 0.087 | AG vs RB | 0.21 | 0.18 | 1.15 | 0.780 | n.s. |
| FEF | Sp | 16 | Pooled AG/AB/RB | 1.43 | 2, 30 | 0.255 | 0.087 | AB vs RB | 0.30 | 0.18 | 1.65 | 0.329 | n.s. |
