## Supplemental Table 12 for "Parallel basal ganglia and frontal cortical outputs differentially encode context-dependent evaluation and categorical commitment during choice"

**Table S12. Scene-level permutation statistics for ipsi target-onset panels in Fig. S5.**

Statistics compare scene-specific normalized target-period firing rates within each target-condition group, separately for each monkey and recording area.

Test: repeated-measures permutation ANOVA across scenes (10,000 permutations; seed 20260528). When the overall test was significant, pairwise sign-flip permutation tests were performed with Bonferroni correction.

Mean diff is the first minus the second scene for the listed post-hoc contrast. Pairwise tests are shown only for groups with a significant overall permutation ANOVA, matching the Fig. S5 plotting logic.

| **Region** | **Monkey** | **Group** | **Scenes** | **n** | **F** | **Overall P** | **Post-hoc** | **n pairs** | **Mean diff** | **Raw P** | **Adj. P** | **Sig.** |
| --- | --- | --- | --- | --- | --- | --- | --- | --- | --- | --- | --- | --- |
| SNr | Ch | Accept Good | S1/S2/S3/S4/S5/S6 | 32 | 5.39 | 2.0 × 10⁻⁴ | S1 vs S2 | 32 | 0.57 | 0.012 | 0.187 | n.s. |
| SNr | Ch | Accept Good | S1/S2/S3/S4/S5/S6 | 32 | 5.39 | 2.0 × 10⁻⁴ | S1 vs S3 | 32 | 0.18 | 0.345 | 1.000 | n.s. |
| SNr | Ch | Accept Good | S1/S2/S3/S4/S5/S6 | 32 | 5.39 | 2.0 × 10⁻⁴ | S1 vs S4 | 32 | 0.47 | 0.037 | 0.558 | n.s. |
| SNr | Ch | Accept Good | S1/S2/S3/S4/S5/S6 | 32 | 5.39 | 2.0 × 10⁻⁴ | S1 vs S5 | 32 | 0.59 | 0.017 | 0.250 | n.s. |
| SNr | Ch | Accept Good | S1/S2/S3/S4/S5/S6 | 32 | 5.39 | 2.0 × 10⁻⁴ | S1 vs S6 | 32 | 1.02 | 1.0 × 10⁻³ | 0.015 | * |
| SNr | Ch | Accept Good | S1/S2/S3/S4/S5/S6 | 32 | 5.39 | 2.0 × 10⁻⁴ | S2 vs S3 | 32 | −0.39 | 0.123 | 1.000 | n.s. |
| SNr | Ch | Accept Good | S1/S2/S3/S4/S5/S6 | 32 | 5.39 | 2.0 × 10⁻⁴ | S2 vs S4 | 32 | −0.10 | 0.570 | 1.000 | n.s. |
| SNr | Ch | Accept Good | S1/S2/S3/S4/S5/S6 | 32 | 5.39 | 2.0 × 10⁻⁴ | S2 vs S5 | 32 | 0.01 | 0.948 | 1.000 | n.s. |
| SNr | Ch | Accept Good | S1/S2/S3/S4/S5/S6 | 32 | 5.39 | 2.0 × 10⁻⁴ | S2 vs S6 | 32 | 0.45 | 0.136 | 1.000 | n.s. |
| SNr | Ch | Accept Good | S1/S2/S3/S4/S5/S6 | 32 | 5.39 | 2.0 × 10⁻⁴ | S3 vs S4 | 32 | 0.29 | 0.082 | 1.000 | n.s. |
| SNr | Ch | Accept Good | S1/S2/S3/S4/S5/S6 | 32 | 5.39 | 2.0 × 10⁻⁴ | S3 vs S5 | 32 | 0.40 | 0.038 | 0.571 | n.s. |
| SNr | Ch | Accept Good | S1/S2/S3/S4/S5/S6 | 32 | 5.39 | 2.0 × 10⁻⁴ | S3 vs S6 | 32 | 0.84 | 4.0 × 10⁻⁴ | 0.006 | ** |
| SNr | Ch | Accept Good | S1/S2/S3/S4/S5/S6 | 32 | 5.39 | 2.0 × 10⁻⁴ | S4 vs S5 | 32 | 0.12 | 0.473 | 1.000 | n.s. |
| SNr | Ch | Accept Good | S1/S2/S3/S4/S5/S6 | 32 | 5.39 | 2.0 × 10⁻⁴ | S4 vs S6 | 32 | 0.55 | 0.021 | 0.310 | n.s. |
| SNr | Ch | Accept Good | S1/S2/S3/S4/S5/S6 | 32 | 5.39 | 2.0 × 10⁻⁴ | S5 vs S6 | 32 | 0.43 | 0.049 | 0.739 | n.s. |
| SNr | Ch | Accept Bad | S3/S5 | 31 | 5.67 | 0.023 | S3 vs S5 | 31 | −0.47 | 0.023 | 0.023 | * |
| SNr | Ch | Reject Bad | S1/S2/S4/S6 | 32 | 6.35 | 9.0 × 10⁻⁴ | S1 vs S2 | 32 | −0.14 | 0.427 | 1.000 | n.s. |
| SNr | Ch | Reject Bad | S1/S2/S4/S6 | 32 | 6.35 | 9.0 × 10⁻⁴ | S1 vs S4 | 32 | −0.45 | 0.023 | 0.141 | n.s. |
| SNr | Ch | Reject Bad | S1/S2/S4/S6 | 32 | 6.35 | 9.0 × 10⁻⁴ | S1 vs S6 | 32 | 0.28 | 0.137 | 0.825 | n.s. |
| SNr | Ch | Reject Bad | S1/S2/S4/S6 | 32 | 6.35 | 9.0 × 10⁻⁴ | S2 vs S4 | 32 | −0.31 | 0.009 | 0.054 | n.s. |
| SNr | Ch | Reject Bad | S1/S2/S4/S6 | 32 | 6.35 | 9.0 × 10⁻⁴ | S2 vs S6 | 32 | 0.42 | 0.039 | 0.236 | n.s. |
| SNr | Ch | Reject Bad | S1/S2/S4/S6 | 32 | 6.35 | 9.0 × 10⁻⁴ | S4 vs S6 | 32 | 0.74 | 1.0 × 10⁻⁴ | 6.0 × 10⁻⁴ | *** |
| SNr | Cr | Accept Good | S1/S2/S3/S4/S5/S6 | 37 | 3.84 | 0.002 | S1 vs S2 | 37 | 0.14 | 0.393 | 1.000 | n.s. |
| SNr | Cr | Accept Good | S1/S2/S3/S4/S5/S6 | 37 | 3.84 | 0.002 | S1 vs S3 | 37 | 0.36 | 0.068 | 1.000 | n.s. |
| SNr | Cr | Accept Good | S1/S2/S3/S4/S5/S6 | 37 | 3.84 | 0.002 | S1 vs S4 | 37 | 0.58 | 3.0 × 10⁻⁴ | 0.004 | ** |
| SNr | Cr | Accept Good | S1/S2/S3/S4/S5/S6 | 37 | 3.84 | 0.002 | S1 vs S5 | 37 | 0.19 | 0.278 | 1.000 | n.s. |
| SNr | Cr | Accept Good | S1/S2/S3/S4/S5/S6 | 37 | 3.84 | 0.002 | S1 vs S6 | 37 | 0.73 | 0.003 | 0.042 | * |
| SNr | Cr | Accept Good | S1/S2/S3/S4/S5/S6 | 37 | 3.84 | 0.002 | S2 vs S3 | 37 | 0.22 | 0.297 | 1.000 | n.s. |
| SNr | Cr | Accept Good | S1/S2/S3/S4/S5/S6 | 37 | 3.84 | 0.002 | S2 vs S4 | 37 | 0.44 | 0.034 | 0.505 | n.s. |
| SNr | Cr | Accept Good | S1/S2/S3/S4/S5/S6 | 37 | 3.84 | 0.002 | S2 vs S5 | 37 | 0.05 | 0.801 | 1.000 | n.s. |
| SNr | Cr | Accept Good | S1/S2/S3/S4/S5/S6 | 37 | 3.84 | 0.002 | S2 vs S6 | 37 | 0.59 | 0.017 | 0.253 | n.s. |
| SNr | Cr | Accept Good | S1/S2/S3/S4/S5/S6 | 37 | 3.84 | 0.002 | S3 vs S4 | 37 | 0.21 | 0.106 | 1.000 | n.s. |
| SNr | Cr | Accept Good | S1/S2/S3/S4/S5/S6 | 37 | 3.84 | 0.002 | S3 vs S5 | 37 | −0.18 | 0.315 | 1.000 | n.s. |
| SNr | Cr | Accept Good | S1/S2/S3/S4/S5/S6 | 37 | 3.84 | 0.002 | S3 vs S6 | 37 | 0.37 | 0.210 | 1.000 | n.s. |
| SNr | Cr | Accept Good | S1/S2/S3/S4/S5/S6 | 37 | 3.84 | 0.002 | S4 vs S5 | 37 | −0.39 | 0.027 | 0.411 | n.s. |
| SNr | Cr | Accept Good | S1/S2/S3/S4/S5/S6 | 37 | 3.84 | 0.002 | S4 vs S6 | 37 | 0.15 | 0.528 | 1.000 | n.s. |
| SNr | Cr | Accept Good | S1/S2/S3/S4/S5/S6 | 37 | 3.84 | 0.002 | S5 vs S6 | 37 | 0.54 | 0.011 | 0.162 | n.s. |
| SNr | Cr | Accept Bad | S3/S5 | 22 | 0.06 | 0.810 | Overall only |  |  |  |  |  |
| SNr | Cr | Reject Bad | S1/S2/S4/S6 | 36 | 0.38 | 0.766 | Overall only |  |  |  |  |  |
| SNr | Sp | Accept Good | S1/S2/S3/S4/S5/S6 | 22 | 1.12 | 0.350 | Overall only |  |  |  |  |  |
| SNr | Sp | Accept Bad | S1 | 22 | — | — | Overall only |  |  |  |  |  |
| SNr | Sp | Reject Bad | S2/S3/S4/S5/S6 | 22 | 1.05 | 0.383 | Overall only |  |  |  |  |  |
| FEF | Ch | Accept Good | S1/S2/S3/S4/S5/S6 | 33 | 1.21 | 0.309 | Overall only |  |  |  |  |  |
| FEF | Ch | Accept Bad | S3/S5 | 33 | 0.01 | 0.906 | Overall only |  |  |  |  |  |
| FEF | Ch | Reject Bad | S1/S2/S4/S6 | 33 | 2.13 | 0.101 | Overall only |  |  |  |  |  |
| FEF | Cr | Accept Good | S1/S2/S3/S4/S5/S6 | 50 | 1.48 | 0.204 | Overall only |  |  |  |  |  |
| FEF | Cr | Accept Bad | S3/S5 | 49 | 2.39 | 0.126 | Overall only |  |  |  |  |  |
| FEF | Cr | Reject Bad | S1/S2/S4/S6 | 49 | 3.55 | 0.013 | S1 vs S2 | 49 | −0.02 | 0.861 | 1.000 | n.s. |
| FEF | Cr | Reject Bad | S1/S2/S4/S6 | 49 | 3.55 | 0.013 | S1 vs S4 | 49 | −0.04 | 0.676 | 1.000 | n.s. |
| FEF | Cr | Reject Bad | S1/S2/S4/S6 | 49 | 3.55 | 0.013 | S1 vs S6 | 49 | 0.31 | 0.034 | 0.205 | n.s. |
| FEF | Cr | Reject Bad | S1/S2/S4/S6 | 49 | 3.55 | 0.013 | S2 vs S4 | 49 | −0.02 | 0.787 | 1.000 | n.s. |
| FEF | Cr | Reject Bad | S1/S2/S4/S6 | 49 | 3.55 | 0.013 | S2 vs S6 | 49 | 0.33 | 0.021 | 0.126 | n.s. |
| FEF | Cr | Reject Bad | S1/S2/S4/S6 | 49 | 3.55 | 0.013 | S4 vs S6 | 49 | 0.35 | 0.007 | 0.043 | * |
| FEF | Sp | Accept Good | S1/S2/S3/S4/S5/S6 | 16 | 0.41 | 0.842 | Overall only |  |  |  |  |  |
| FEF | Sp | Accept Bad | S1 | 15 | — | — | Overall only |  |  |  |  |  |
| FEF | Sp | Reject Bad | S2/S3/S4/S5/S6 | 16 | 0.91 | 0.469 | Overall only |  |  |  |  |  |
