## Supplemental Table 13 for "Parallel basal ganglia and frontal cortical outputs differentially encode context-dependent evaluation and categorical commitment during choice"

**Table S13. Scene-level permutation statistics for contra forced-choice target-onset panels in Fig. S6.**

Statistics compare scene-specific normalized target-period firing rates for Accept Good responses across forced-choice scenes 7–10, separately for each monkey and recording area.

Test: repeated-measures permutation ANOVA across scenes (10,000 permutations; seed 20260528). When the overall test was significant, pairwise sign-flip permutation tests were performed with Bonferroni correction.

Mean diff is the first minus the second scene for the listed post-hoc contrast. Pairwise tests are shown only for region/monkey combinations with a significant overall permutation ANOVA.

| **Region** | **Monkey** | **n** | **Scenes** | **F** | **Overall P** | **Post-hoc** | **n pairs** | **Mean diff** | **Raw P** | **Adj. P** | **Sig.** |
| --- | --- | --- | --- | --- | --- | --- | --- | --- | --- | --- | --- |
| SNr | Ch | 32 | S7/S8/S9/S10 | 23.17 | 1.0 × 10⁻⁴ | S7 vs S8 | 32 | 0.33 | 0.078 | 0.469 | n.s. |
| SNr | Ch | 32 | S7/S8/S9/S10 | 23.17 | 1.0 × 10⁻⁴ | S7 vs S9 | 32 | 0.01 | 0.946 | 1.000 | n.s. |
| SNr | Ch | 32 | S7/S8/S9/S10 | 23.17 | 1.0 × 10⁻⁴ | S7 vs S10 | 32 | −1.13 | 1.0 × 10⁻⁴ | 6.0 × 10⁻⁴ | *** |
| SNr | Ch | 32 | S7/S8/S9/S10 | 23.17 | 1.0 × 10⁻⁴ | S8 vs S9 | 32 | −0.32 | 0.123 | 0.736 | n.s. |
| SNr | Ch | 32 | S7/S8/S9/S10 | 23.17 | 1.0 × 10⁻⁴ | S8 vs S10 | 32 | −1.45 | 1.0 × 10⁻⁴ | 6.0 × 10⁻⁴ | *** |
| SNr | Ch | 32 | S7/S8/S9/S10 | 23.17 | 1.0 × 10⁻⁴ | S9 vs S10 | 32 | −1.14 | 1.0 × 10⁻⁴ | 6.0 × 10⁻⁴ | *** |
| SNr | Cr | 37 | S7/S8/S9/S10 | 6.64 | 3.0 × 10⁻⁴ | S7 vs S8 | 37 | 0.60 | 0.004 | 0.025 | * |
| SNr | Cr | 37 | S7/S8/S9/S10 | 6.64 | 3.0 × 10⁻⁴ | S7 vs S9 | 37 | −0.14 | 0.617 | 1.000 | n.s. |
| SNr | Cr | 37 | S7/S8/S9/S10 | 6.64 | 3.0 × 10⁻⁴ | S7 vs S10 | 37 | −0.46 | 0.118 | 0.706 | n.s. |
| SNr | Cr | 37 | S7/S8/S9/S10 | 6.64 | 3.0 × 10⁻⁴ | S8 vs S9 | 37 | −0.74 | 0.002 | 0.010 | ** |
| SNr | Cr | 37 | S7/S8/S9/S10 | 6.64 | 3.0 × 10⁻⁴ | S8 vs S10 | 37 | −1.06 | 2.0 × 10⁻⁴ | 0.001 | ** |
| SNr | Cr | 37 | S7/S8/S9/S10 | 6.64 | 3.0 × 10⁻⁴ | S9 vs S10 | 37 | −0.32 | 0.132 | 0.790 | n.s. |
| SNr | Sp | 22 | S7/S8/S9/S10 | 10.17 | 2.0 × 10⁻⁴ | S7 vs S8 | 22 | −0.27 | 0.390 | 1.000 | n.s. |
| SNr | Sp | 22 | S7/S8/S9/S10 | 10.17 | 2.0 × 10⁻⁴ | S7 vs S9 | 22 | −0.12 | 0.724 | 1.000 | n.s. |
| SNr | Sp | 22 | S7/S8/S9/S10 | 10.17 | 2.0 × 10⁻⁴ | S7 vs S10 | 22 | −1.42 | 1.0 × 10⁻⁴ | 6.0 × 10⁻⁴ | *** |
| SNr | Sp | 22 | S7/S8/S9/S10 | 10.17 | 2.0 × 10⁻⁴ | S8 vs S9 | 22 | 0.15 | 0.714 | 1.000 | n.s. |
| SNr | Sp | 22 | S7/S8/S9/S10 | 10.17 | 2.0 × 10⁻⁴ | S8 vs S10 | 22 | −1.15 | 9.0 × 10⁻⁴ | 0.005 | ** |
| SNr | Sp | 22 | S7/S8/S9/S10 | 10.17 | 2.0 × 10⁻⁴ | S9 vs S10 | 22 | −1.30 | 2.0 × 10⁻⁴ | 0.001 | ** |
| FEF | Ch | 33 | S7/S8/S9/S10 | 0.45 | 0.724 | Overall only |  |  |  |  |  |
| FEF | Cr | 50 | S7/S8/S9/S10 | 4.90 | 0.004 | S7 vs S8 | 50 | −0.04 | 0.528 | 1.000 | n.s. |
| FEF | Cr | 50 | S7/S8/S9/S10 | 4.90 | 0.004 | S7 vs S9 | 50 | −0.03 | 0.673 | 1.000 | n.s. |
| FEF | Cr | 50 | S7/S8/S9/S10 | 4.90 | 0.004 | S7 vs S10 | 50 | 0.21 | 0.021 | 0.127 | n.s. |
| FEF | Cr | 50 | S7/S8/S9/S10 | 4.90 | 0.004 | S8 vs S9 | 50 | 0.01 | 0.882 | 1.000 | n.s. |
| FEF | Cr | 50 | S7/S8/S9/S10 | 4.90 | 0.004 | S8 vs S10 | 50 | 0.25 | 0.002 | 0.015 | * |
| FEF | Cr | 50 | S7/S8/S9/S10 | 4.90 | 0.004 | S9 vs S10 | 50 | 0.24 | 9.0 × 10⁻⁴ | 0.005 | ** |
| FEF | Sp | 16 | S7/S8/S9/S10 | 0.16 | 0.924 | Overall only |  |  |  |  |  |
