## Supplemental Table 14 for "Parallel basal ganglia and frontal cortical outputs differentially encode context-dependent evaluation and categorical commitment during choice"

**Table S14. Scene-level permutation statistics for ipsi forced-choice target-onset panels in Fig. S7.**

Statistics compare scene-specific normalized target-period firing rates for Accept Good responses across forced-choice scenes 7–10, separately for each monkey and recording area.

Test: repeated-measures permutation ANOVA across scenes (10,000 permutations; seed 20260528). When the overall test was significant, pairwise sign-flip permutation tests were performed with Bonferroni correction.

Mean diff is the first minus the second scene for the listed post-hoc contrast. Pairwise tests are shown only for region/monkey combinations with a significant overall permutation ANOVA.

| **Region** | **Monkey** | **n** | **Scenes** | **F** | **Overall P** | **Post-hoc** | **n pairs** | **Mean diff** | **Raw P** | **Adj. P** | **Sig.** |
| --- | --- | --- | --- | --- | --- | --- | --- | --- | --- | --- | --- |
| SNr | Ch | 32 | S7/S8/S9/S10 | 7.21 | 6.0 × 10⁻⁴ | S7 vs S8 | 32 | 0.60 | 0.020 | 0.120 | n.s. |
| SNr | Ch | 32 | S7/S8/S9/S10 | 7.21 | 6.0 × 10⁻⁴ | S7 vs S9 | 32 | 0.12 | 0.565 | 1.000 | n.s. |
| SNr | Ch | 32 | S7/S8/S9/S10 | 7.21 | 6.0 × 10⁻⁴ | S7 vs S10 | 32 | −0.36 | 0.057 | 0.341 | n.s. |
| SNr | Ch | 32 | S7/S8/S9/S10 | 7.21 | 6.0 × 10⁻⁴ | S8 vs S9 | 32 | −0.48 | 0.029 | 0.173 | n.s. |
| SNr | Ch | 32 | S7/S8/S9/S10 | 7.21 | 6.0 × 10⁻⁴ | S8 vs S10 | 32 | −0.96 | 5.0 × 10⁻⁴ | 0.003 | ** |
| SNr | Ch | 32 | S7/S8/S9/S10 | 7.21 | 6.0 × 10⁻⁴ | S9 vs S10 | 32 | −0.48 | 0.012 | 0.070 | n.s. |
| SNr | Cr | 37 | S7/S8/S9/S10 | 3.50 | 0.016 | S7 vs S8 | 37 | 0.35 | 0.159 | 0.953 | n.s. |
| SNr | Cr | 37 | S7/S8/S9/S10 | 3.50 | 0.016 | S7 vs S9 | 37 | −0.21 | 0.389 | 1.000 | n.s. |
| SNr | Cr | 37 | S7/S8/S9/S10 | 3.50 | 0.016 | S7 vs S10 | 37 | −0.42 | 0.128 | 0.770 | n.s. |
| SNr | Cr | 37 | S7/S8/S9/S10 | 3.50 | 0.016 | S8 vs S9 | 37 | −0.56 | 0.020 | 0.121 | n.s. |
| SNr | Cr | 37 | S7/S8/S9/S10 | 3.50 | 0.016 | S8 vs S10 | 37 | −0.77 | 0.011 | 0.068 | n.s. |
| SNr | Cr | 37 | S7/S8/S9/S10 | 3.50 | 0.016 | S9 vs S10 | 37 | −0.21 | 0.325 | 1.000 | n.s. |
| SNr | Sp | 22 | S7/S8/S9/S10 | 2.74 | 0.045 | S7 vs S8 | 22 | −0.03 | 0.889 | 1.000 | n.s. |
| SNr | Sp | 22 | S7/S8/S9/S10 | 2.74 | 0.045 | S7 vs S9 | 22 | −0.17 | 0.639 | 1.000 | n.s. |
| SNr | Sp | 22 | S7/S8/S9/S10 | 2.74 | 0.045 | S7 vs S10 | 22 | −0.73 | 0.059 | 0.357 | n.s. |
| SNr | Sp | 22 | S7/S8/S9/S10 | 2.74 | 0.045 | S8 vs S9 | 22 | −0.15 | 0.637 | 1.000 | n.s. |
| SNr | Sp | 22 | S7/S8/S9/S10 | 2.74 | 0.045 | S8 vs S10 | 22 | −0.71 | 0.023 | 0.141 | n.s. |
| SNr | Sp | 22 | S7/S8/S9/S10 | 2.74 | 0.045 | S9 vs S10 | 22 | −0.56 | 0.014 | 0.085 | n.s. |
| FEF | Ch | 33 | S7/S8/S9/S10 | 1.94 | 0.128 | Overall only |  |  |  |  |  |
| FEF | Cr | 50 | S7/S8/S9/S10 | 8.83 | 1.0 × 10⁻⁴ | S7 vs S8 | 50 | 0.10 | 0.213 | 1.000 | n.s. |
| FEF | Cr | 50 | S7/S8/S9/S10 | 8.83 | 1.0 × 10⁻⁴ | S7 vs S9 | 50 | 0.20 | 0.040 | 0.239 | n.s. |
| FEF | Cr | 50 | S7/S8/S9/S10 | 8.83 | 1.0 × 10⁻⁴ | S7 vs S10 | 50 | 0.42 | 1.0 × 10⁻⁴ | 6.0 × 10⁻⁴ | *** |
| FEF | Cr | 50 | S7/S8/S9/S10 | 8.83 | 1.0 × 10⁻⁴ | S8 vs S9 | 50 | 0.10 | 0.241 | 1.000 | n.s. |
| FEF | Cr | 50 | S7/S8/S9/S10 | 8.83 | 1.0 × 10⁻⁴ | S8 vs S10 | 50 | 0.32 | 2.0 × 10⁻⁴ | 0.001 | ** |
| FEF | Cr | 50 | S7/S8/S9/S10 | 8.83 | 1.0 × 10⁻⁴ | S9 vs S10 | 50 | 0.22 | 0.010 | 0.063 | n.s. |
| FEF | Sp | 16 | S7/S8/S9/S10 | 0.94 | 0.426 | Overall only |  |  |  |  |  |
